# Modeling how memory CD8 T cells can elicit post-treatment control of HIV infection

**DOI:** 10.64898/2026.09.01.748582

**Authors:** Bharadwaj Vemparala, Caroline Passaes, Delphine Desjardins, Valérie Monceaux, Julien Lemaitre, Adeline Mélard, Caroline Charre, Maël Gourvès, Nastasia Dimant, Nathalie Dereuddre-Bosquet, Aurélie Barrail-Tran, Hélène Gouget, Céline Guillaume, Francis Relouzat, Olivier Lambotte, Michaela Müller-Trutwin, Christine Rouzioux, Véronique Avettand-Fenoël, Roger Le Grand, Asier Sáez-Cirión, Narendra M Dixit, Jérémie Guedj

## Abstract

While most people living with HIV suffer progressive disease following cessation of antiretroviral therapy, a small fraction elicits lasting post-treatment control. Understanding the mechanisms underlying this control is key to devising effective HIV remission strategies. Although recent studies implicate memory CD8 T cells, how these cells establish lasting viremic control remains unknown. Here, we combine mathematical modeling and analysis of data from SIV-infected non-human primates to elucidate the underlying mechanisms. We recognized that sustained antigenic stimulation leads to heritable epigenetic remodeling of the CD8 T cell pool, impairing memory cell survivability. Antiretroviral therapy rapidly suppresses viremia, thereby arresting antigenic stimulation and preserving memory potential. The greater this preservation is, the better would be the memory recall response following viral rebound post-treatment. Our mathematical model based on this hypothesis predicts that post-treatment control is an alternative steady state to progressive infection, realized by strong memory-driven recall responses. Our model fits longitudinal virological data spanning the pre-, during-, and post-antiretroviral treatment phases of infection, and recapitulates the outcomes of progressive disease and long-term remission realized, the latter predominantly with early treatment initiation. It shows, consistently with data, that memory CD8 T cells could drive post-treatment control independently of the size of the latent reservoir, explaining how such control may be realized more widely than estimated with prevalent hypotheses. Our model further explains the existence of a window of treatment initiation times that maximizes the chances of post-treatment control. Finally, model predictions inform interventions targeting memory CD8 T cells for HIV remission.

**Graphical abstract:** 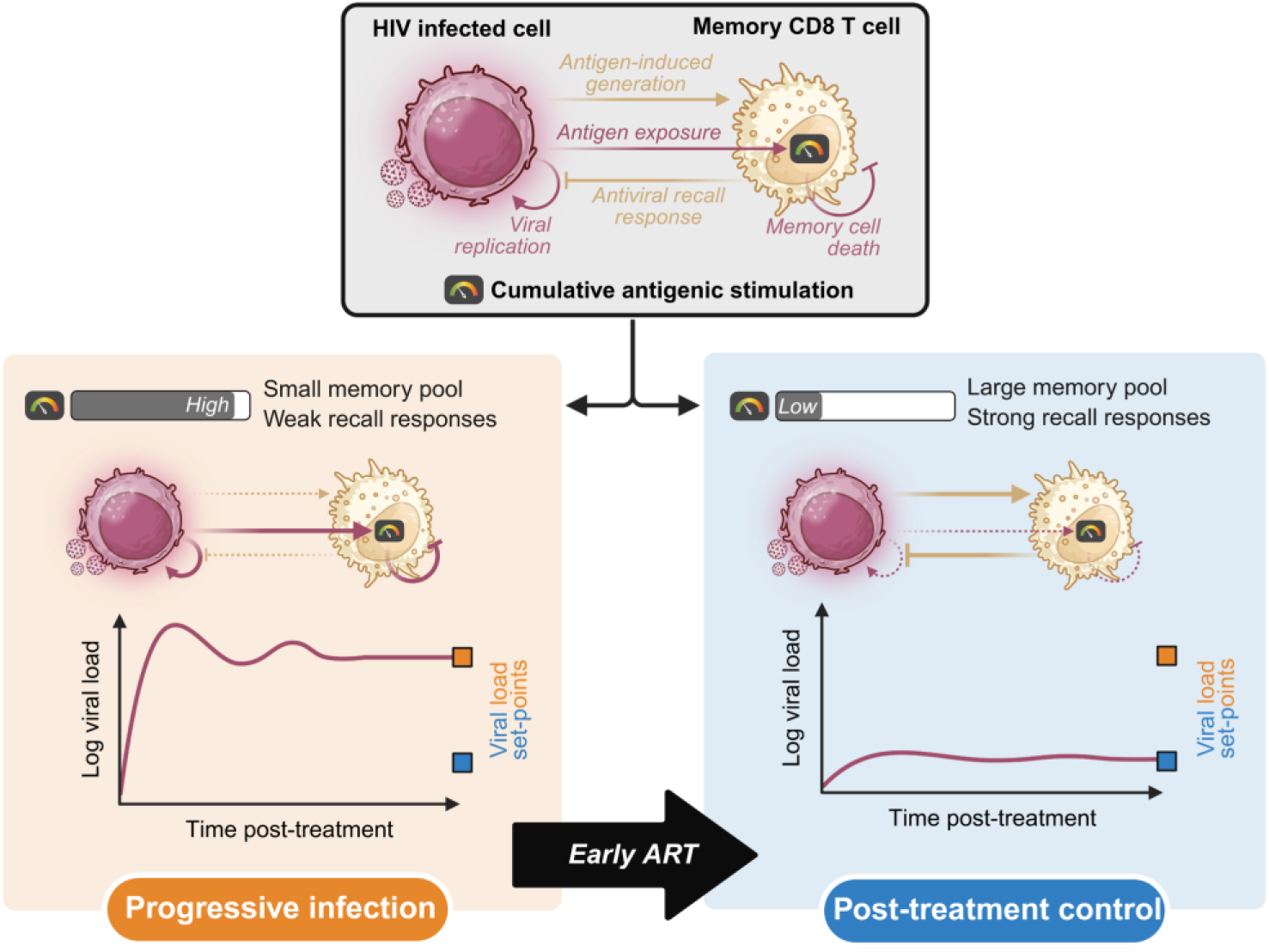

- Mathematical model explains how arresting the loss of memory potential of CD8 T cells with antiretroviral therapy (ART) can trigger post-treatment control of HIV infection
- Post-treatment control is an alternative steady state to progressive infection, and early ART initiation increases the chances of accessing it
- Model predictions recapitulate longitudinal *in vivo* pre-clinical data and describe distinct post-ART outcomes in immunologically similar individuals
- Model presents routes to establish quantitative targets for CD8 T cell–based interventions for sustained HIV remission

## Introduction

Antiretroviral therapy (ART) for people living with HIV (PLWH) is potent at suppressing the viral load but is not curative^1^. For most PLWH, interrupting ART leads to viral rebound and progressive disease; such individuals, termed chronic progressors (CPs), must therefore receive ART lifelong^2^. In contrast, a small fraction of PLWH, called post-treatment controllers (PTCs), exerts long-term viral control after cessation of ART^3^. PTCs offer an important proof of concept that durable viremic control can be achieved without lifelong treatment. Efforts are underway to devise interventions that would elicit such control in CPs^4^. A challenge is that the mechanisms underlying viremic control in PTCs are yet to be established.

Viral rebound post-ART is driven by the stochastic reactivation of latently infected cells, which begin forming early in infection^5^ and can outlast ART^2^. The reservoir of these latently infected cells is central to the prevalent hypothesis underlying post-treatment control. Several human and non-human primate studies have suggested that PTCs have smaller latent reservoirs compared to CPs at treatment interruption^6-10^. The magnitude of the post-treatment viral rebound is typically proportional to the reservoir size^11^. Thus, the rebound is expected to be smaller in PTCs and therefore potentially amenable to control by the immune system. Furthermore, early ART initiation, which substantially improves the chances of post-treatment control^6,12-16^, may do so by arresting viral replication early in infection and thereby limiting the reservoir size. This hypothesis, while plausible, appears to have restricted applicability. Challenging its premise, several studies report similar reservoir sizes in PTCs and CPs^17,18^. Furthermore, only a minority of PLWH harboring small reservoirs achieve post-treatment control^17-19^, and low level of viral replication in pharmacological sanctuary niches may contribute to viral rebound independently of the viral reservoir size^2^. These observations warrant an alternative explanation of post-treatment control.

In an important advance, findings of the recent pVISCONTI study offer insights that carry seeds of an alternative explanation^15^. In this study, SIV-infected macaques were administered ART for two years beginning early (28 days) or late (168 days) post-inoculation and monitored subsequently for their post-treatment status^15^. In agreement with earlier studies^16,20^, a substantially larger proportion of early-treated macaques became PTCs compared to late-treated macaques. Importantly, the control was associated with superior post-ART immune responses driven by polyfunctional memory CD8 T cells. While most early-treated macaques mounted these responses, they were largely missing in late-treated macaques. Notably, the groups had similar reservoir sizes of total SIV DNA in peripheral blood mononuclear cells (PBMCs) at treatment interruption. These findings are consistent with several independent studies that have reported the presence of stem-like CD8 T cells with memory features in HIV and SIV control^21-26^. Furthermore, several recent human studies have reported how a CD8 T cell pool with robust memory-like properties, such as proliferative ability and/or antiviral capacity, is associated with viral control after ART interruption^27^ and/or administration of immunotherapies^28,29^. Thus, strong memory CD8 T cell responses seem to establish sustained post-treatment control independently of the proviral reservoir size in PBMCs.

Several questions arise. First, how do memory CD8 T cell responses elicit post-treatment control? Second, can their effects be quantified, enabling estimation of their minimum strength required for control? Third, how could early treatment initiation lead to strong memory CD8 T cell responses and, thereby, post-treatment control? Fourth, can the maximum delay in treatment initiation beyond which post-treatment control would not be possible be estimated? Addressing these questions would help understand the origins of post-treatment control and inform interventions aimed at eliciting it. Here, we combine mathematical modeling and analysis of longitudinal macaque data to address these questions. To our knowledge, no models to date describe how memory CD8 T cells trigger post-treatment control. Besides, no models have been shown to recapitulate the complex longitudinal datasets comprising virological measurements before, during, and after ART in CPs and PTCs.

We developed a model of HIV dynamics that incorporated the role of memory CD8 T cells. Specifically, we accounted for the generation, survival, and reactivation of memory CD8 T cells in the face of sustained antigenic challenge as well as competition from primary responses. A key phenomenon we incorporated is the gradual loss of memory cell survivability in chronic infection, triggered by heritable epigenetic changes induced by antigenic stimulation^30-35^. This loss impairs recall responses and may limit the chance of post-treatment control. Our model predicted bistability, with one stable steady state characterized by poor recall responses and high viremia, representing progressive disease, and the other by strong recall responses and low viremia, indicating lasting control. While earlier studies too have predicted post-treatment control as an alternative steady state, the origins are distinct and independent of memory responses^36,37^. Our model estimated the minimum strength of the recall responses necessary for driving the system to the control state. We then examined the role of ART. The literature is unclear on how memory cells may arise and help establish post-treatment control in a persistent infection^15,26^. Our formalism, carefully combining virus dynamics with T cell immunology, enabled the construction of a hypothesis to explain this. We reasoned that ART, by halting active viral replication, arrests antigenic stimulation and hence the loss of memory potential. Cessation of ART thus enables memory cells to mount recall responses. If treatment is initiated early, memory potential is better preserved, increasingly mimicking a resolved acute infection. If the resulting recall responses exceed the threshold strength estimated above, post-treatment control would result. Model predictions based on this hypothesis quantitatively recapitulated longitudinal data from the pVISCONTI study. Furthermore, the model predicted that the chance of post-treatment control peaks within a short while after exposure to virus, suggesting that a ‘window of opportunity’ for treatment initiation exists. Finally, we show how the model could be applied to deduce quantitative targets for CD8 T cell–based interventions, under active development for PLWH.

## Results

### Mathematical model of HIV dynamics with memory CD8 T cell responses

We provide an overview of the model here (Fig. 1). The detailed development of the model is in Methods.

**Fig. 1.**
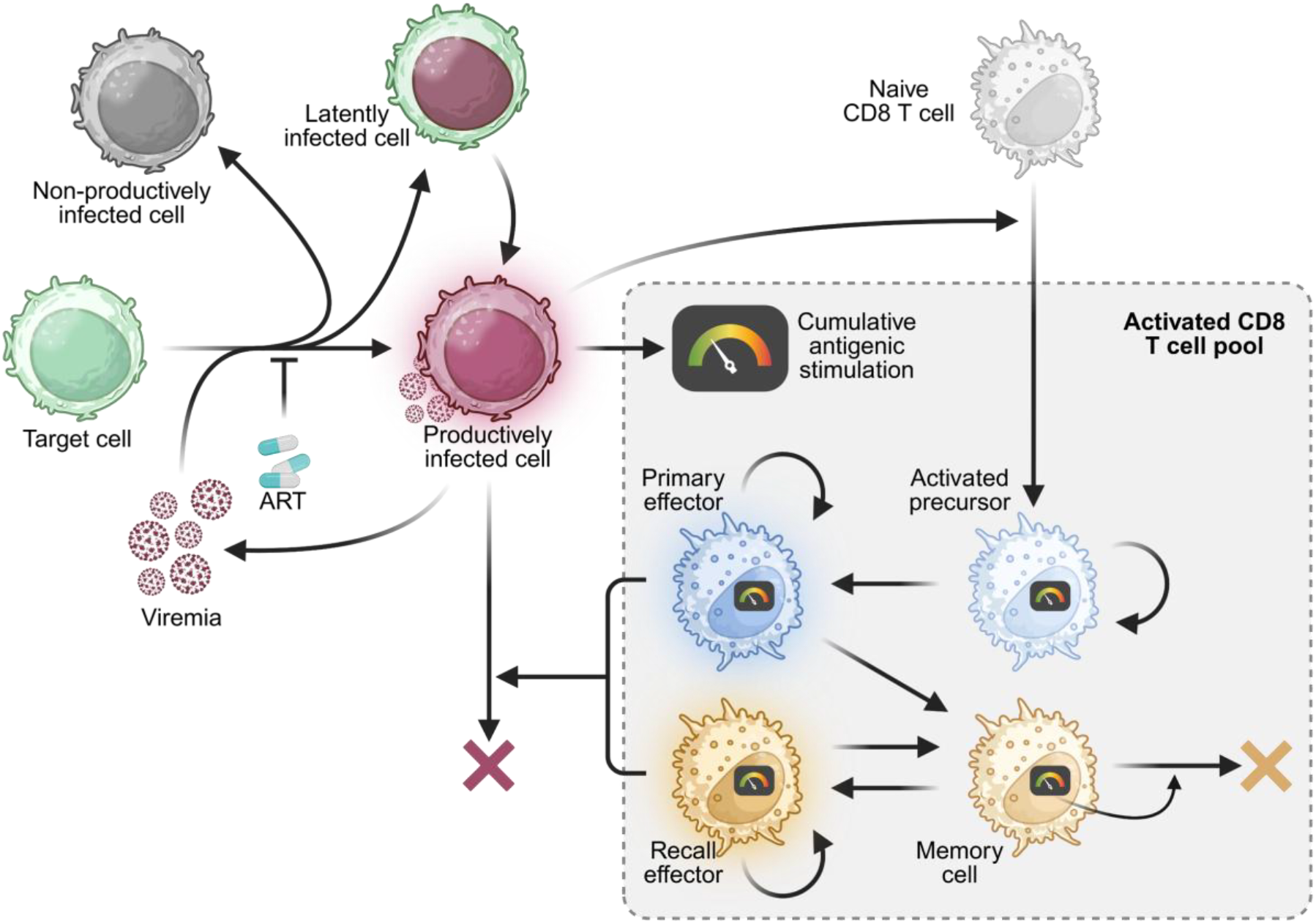
Schematic of the HIV dynamics model with memory CD8 T cells. The various players in our model and their interactions, modeled in equations (1)–(12), are illustrated. Crosses indicate death/loss. Details are in Methods.

Following standard HIV dynamics models^3,36,38,39^, we considered target CD4 T cells, which upon infection by free virions, become either productively infected, non-productively infected, or latently infected. Productively and non-productively infected cells produce free virions and can be killed by virus-induced cytopathicity or by effector CD8 T cells. Free virions produced by non-productively infected cells are replication- and/or infection-incompetent. Latently infected cells have long half-lives and can be reactivated into productively infected cells. Effector CD8 T cells arise in two ways. First, in the presence of antigen, naïve CD8 T cells can be activated into precursor cells, which in turn differentiate into effector CD8 T cells, which we term ‘primary’ effector cells. Most primary effector cells proliferate in an antigen-dependent manner and die. A small fraction of these cells differentiates into memory CD8 T cells. Second, memory cells can differentiate back into effector cells in the presence of antigen, yielding ‘recall’ effector cells. Like primary effectors, most recall effectors proliferate and die while a small fraction differentiates back into memory cells. Activation and proliferation processes of both effectors require interaction with antigen presenting cells. The two effector types thus compete for engagement with antigen presenting cells, in effect inhibiting each other.

An important departure our model makes from previous approaches is in its consideration of ‘cumulative antigenic stimulation’ of CD8 T cells. Epigenetic remodeling following antigenic exposure results in the heritable loss of memory characteristics, via impaired access to stem-associated genes like *TCF7* and *MYB*^32,35,40^ and enhanced access to transcription factors that promote terminal differentiation, like *TOX, NR4A*, and *TBX21*^35,41,42^. This results in the loss of memory cell survivability^30-35^. We therefore developed an age-structured framework that tracks CD8 T cells as they progressively accumulate antigenic stimulation and lets the associated memory cells die at proportionally enhanced rates.

ART blocks new infections of cells, rapidly lowering antigen levels. This arrests the accumulation of antigenic stimulation of CD8 T cells. Viral rebound post-ART resumes this accumulation from the pre-ART level. If the accumulated antigenic stimulation is large pre-ART, then the recall responses post-ART are weak, both due to the smaller surviving memory pool as well as its enhanced death rate. Effector responses may then be largely due to primary effectors, which prove inadequate to control viremia, as with *de novo* infection. In contrast, if the accumulated antigenic stimulation pre-ART is small, swift and strong recall responses emerge from a large memory pool and may exert lasting viremic control post-ART. Early ART initiation likely preserves the memory potential of the CD8 T cell pool by limiting cumulative antigenic stimulation and enables robust recall responses, thereby improving the chances of viral control.

We developed equations to describe the above processes (Methods). The age-structured framework results in partial differential equations, which are difficult to solve and apply for data fitting. We therefore simplified the model by recognizing that most naïve CD8 T cells are recruited into the progenitor pool very early after infection^43^. This allowed us to replace lineage-specific antigenic stimulation with an overall antigenic stimulation applicable to all activated CD8 T cell lineages. The partial differential equations were then transformed into a set of ordinary differential equations (Methods). We solved the latter equations using parameter values representative of SIV infection (Table S1, Fig. S1) and applied them to data fitting. We describe model predictions and the inferences from data fitting next.

### Memory CD8 T cell responses underlie post-treatment control

We first examined whether our model (equations (13)–(22), Methods) could describe post-treatment control driven by memory CD8 T cell responses. Three key quantities at analytic treatment interruption (ATI) are likely to influence the post-treatment outcome: the latent reservoir size (*L*), memory pool size (*M* ^*^), and the level of cumulative antigenic stimulation (*A*^*^). To account for variations in these quantities, we created 5,000 virtual individuals by sampling each of the quantities from reasonably wide distributions (Table S2, Fig. 2) and simulated post-treatment infection dynamics. We found that despite widely differing values of *L, M* ^*^, and *A*^*^ at ATI, the post-treatment viral load converged to one of two set-points separated by ∼4 log_10_ copies mL^-1^ (Fig. 2a, Methods). We defined the state with the higher viremic set-point as the progressive infection state and the state with the lower viremic set-point as the post-treatment control state. Compared with the progressive state, the control state had ∼2-fold superior effector responses (Fig. 2b), driven by a substantially larger memory pool (Fig. 2c) and a lower cumulative antigenic stimulation level (Fig. 2d). Furthermore, the effector response was dominated by recall responses in the control state, whereas the progressive state displayed a predominance of primary responses (Fig. S2a). Notably, the fraction of PTCs was weakly dependent on the size of the latent reservoir (Fig. 2e, Fig. S2b). In contrast, a large memory pool and/or a low level of cumulative antigenic stimulation at ATI led to post-treatment control; progressive infection resulted when memory CD8 T cell responses were compromised (Fig. 2e, Fig. S2c, d). Thus, our model predicts how memory CD8 T cells at ATI, independently of the reservoir size, could determine post-treatment outcome. We next assessed the durability and robustness of the post-treatment control state.

**Fig. 2.**
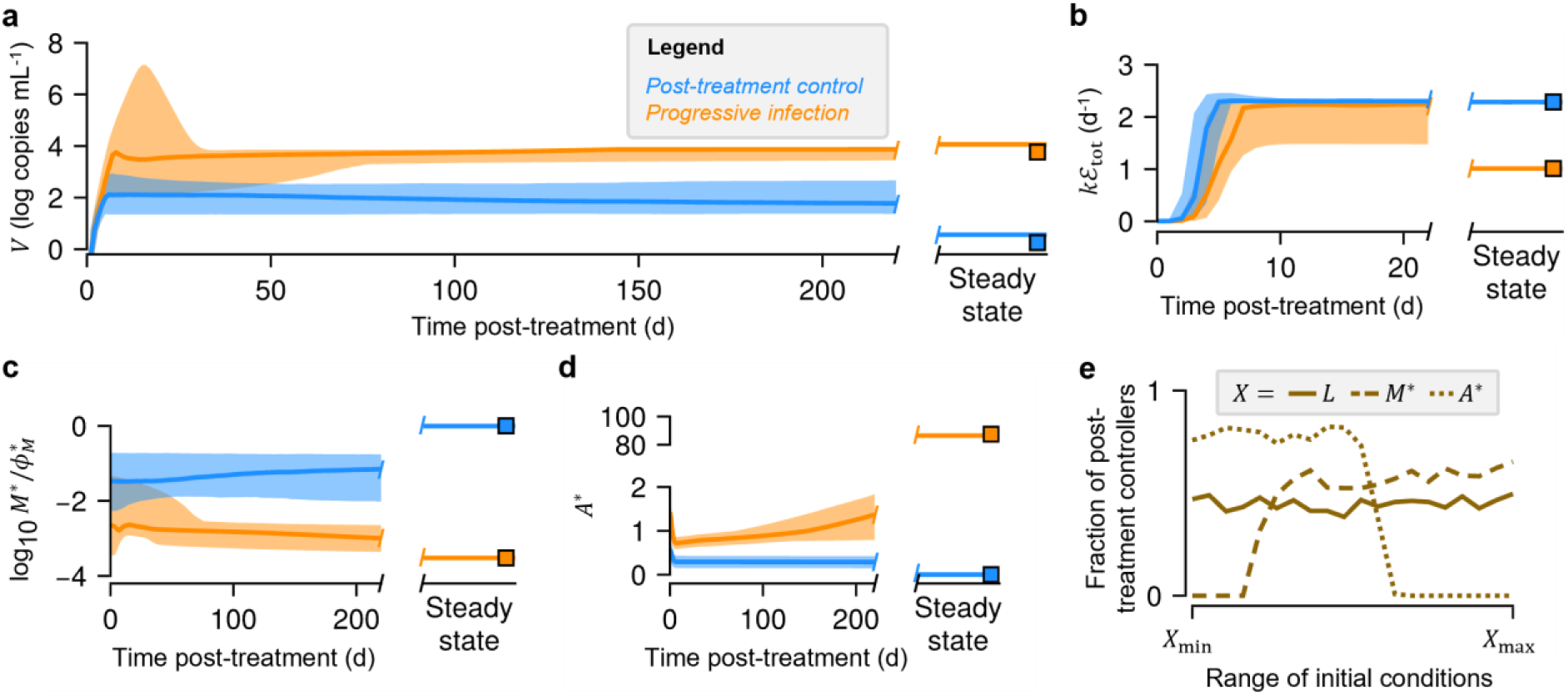
Model describes post-treatment control of HIV infection driven by memory CD8 T cells. Post-treatment time courses predicted for **(a)** viral load, **(b)** total effector responses, **(c)** memory pool size scaled by its carrying capacity 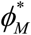, and **(d)** cumulative antigenic stimulation of the CD8 T cell pool for 5,000 virtual individuals obtained by randomly sampling the latent reservoir size, memory pool size, and cumulative antigenic stimulation at treatment interruption (details of the sampled distributions are in Table S2, Fig. S2). Thick lines indicate median values while the colored bands are inter-quartile ranges. Squares are steady state values for the post-treatment control (blue) and progressive infection (orange) states, respectively, estimated by solving the model for stable steady states (Methods). **(e)** Fraction of the simulated samples resulting in post-treatment control when separated by the latent reservoir size (solid line), memory pool size (dashed line), and cumulative antigenic stimulation level (dotted line) at ATI across 20 bins (Table S2).

### Post-treatment control is an alternative stable steady state

Previous modeling studies have argued that post-treatment control is a stable steady state of the system, albeit rarely accessed^36,37^. Steady state would imply that the control would be durable. If such a state were to exist over wide parameter ranges, it would indicate its robustness. We solved our model for its steady states over known ranges of key parameters like the viral infectivity, reactivation rate of latent cells, and proliferation and death rates of effector cells (Fig. 3a, Fig. S3). We found that the model exhibits two stable steady states (bistability) with substantially different viral set-points over wide parameter ranges (Fig. 3a, Fig. S3). In agreement with our simulations above, the low viral load state, or the viremic control state, is associated with robust recall effector responses (Fig. 3b), driven by a large memory pool (Fig. 3c) with minimal cumulative antigenic stimulation (Fig. 3d). In contrast, the high viral load state, or the progressive infection state, is marked by poor effector responses, small memory cell pool sizes and high levels of cumulative antigenic stimulation (Fig. 3a–d). While the magnitudes of the viral load in the two states varied with parameter values, bistability remained a robust feature of the model (Fig. 3a, Fig. S3). (Under extreme parameter settings, the model predicts a third stable steady state with viral load similar to that in progressive infection (Text S4, Fig. S3, S4). This state is associated with dominant recall responses but a small memory cell pool. Note that recall effectors can be sustained by proliferation once they emerge. However, with a small memory cell pool, they are unlikely to emerge^44,45^. The steady state is thus not ordinarily realizable and is therefore not dwelt upon further.)

**Fig. 3.**
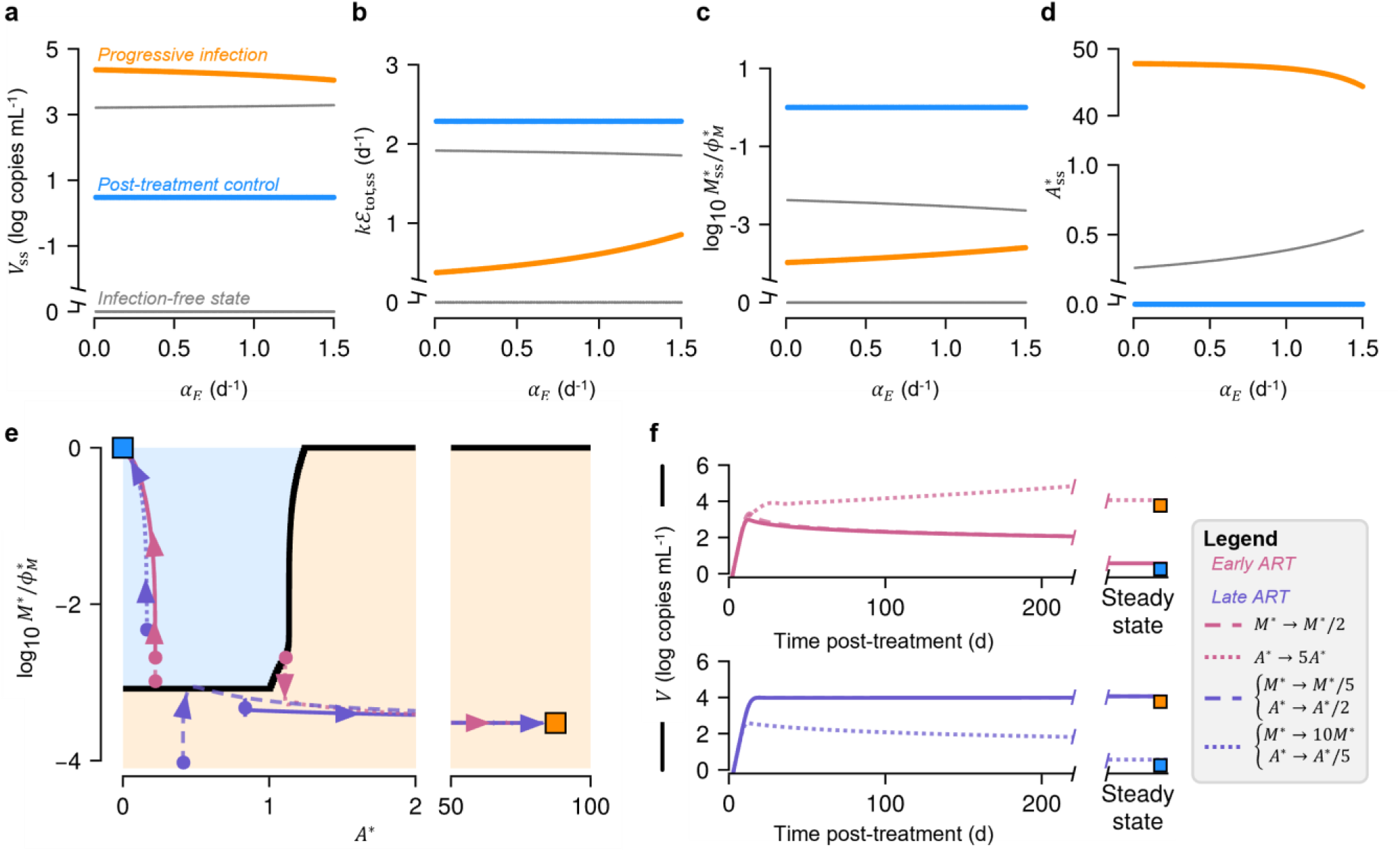
Post-treatment control is an alternative steady state to progressive infection and early ART initiation increases the likelihood of accessing it. **(a–d)** Steady states (denoted with the subscript ss for the variables) of the model estimated by solving it for different values of per-capita proliferation rate of effector cells, *α*_*E*_ (Methods). Bifurcation diagrams for other parameters are presented in Fig. S3. Thin grey branches indicate unstable states while the thick colored branches indicate stable states (blue: post-treatment control, orange: progressive infection). Estimates of the **(a)** steady state viral load, **(b)** total effector responses, **(c)** size of the memory pool scaled by its carrying capacity 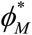, and **(d)** cumulative antigenic stimulation are plotted for different values of *α*_*E*_ . **(e)** Phase portrait of the system, with the phase boundary (black) separating the basins of attraction of the two stable steady states (blue square: post-treatment control; orange square: progressive infection) for a representative set of parameter values (Methods, Table S1). Directed lines indicate post-treatment trajectories of early (pink) and late (slate blue) ART initiation (solid lines), beginning from the correspondingly colored circles. Changing the values of *M* ^*^ and *A*^*^ at treatment interruption (legend), indicated by changes to the location of the colored circles in the phase portrait, may not (dashed lines) or may (dotted lines) switch the post-treatment outcomes. Corresponding viral load trajectories are plotted in **(f)**. Note that the dashed lines overlap with solid lines in **(f)**.

To elucidate conditions leading to the progressive versus the control state, we solved our model equations with different initial sizes of the memory pool, *M* ^*^, and cumulative antigenic stimulation level, *A*^*^, at ATI and predicted the resulting steady state. We found that the *M* ^*^ − *A*^*^ plane split into two regions: high *M* ^*^ and/or low *A*^*^ at ATI resulted in post-treatment control, whereas the opposite conditions led to progressive infection (Fig. 3e). Eliciting post-treatment control would thus require ensuring high *M* ^*^ and/or low *A*^*^ at ATI. Because *A*^*^ builds up with time, early treatment initiation may limit its growth, favoring post-treatment control. We therefore examined next whether the timing of treatment initiation offered a handle to ensure high *M* ^*^ and/or low *A*^*^ at ATI.

### Early ART initiation elicits post-treatment control by preserving memory CD8 T cell potential

We considered an individual, who if untreated, would exhibit progressive infection (Fig. S1). We asked how the outcome would change if the individual were initiated on ART at different times post-infection. Different treatment initiation times would correspond to different initial conditions on the *M* ^*^ − *A*^*^ phase plane, and thus different outcomes. When untreated, the individual is predicted to experience progressive infection, as (*A*^*^, *M* ^*^) = (0, 0), the condition at the onset of infection, falls in the basin of attraction of the progressive infection state (Fig. S1). If treated late (defined as ART initiated 168 days post-infection^15^), the individual is still predicted to experience progressive disease because the condition at ATI falls in the same basin (low *M* ^*^ and high *A*^*^ ; solid line, slate blue, Fig. 3e, f). However, if the individual were to be treated early (ART initiated 28 days post-infection^15^), the condition at ATI falls in the basin of attraction of the post-treatment control state (high *M* ^*^ and low *A*^*^ ; solid line, pink, Fig. 3e, f), leading to viral control. Thus, early treatment initiation elicits post-treatment control by limiting cumulative antigenic stimulation and preserving the size and/or lifespan of the memory cell pool. The outcome is preserved with relatively modest changes (early-treated with *M* ^*^ decreased to 50%; late-treated with *M* ^*^ decreased to 20% and *A*^*^ decreased to 50%) to the condition at ATI (dashed lines, Fig. 3e, f). However, if the changes are sizeable enough to move the condition to the alternative basin (early-treated with *A*^*^ increased 4-fold; late-treated with *M* ^*^ increased 9-fold and *A*^*^ decreased to 20%), the outcome switches (dotted lines, Fig. 3e, f).

In sum, these model predictions describe how the memory pool size and cumulative antigenic stimulation levels at ATI may drive post-treatment outcome, with control likely when ART is initiated early. To assess whether our model describes *in vivo* measurements, we next applied our model to analyze data from SIV-infected macaques.

### Model recapitulates pre-clinical data

We fit our model to longitudinal plasma viral load (SIV-RNA) and total PBMC proviral reservoir size (SIV-DNA) data from infected macaques from the pVISCONTI study^15^ (Methods, Text S5). The study design was as follows (Fig. 4a). Thirty seven cynomolgus macaques that did not carry any SIV-protective MHC haplotypes were infected with SIVmac251^15^. 22 of them underwent ART for two years post-infection (p.i.), with 11 each in the early-treated (initiated 28 days p.i.) and late-treated (initiated 168 days p.i.) groups. These macaques were monitored for ∼1 year after ATI. The remaining 15 macaques were monitored untreated. Macaques with viremic set-point <400 copies mL^-1^ following ATI were termed PTCs. Early ART initiation resulted in a significantly higher proportion of PTCs (9 of 11 with early ART vs. 2 of 11 with late ART).

**Fig. 4.**
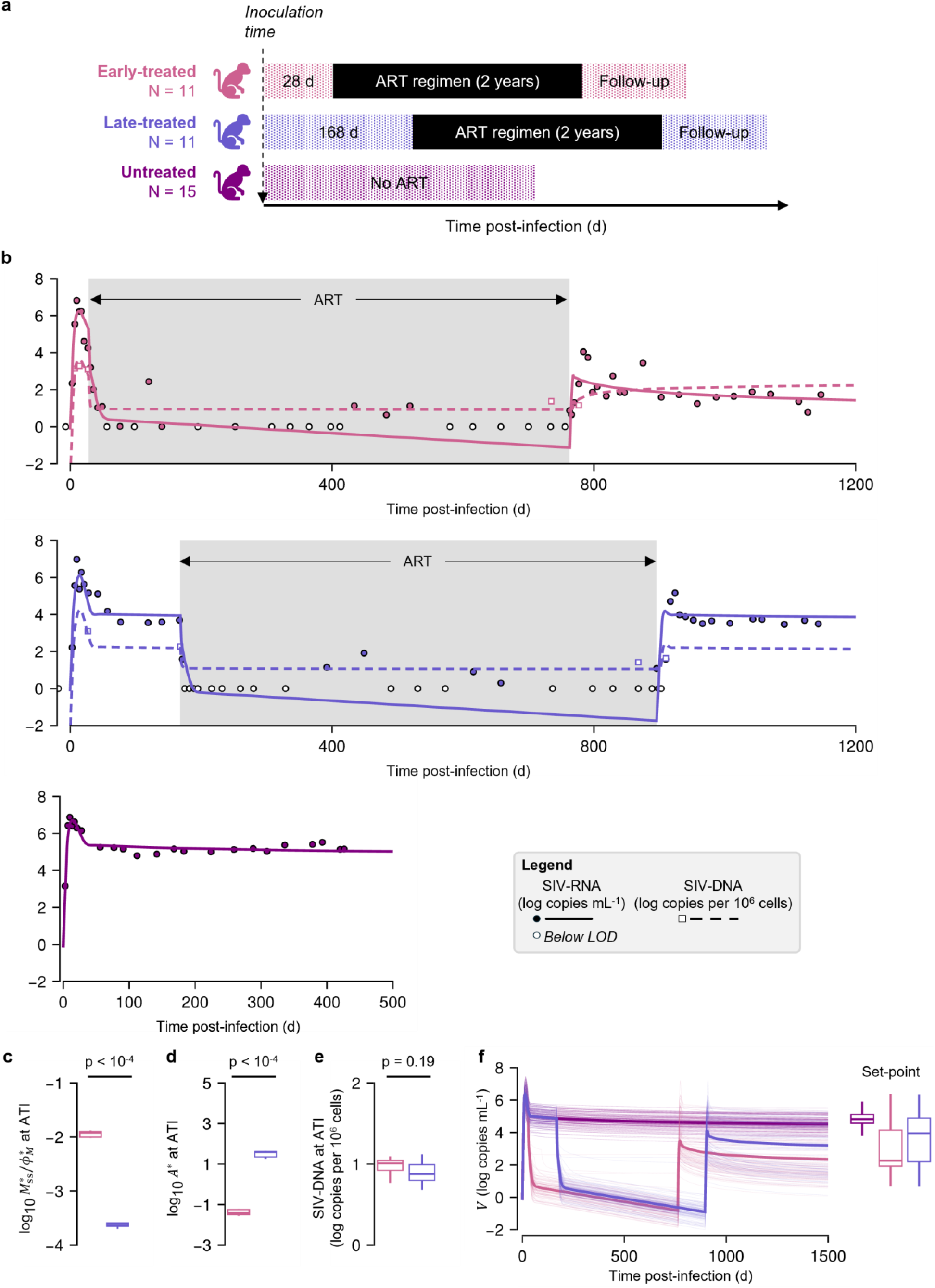
Model recapitulates pre-clinical data and predicts that a superior memory CD8 T cell pool underlies post-treatment control. **(a)** Schematic illustrating the study design with 11 early-treated (pink), 11 late-treated (slate blue), and 15 untreated (purple) macaques^15^. **(b)** Our model fits to the longitudinal viral load (SIV-RNA) and total PBMC proviral reservoir size (SIV-DNA) in one of the early-treated, late-treated, and untreated macaques are presented. Filled circles are viral load measurements and empty circles are observations below the limit of detection (10^0^ copies mL^-1^); squares are reservoir size measurements^15^. The grey-shaded area indicates ART. Macaques with viral load set-point <400 copies mL^-1^ after ART were termed PTCs. Fits to the remaining macaques are presented in Figs. S5 (early-treated), S6 (late-treated), and S7 (untreated). Population parameter estimates are in Table 1, while individual parameter estimates are listed in Table S4. Model-predicted **(c)** size of the memory CD8 T cell pool, **(d)** cumulative antigenic stimulation levels, and **(e)** total SIV-DNA levels in PBMCs at treatment interruption are compared between the groups from the pVISCONTI study. Statistical comparisons were performed using the Mann-Whitney U test. **(f)** Infection dynamics (left) in a virtual population of 10,000 macaques (Methods) constructed using parameter estimates sampled from model fitting and the predicted distribution of set-points (right). Thick lines indicate the median values and thin lines are individual macaques. For convenience, predictions of 200 macaques are plotted.

Our model fit the data well (Fig. 4b, Fig. S5–S7), recapitulating the viral load trajectories in PTCs and CPs. The standard errors for the estimated parameters were small and the visual predictive check (VPC) plots indicated that the variability in predictions across the macaques was well captured (Fig. S8). We estimated eight model parameters (Table 1, Table S4) and found their values to be in agreement with those reported in the literature. For instance, from the parameter *γ*, the burst size was estimated to be ∼5.5×10^4^ virions, which is within the reported range of 4.0×10^4^ and 5.5×10^4^ virions for SIV-infected macaques^46^. The estimate of infectivity, *β*, was ∼10^-8^ mL/d and efficacy of the ART regimen, *ϵ*, was 0.90, respectively, consistent with previous reports^47,48^. The recruitment rate of primary effectors, *ω*_*P*_ *P*^*^ ≈ *ωϕ*_*P*_ was 0.90, close to independent estimates^47^. Sensitivity analysis showed that our predictions were robust to variations in the values of the fixed parameters (Fig. S9, S10, Methods). We also assessed whether existing mathematical models could describe the pVISCONTI data, and found that they failed to distinguish between PTCs and CPs in their fits (Text S6, Fig. S11–S19, Table S5–S7).

**Table 1.** Population parameter estimates of our model. The model was fit (Fig. 4, Fig. S5–S7) in two stages (Methods) with the parameters *β* ^*^, *γ, ψ, ω, α*_*E*_, and *T* (0) estimated in the first stage, and *ϵ* and 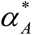 estimated in the second stage, respectively. The remaining parameters of our model were fixed to values from the literature (Table S3). Random effects were removed for *β* ^*^, *ω*, and *T* (0) as they were negligible. RSE: Relative standard error.

| Parameter | Fixed effect (%RSE) | Random effect (%RSE) |
| --- | --- | --- |
| $\log_{10} \beta^*$ (log mL cells<sup>-1</sup> d<sup>-1</sup>) | -4.96 (2.90) | - |
| $\gamma$ (copies cells<sup>-1</sup>) | 1192.78 (21.6) | 0.96 (13.6) |
| $\epsilon$ | 0.90 (2.23) | 0.44 (34.5) |
| $\psi_D$ | 0.91 (43.5) | 0.43 (77.0) |
| $\log_{10} \omega$ (log d<sup>-1</sup>) | 0.26 (38.6) | - |
| $\alpha_E$ (d<sup>-1</sup>) | 1.91 (4.64) | 0.02 (24.6) |
| $\alpha_A^*$ (d<sup>-1</sup>) | 4.8×10<sup>-3</sup> (81.9) | 2.35 (26.2) |
| log<sub>10</sub> $T(0)$ (log cells mL<sup>-1</sup>) | 6.56 (1.97) | - |

In sum, these results suggest that our model accurately describes infection dynamics in individuals with distinct post-ART outcomes governed by memory CD8 T cells. We next examined what mechanisms prior to ATI, predicted by our model, may have resulted in post-treatment control and a higher proportion of PTCs in the early-treated group.

### Superior memory CD8 T cell responses are sufficient to elicit post-treatment control

The best-fit predictions support our mechanistic hypothesis for post-treatment control. The memory CD8 T cell pool was estimated to be ∼50 fold larger (p<10^-4^; Fig. 4c) and the cumulative antigenic stimulation to be ∼10^3^ fold smaller (p<10^-4^; Fig. 4d) in the early-treated group than the late-treated group at ART initiation. The total PBMC proviral reservoir was predicted to be similar between the groups (Fig. 4e), in agreement with experimental observations^15,49^.

To establish the robustness of our predictions, we performed virtual macaque simulations mimicking the pVISCONTI protocols. Using the estimated parameter distributions, we generated a virtual population of 10,000 macaques (Methods) and simulated infection. Our model predicted that most macaques would experience progressive disease if left untreated (viral load set-point >400 copies mL^-1^; Fig. 4f), in line with experimental observations^3^. The same macaques were then subjected in our model to ART for two years (Fig. 4f), starting either early (28 days p.i.) or late (168 days p.i.), following the pVISCONTI study protocol. Our model predicted that a larger fraction (early treatment: median = 72.8%, 95% PI = [36.4%, 100%]; late treatment: median = 45.5%, 95% PI = [18.8%, 72.8%]; PI: prediction interval) of the virtual population elicited post-treatment control with early ART initiation (Fig. 4f). This was due to the cumulative antigenic stimulation being restricted to low levels (Fig. S20a), leading to a larger memory pool (Fig. S20b) with early ART initiation. Again, the total PBMC proviral reservoir was not different between the groups at ATI (Fig. S20c).

Our model thus represents not only a conceptual advance in describing the mechanistic underpinnings of post-treatment control but is also sophisticated enough to recapitulate complex longitudinal *in vivo* datasets, which existing models fail to do. We next applied our model to assess its wider implications for interventions.

### Implications for interventions: ART and beyond

Studies have recognized that a narrow range of ART initiation times, usually referred to as the ‘window of opportunity’, confers a high chance of post-treatment control^50^. To assess whether our model could recapitulate and explain this window of opportunity, we simulated two-year-long ART initiated over a range of post-inoculation times in our virtual population and estimated the post-treatment viral set-point (Fig. 5a). The model predicted that the post-ATI set-point was high if treatment was initiated very early (<5 days post-infection). It then began to decrease, attained a trough with initiation times ∼10 days post-infection, and then rose again to high levels beyond initiation times of ∼150 days post-infection (Fig. 5a). Correspondingly, the fraction of individuals treated that attained post-treatment control started low, peaked with ART initiation times ∼10 days post-infection, and then declined again to low values (Fig. 5b). These estimates are consistent with a recent study, which collated data from multiple experimental studies on SIV-infected macaques and found that the post-treatment set-point viral load was the lowest when ART was initiated ∼20 days post-inoculation^51^. Furthermore, our model offers an explanation of this window. Initiation of ART too early did not allow memory responses to be established in our model, whereas ART initiation too late substantially shrunk the memory CD8 T cell pool (Fig. S21). Both scenarios would lead to progressive disease post-treatment. Intermediate times of ART initiation balance this trade-off, offering a window where superior memory CD8 T cells drive post-treatment control.

**Fig. 5.**
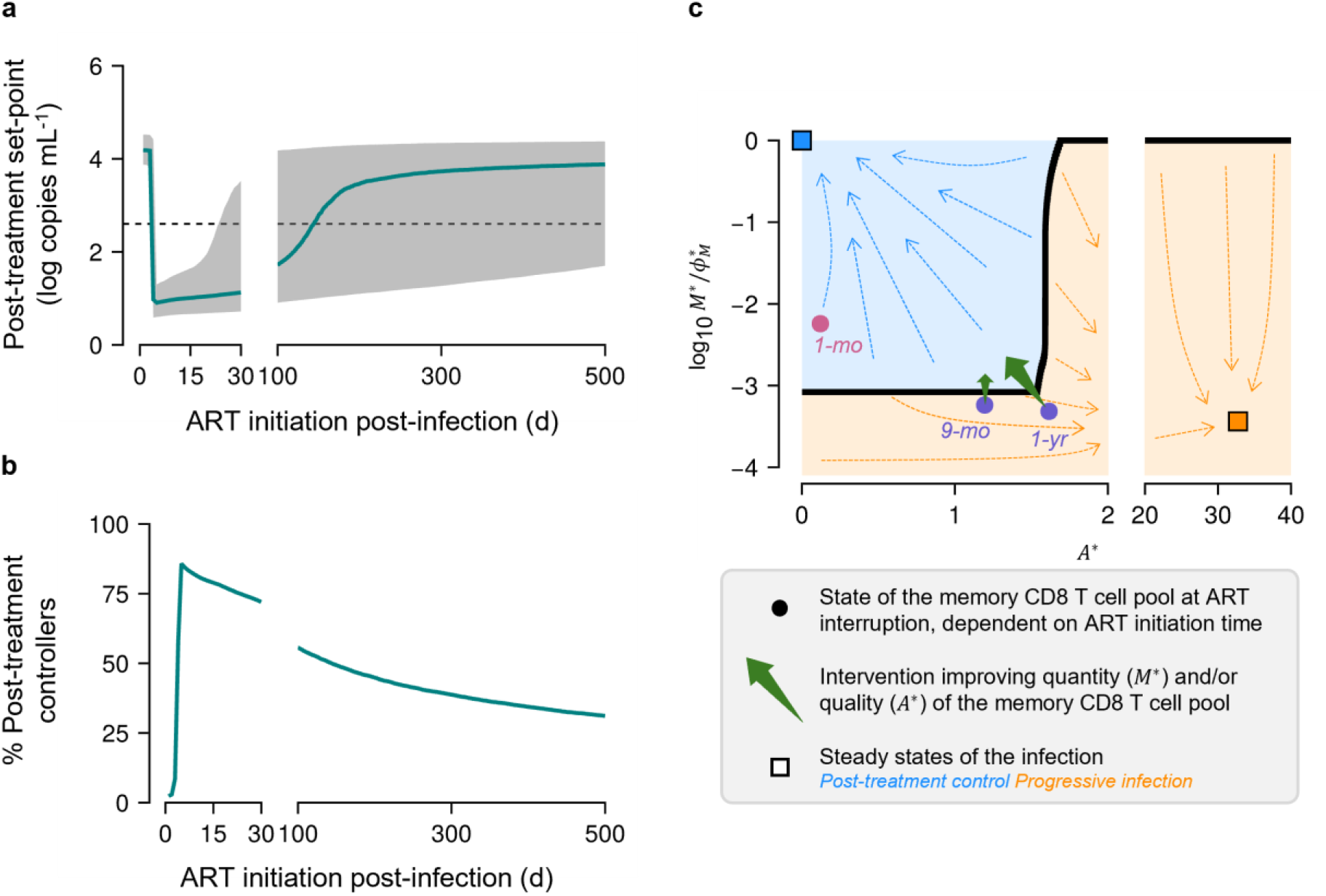
Implications for interventions. **(a, b)** The chance of post-treatment control is maximum for intermediate ART initiation times. **(a)** Post-ART set-point viral load and **(b)** the percentage of PTCs predicted in the virtual population with varying ART initiation time. In **(a)**, the green line indicates the median value, the grey band is the inter-quartile range, and the dashed line is the threshold value of 400 copies mL^-1^ distinguishing PTCs from CPs. **(c)** Model-guided designing of CD8 T cell–based interventions for sustained HIV remission. *M* ^*^ − *A*^*^ landscape predicted by the model (similar to Fig. 4e) showing how the state of the memory CD8 T cell pool at ART interruption, dependent on treatment initiation time (circles), dictates the post-treatment outcome (blue square: post-treatment control; orange square: progressive infection). A phase boundary (black curve) separates the landscape into regions that lead to one of the two outcomes. While early ART (pink circle) leads to post-treatment control, late ART (blue circles) leads to progressive infection. Depending on the state of the CD8 T cell pool at treatment interruption, CD8 T cell–based interventions (green arrows) would need to proportionately decrease *A*^*^ and/or increase *M* ^*^ (indicated by arrow size). Population parameters (Table 1) were used to obtain the landscape.

Our model also predicted that changing the ART duration (varied between 2 months and 5 years, Fig. S22a, b) or increasing the efficacy of the ART regimen (Fig. S22c, d) may negligibly affect these estimates, in accordance with independent experimental observations^16^. Decreasing the ART efficacy, however, which may occur to poor adherence to the drug regimen^1^ or suboptimal penetration of the ART into drug sanctuaries^52^, worsened the chance of post-treatment control (Fig. S22c, d). This is because ineffective ART would fail to stop the antigenic stimulation of CD8 T cells, as has been suggested in independent studies^53^, damaging the memory CD8 T cell pool.

Our model also has implications for interventions beyond ART. Specifically, it can be applied to predict the extent to which the memory pool and/or the cumulative antigenic stimulation should be changed in individuals otherwise destined to progressive infection in order to elicit post-treatment control in them (Fig. 5c).

## Discussion

Achieving durable viral control without continuing ART has important implications. For PLWH, it mitigates adverse effects of long-term drug-induced toxicity, reduces care costs, and improves quality of life^1^. At the population level, estimates suggest that sustained remission is as effective as cure at reducing the risk of transmission^54^. Thus, treatment-free remission through short-term interventions remains one of the key goals of current HIV research^55^. Mechanisms underlying post-treatment control are promising therapeutic targets for achieving this goal. Here, combining mathematical modeling with analysis of *in vivo* pre-clinical data, we predict how memory CD8 T cell responses may drive post-treatment control and how early ART initiation may ensure these responses. Building on recent advances in T cell immunology, we constructed a hypothesis that describes the dynamics of memory CD8 T cells in a persistent infection. Accordingly, exposure to antigen, which induces heritable epigenetic remodeling, results in loss of memory CD8 T cells, a process halted when antigen levels are suppressed with efficacious treatment. Preservation of the memory pool results in robust recall responses upon treatment cessation, which establish lasting viremic control. We developed a mathematical model of HIV dynamics based on this hypothesis and applied it to describe preclinical data. To our knowledge, this is the first study to have fit a model to longitudinal *in vivo* data from all three phases of HIV infection—pre-, during-, and post-ART— and to recapitulate distinct post-treatment outcomes. Furthermore, our model offers a mechanistic explanation of the existence of a window of ART initiation time that maximizes the chances of post-treatment control.

The fate of CD8 T cells is inextricably tied to the fate of an infection^34,56^. In acute-resolved infections, effector CD8 T cells generated in response to the infection die after the pathogen is cleared, leaving behind a functional memory CD8 T cell pool that mounts swift and robust responses upon re-challenge^31,41^. The latter cells are long-lived and thus offer protection from the pathogen for prolonged durations, possibly extending to the lifespan of the individual^57^. On the other hand, in persistent infections, the CD8 T cell pool is largely dysfunctional and memory cells are absent^31,32,35,58^. These seemingly distinct differentiation paths in acute and chronic infection, however, share similarities, especially early in the course of infection. Multiple studies have shown memory cells to be present in the early phases of persistent infections, and, similarly, exhausted cells to be present in the early phases of acute-resolved infections^59-61^. Furthermore, when CD8 T cells isolated early from different infected hosts suffering acute-resolvable or persistent infection, respectively, were adoptively transferred to infected hosts destined to realize the alternative fate, namely, persistent or acute-resolvable infection, the transferred cells switched fates to go down the differentiation pathways appropriate to their new hosts, namely, dysfunction or memory^59-61^. In contrast, CD8 T cells isolated from chronic infections largely failed to acquire any memory features^30,33,60-62^. Thus, the commitment of the CD8 T cell pool to a specific fate is progressive and memory cells can be irreversibly lost as the infection turns chronic. Our model accounts for this progressive loss of memory CD8 T cells by letting their lifespans decrease as cumulative antigenic stimulation grows. Because ART is highly efficacious, early initiation mimics an acute-resolved HIV infection, preserving memory CD8 T cells and eventually improving the chance of post-treatment control. Delaying ART initiation permanently shrinks the memory pool, leading to progressive disease in the absence of ART. This explanation of post-treatment control and the role of early ART initiation based on the progressive dynamics of memory CD8 T cells marks a departure from prevalent hypotheses and therefore an important conceptual advance in our understanding of HIV treatment outcomes.

The prevalent explanation of post-treatment control^4,12,36,63^, which hinges on the latent reservoir, while insightful, appears to have limited applicability. Only a minority of PTCs have smaller reservoirs at treatment interruption than CPs^17-19^. Besides, studies have observed similar proviral reservoir sizes at ATI in PTCs and CPs^17^. Our formalism offers an alternative, potentially more widely applicable explanation of post-treatment control. Smaller reservoirs improving post-treatment control is not inconsistent *per se* with our formalism. When memory CD8 T cell responses are not distinguishing, the reservoir size may become a defining feature. Our model suggests, however, consistently with experiments, that memory CD8 T cell responses tend to be distinct between PTCs and CPs and may dominate differences in the reservoir sizes. It is possible that a strong recall response may suppress rebounding viremia even from a large reservoir and establish control, whereas a weak memory response may fail to suppress rebounding viremia even from a small reservoir and lead to progressive disease. How large a reservoir must be for it to overwhelm the recall responses and determine long-term outcomes remains to be ascertained. Addressing this question may require the application of our model to alternative datasets with much wider differences in the reservoir sizes between PTCs and CPs than seen in the present dataset. Furthermore, future studies may advance our formalism to incorporate reservoir dynamics from a wider set of anatomical regions. Nonetheless, our aim here was to elucidate a mechanism of post-treatment control that is independent of the reservoir size, for which the present dataset, with negligible differences in the PBMC reservoir sizes between the two groups, is well suited.

Independently, several recent human studies also support the idea that memory CD8 T cell responses may be the key driver of long-term HIV control. For example, Crain et al. reported that relative to CPs, PTCs have a larger pool of stem-like CD8 T cells which differentiate into cytolytic effectors^27^. Moreover, the cells were found to target a broader range of viral epitopes and were not restricted to conserved epitopes from mutationally constrained regions^27^. In the same way, Kiani et al.^29^ and Peluso et al.^28^ investigated post-intervention controllers (PICs), i.e., PLWH who exert viral control upon receiving immunomodulatory therapies during ART. Despite administering disparate combinations of multiple broadly neutralizing antibodies (bNAbs) and/or therapeutic vaccines, the studies found robust expansion of stem-like CD8 T cells to underlie the PIC phenotype. Remarkably, all the three studies^27-29^ reported heightened levels of TCF-1 in the CD8 T cell pool of controllers, a canonical signature of pro-memory features^32,56^, also reported in the pVISCONTI study^15^. Consistently, our model predicts more potent memory features in long-term HIV control. We note that the origin of the memory-like CD8 T cells underlying control during the course of the infection is unclear. There is substantial evidence arguing that CD8 T cells in the face of antigenic challenge commit to distinct differentiation fates progressively^32,35,60,61^. Accordingly, in our model, the memory potential of the CD8 T cell pool emerges predominantly in the primary infection phase and is gradually lost as the infection persists. Memory CD8 T cells may also emerge *de novo* on ART^64^ and/or with immunotherapies^28,29^. Future studies that analyze the dynamics of memory-like CD8 T cell responses at the clonal level in controllers are necessary to identify the precise mechanisms at play.

Like our model, existing models of post-treatment control also predict bistability, with the two stable steady states representing progressive disease and control, respectively^36,37^. The origin of the bistability, however, is distinct in these models. It relies on CD8 T cells assuming distinct fates depending on the nature of their stimulation: whereas short-term stimulation leads to their activation and the mounting of effector responses, sustained stimulation results in their exhaustion^65^. Exhaustion in CD8 T cells of PLWH with chronic HIV does build up to high levels^26,62^. Early ART initiation may thus restrict exhaustion and help mount more robust responses post-treatment. The models, however, assume complete reversal of exhaustion following the cessation of antigenic exposure. Thus, a long-enough duration of ART must reset the CD8 T cell responses to the same state as that in primary infection. This implies that the response to the rebounding virus post treatment would be at best as good as the primary response to *de novo* infection. This is inconsistent with the improved responses seen in PTCs post-treatment^15^. While PTCs and CPs mount similar responses pre-treatment, PTCs show substantially better responses post-treatment^3^. Also, exhaustion may not be completely reversible^41,66^. Thus, early treatment initiation may prevent or restrict the extent of irreversible exhaustion, potentially enabling better responses post-treatment. Whether this is adequate to explain the improved responses in PTCs and hence post-treatment control remains to be ascertained.

Models have also been constructed to explain the higher prevalence of sustained HIV remission in pre-clinical studies with other interventions, like passive administration of bNAbs and latency-reversal agents (LRAs), individually as well as in combinations^37,67-70^. For instance, pleiotropic effects of bNAbs, which include clearance of antigen and enhancement of antigen presentation to effector cells, are posited to trigger viral control^37^. Importantly, the enhanced antigen presentation is argued to improve CD8 T cell responses post-treatment beyond that elicited during primary infection. Whether the post-treatment CD8 T cell response elicited by these alternative interventions is similar to what memory CD8 T cells elicit in our model would be interesting to assess.

A key prediction of our model is the existence of a ‘window of opportunity’ for initiating ART that maximizes the chance of post-treatment control. Prior studies have argued that such a window exists^50,51^. Our model offers a plausible mechanistic explanation of the window. Initiating treatment very early precludes the formation of adequate memory CD8 T cells. Late treatment initiation causes the formed memory pool to shrink due to enhanced cumulative antigenic stimulation. Initiating ART in the window maximizes memory responses and hence post-treatment control. We realize that the timing of initiation maximizing the likelihood of post-treatment control, predicted by our model for macaques (∼10 days post infection), differs from those reported in experiments (∼20 days post infection)^51^. This is likely due to some of our model parameters not being estimable and hence fixed based on literature values. Future studies may calibrate the model against additional data that may help estimate all the relevant parameters and improve the accuracy of the predictions, bringing it a step closer to translation to PLWH.

Our model also predicted that the chance of post-treatment control was not sensitive to the ART duration or efficacy (above a threshold). The ART initiation time, and thus the preservation of memory CD8 T cell potential, was the factor deciding the post-treatment outcome. There is evidence in support of this prediction. A recent meta-analysis of all human post-treatment control studies identified that the only factor distinguishing controllers from progressors is the ART initiation time, which was much earlier in controllers^16^. This was echoed in a later study too^20^. We note this agreement with our predictions though with a caveat. Very long ART durations may have effects beyond those considered in our study. For instance, a recent study reported that prolonged ART (>10 years) resulted in rejuvenated CD8 T cell responses due to clonal succession, wherein previously dominant CD8 T cell clones that accumulated signatures of dysfunction were replaced with clones with stemness and robust antiviral properties^64^. While we did not distinguish between multiple CD8 T cell clones in our study, our model could be extended to explicitly account for clones and their interplay to draw insights into the role of ART duration on viral control, another promising direction for future research.

Our study has implications for alternative interventions targeting CD8 T cells for HIV remission. For instance, reprogramming the CD8 T cell pool in CPs to confer memory-like features^71^ is a promising therapeutic strategy currently under development^55^. Our model could be employed to set quantitative targets for such interventions.

More broadly, our model offers a new formalism to study the dynamics of different CD8 T cell subsets—naïve, effector, and memory cells—and their interplay. Although the present context is HIV infection, the formalism could be translated to other settings like acute infections and cancer, informing associated CD8 T cell–based interventions, which remain of great interest for those settings too^41,58,72^.

### Limitations of the study

Our study has limitations. First, our model does not distinguish between the different memory CD8 T cell subsets, namely, central memory, transitional memory, resident memory, and stem cell–like memory cells^31^. Accounting for the relative prevalence of these subsets could provide better predictions of post-treatment control, especially if only some of them drive post-treatment responses. For instance, multiple studies recognize that the levels of lymph node– homing memory cells are highly correlated with viral control^22,23^. Data, however, to accurately describe the dynamics of these memory compartments is currently not available. Second, we neglected the role of other immune players and modalities of responses. Non-cytolytic effects of CD8 T cells, although known to mediate viral control^73^, were not essential for control in the pVISCONTI study^15^. Potent antibody responses by bNAbs as well as autologous antibodies have been reported to exert control post-treatment, especially in the absence of CD8 T cell responses^28,74^. The levels of cytokines like, IFN-α, IFN-γ and IL-10, secreted by other immune cells along with CD8 T cells, were also reported to be correlated with post-treatment control^8,75,76^. Natural killer (NK) cells are understood to govern the transient viral kinetics, especially in lymph nodes, after ART interruption^8,15,77-79^. NK cells may be particularly important in combating viral rebound during the initial hours-days after treatment interruption before the memory-derived CD8 T cell responses can elicit sustained viral control^79,80^. Further studies are necessary to assess the relative importance of these distinct immunological compartments in eliciting post-treatment control. Third, in the pVISCONTI study, differences in exhaustion levels between controllers and progressors were insignificant^15^. Accordingly, we did not explicitly account for exhaustion in our model. Fourth, we considered data of total SIV DNA in PBMCs as a marker of reservoir size as they were more frequently measured and amenable to model fitting. Less frequent measurements have been made of SIV DNA in peripheral lymph nodes and other anatomical sites (the latter only at euthanasia). Differences were observed in the intact SIV DNA in peripheral lymph nodes between PTCs and CPs, the latter showing higher reservoir sizes at treatment interruption^49^. The implications of these differences remain to be ascertained, especially since cell-associated SIV RNA levels, indicative of the reservoir that is transcriptionally active, were similar between the two groups in the peripheral lymph nodes^49^. Interestingly, the differences in the intact proviral reservoir have been attributed to superior CD8 T cell responses in the lymph nodes of PTCs during ART. As discussed above, future studies may report more frequent measurements of these markers, enabling advancement of our formalism to incorporate wider reservoir dynamics and associated immune responses. Finally, we did not account for the evolution of the virus off ART, as it poses a significant risk of the emergence of CD8 T cell escape variants^81^. However, unlike in natural controllers^82-84^, post-treatment controllers appear to exert sustained viral control despite evidence of viral evolution and superinfection^4^. More long-term studies that simultaneously sequence the virus and CD8 T cell clones are necessary to quantify the impact of viral evolution on the durability of post-treatment control.

In summary, we developed a framework that describes how memory CD8 T cell responses underlie post-treatment HIV control. Models that recapitulate the complex longitudinal datasets comprising virological measurements before, during, and after ART in CPs and PTCs were missing so far. These model predictions have the capacity to inform future interventions targeting memory CD8 T cells for HIV remission.

## Methods

### Mathematical model of HIV dynamics with memory CD8 T cells

We constructed an age-structured model of within-host HIV dynamics that integrates the roles of the different subtypes of virus-specific CD8 T cells, including memory cells (Fig. 1). The following equations (1)–(12) comprise the model.

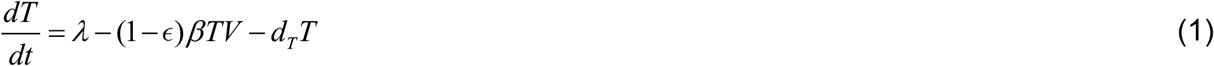

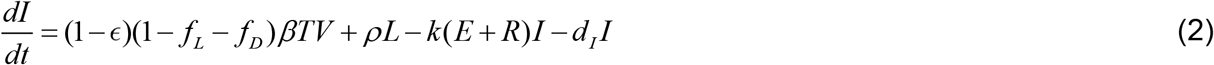

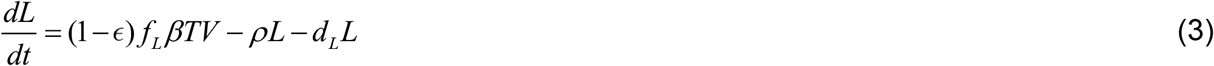

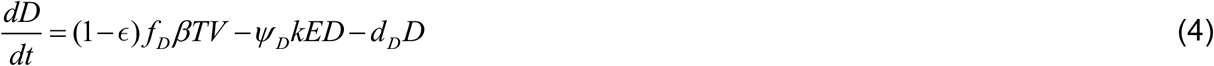

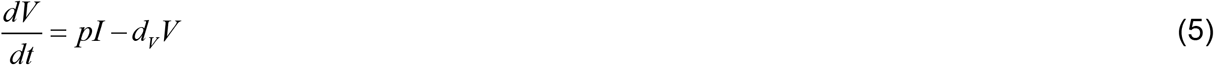

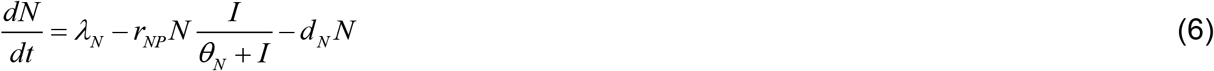

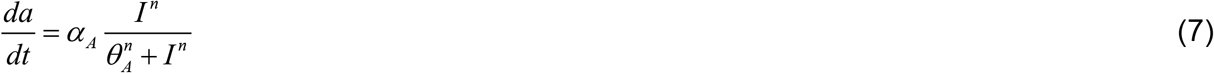

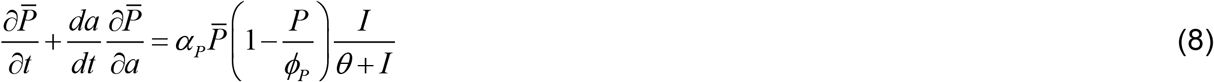

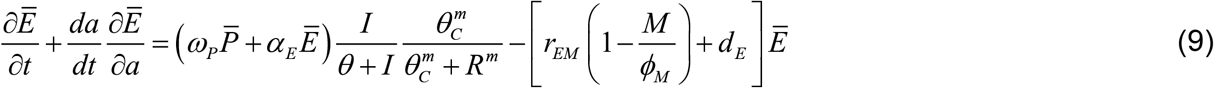

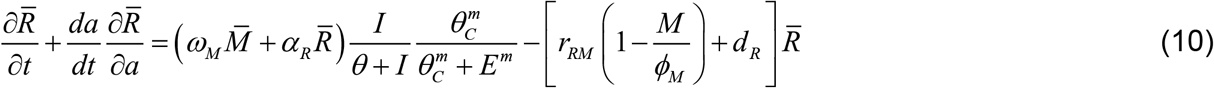

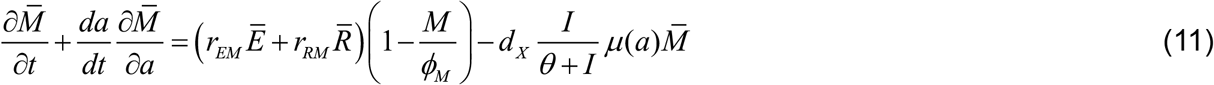

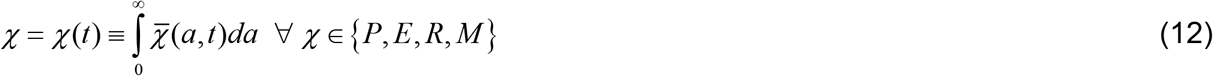

Here, uninfected CD4 T cells, *T*, are recruited at the rate *λ* and die at the rate *d*_*T*_*T* (Eq. (1)). Virus particles, *V*, infect these cells at the maximal rate *βTV* . The infectivity, *β*, is reduced to (1− *ϵ*)*β* under an ART regimen of efficacy *ϵ*. (A combination of reverse transcriptase and integrase strand transfer inhibitors, which inhibit the infection of target cells, comprised the ART regimen in the pVISCONTI study^1,15^. The model could be modified for other classes of antiretrovirals, like protease inhibitors, which hamper viral production from infected cells^85^.) Fractions *f*_*L*_ and *f*_*D*_ of the infection events result in latently and non-productively infected cells, denoted *L* (Eq. (3)) and *D* (Eq. (4)), respectively. The remaining fraction yields productively infected cells, *I* (Eq. (2))^3^. The natural lifespans of these three cell types are 1/ *d*_*L*_, 1/ *d*_*D*_, and 1/ *d*_*I*_, respectively. Productively infected cells are eliminated by the cytolytic activity of virus-specific effector CD8 T cells, with naïve-derived effectors denoted *E* and memory-derived effectors denoted *R*, respectively, with *k* the killing rate constant. ℰ_tot_ = *E* + *R* denotes the total effector cell concentration, and *k*ℰ_tot_ the total effector response. By keeping the killing rate *k* the same for both cell types, we assume that they have the same per cell killing efficiency; stronger memory responses would thus be due to greater expansion of recall cells. Latently infected cells are reactivated at rate *ρ*, following which they become productive. Non-productively infected cells are killed by effector cells. This killing efficiency may depend on the half-life of the defective proviruses and the specific epitopes they present^86,87^. We let the killing rate constant differ from that of productively infected cells by the factor *ψ*_*D*_ . Further, dominant CD8 T cell responses, which form the majority of memory cells, are expected to be driven by intact proviruses. We therefore neglected recall responses against defective proviruses^86^. Productively infected cells produce virions at the rate *pI*, which are cleared at the rate *d*_*V*_*V* (Eq. (5)).

Naïve CD8 T cells, *N*, are produced at the constant rate *λ*_*N*_ and have a lifespan 1/ *d*_*N*_ (Eq. (6)). During an infection, naïve cells are stimulated by antigen and become activated precursors, 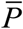. (The overbars represent densities; see below.) These precursors give rise to the downstream precursor-derived primary effectors, 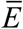, which differentiate into memory cells, 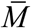. The latter cells produce recall effectors, 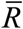, which in turn can differentiate back to 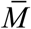 . All of the cell types, beginning with the precursors, are activated. They experience antigenic stimulation by engaging their T cell receptors with viral peptide–loaded MHC complexes on host cells^88^. This antigenic stimulation is assumed to be accumulated irreversibly due to epigenetic remodeling. We quantify the level of cumulative antigenic stimulation as *a* (Eq. (7)), which increases in a saturable manner with the antigen level, the latter determined by *I* . Accumulation begins when a cell is first activated from its naïve state and is carried forward through its lineage as it proliferates and differentiates into different subtypes^32,35,40^. We define 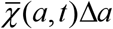 as the concentration (or number density) of activated CD8 T cells of subtype *χ*, where *χ* ∈{*P, E, R, M* }, with their cumulative antigenic stimulation level in the range [*a, a* + Δ*a*) at time *t* during the course of the infection. We construct partial differential equations describing the dynamics of the four activated CD8 T cell subtype levels (Eq. (8)–(11)) with *a* and *t* the independent variables, which we describe below. 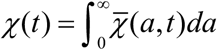 is the total concentration (or number) of the CD8 T cells of the subtype *χ* at time *t* (Eq. (12)).

Naïve cells are activated at the maximal rate *r*_*NP*_, with half-maximal saturation constant *θ*_*N*_, and become activated precursors 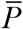 (Eq. (8)). The newly formed precursors have yet to accumulate antigenic experience and thus enter the governing equation for 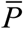 as a boundary condition. Proliferation of precursors is modeled using a logistic equation, with *α*_*P*_ and *ϕ*_*P*_ their growth rate and carrying capacity, respectively. Given their self-renewal capacity, we neglect the death of precursors as an approximation. The logistic form allows recruitment of primary effector cells from the precursor pool at a constant rate, in line with previous models^36,47,89^. This proliferation is regulated by the antigen load, with *θ* being the half-maximal saturation constant. Primary effectors, 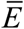, are produced at a rate proportional to the size of the precursor pool, 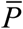, where *ω*_*P*_ is the proportionality constant, and proliferate at the per-capita rate *α*_*E*_ (Eq. (9)). The production and proliferation processes are governed by the level of antigen accessible for stimulation from antigen-presenting cells (APCs)^88^. Primary and recall effectors compete for access to APCs (Text S1, Fig. S23). Here, *θ*_*C*_ is the half-maximal saturation constant and *m* is the Hill coefficient that govern this process. For high relative populations of recall effectors, i.e., *R* ≫ *θ*_*C*_, few APCs remain to engage with *E*, ≫ *θ*_*C*_ limiting their production and proliferation. Similarly, *E* leads to low rates of production and proliferation of *R* . We derived an expression to quantify the impact of this competition on both cell types (Text S1, Fig. S23). Primary effectors have a lifespan 1/ *d*_*E*_ . They differentiate and seed the memory cell pool, 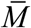, at the maximal rate *r*_*EM*_ . *ϕ*_*M*_ is the upper limit on the memory cell levels. Recall effectors, 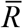, are produced from 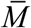 at the per capita rate *ω*_*M*_ (Eq. (10)). Their per-capita proliferation rate is *α*_*R*_, lifespan is 1/ *d*_*R*_, and maximal differentiation rate to memory cells is *r*_*RM*_ . Death of memory cells is governed by both the environment and the antigenic stimulation accumulated (Eq. (11)). An inflammatory milieu is understood to trigger attrition of the memory pool, regulated by transcription factors like T-bet and Blimp-1^41,90-94^. In persistent infections, memory cells are progressively lost due to epigenetic remodeling of the CD8 T cell pool by continuous antigenic stimulation, driven by transcription factors like TOX, triggering irreversible loss of accessibility to pro-memory and pro-survival genes^35,41,72,95^. We account for both these processes in our model by letting memory cells be lost at the rate 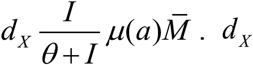 is the maximal death rate of memory cells. The term *I* / (*θ* + *I*) accounts for the influence of the inflammatory environment, with antigen levels, *I*, reflecting the inflammation level. The influence of cumulative antigenic stimulation is captured by the function *μ*(*a*) .

While the partial differential equations above can be solved with suitable initial and boundary conditions, they are not readily amenable to data fitting. We therefore transformed them into a set of ordinary differential equations as follows. Most virus-specific naïve clones are activated immediately after primary infection. The production rate of fresh naïve cells is substantially lower compared to the proliferation rates of the other, activated CD8 T cell subtypes^43^. We exploited this feature in equations (8)–(12), transforming them into ordinary differential equations and yielding an equation for population-averaged cumulative antigenic stimulation, denoted *A* (equations (13)–(22), Methods). All analyses in this study were conducted using these transformed equations. For a representative set of parameter values (Table S1), the model predictions were in agreement with the typical dynamics reported in the literature (Fig. S1).

### Model with population-averaged cumulative antigenic stimulation

Here, we describe how we transformed the model equations (8)–(11) to a set of ordinary differential equations with an equation for population-averaged cumulative antigenic stimulation, denoted *A* . These transformed equations have been employed for analyses throughout this study.

We recognized that most virus-specific naïve clones are activated soon after infection; the pool is not easily replenished as the production rate of naïve cells is low compared to the proliferation rates of the other, activated CD8 T cell subtypes^43^. We therefore assumed that a vast majority of the naïve CD8 T cells enter the activated pool in a short time window immediately after inoculation. Subsequently, only a relatively small fraction enters, limited by the low production rate of naïve cells. We therefore let all activated CD8 T cell lineages carry the same antigenic stimulation at any given time, set by the level of the majority that entered early and corrected to account for the trickle that enters subsequently. We define this population average stimulation level as 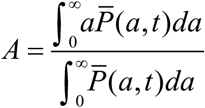. Incorporating this assumption into the equations (11)–(14) transformed them into ordinary differential equations for *P, E, R*, and *M* in *t*, and yielded an equation for *A* (Text S2). The resulting model equations are Equations (S34)–(S44) in Text S2. To solve them, we made a few additional simplifications. Viral production and clearance happen at much faster rates than the other processes in our model^96,97^. So, following earlier studies, we assumed a quasi–steady state between viral production and clearance rates^89^, which implied *pI* ≈ *d*_*V*_*V*, and hence *V* = *γ I* with *γ* = *p* / *d*_*V*_ . We also non-dimensionalized/rescaled our equations using 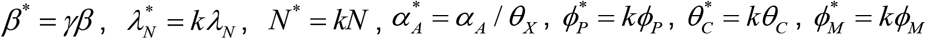, and *χ*^*^ = *k χ* ∀ *χ* ∈{*P, E, R, M* } . This yielded the set of equations below, using which we performed all the analyses in our study:

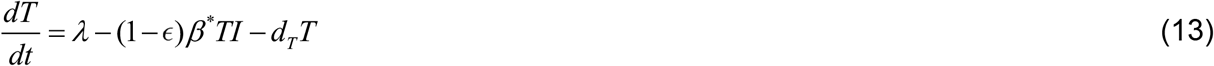

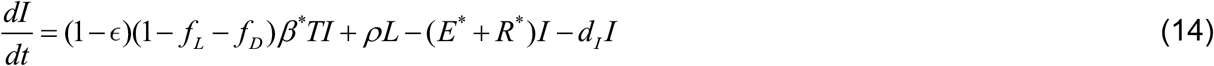

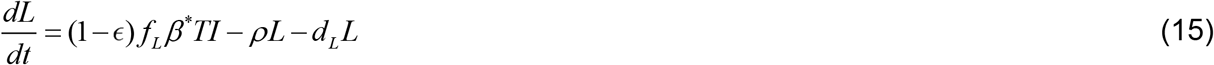

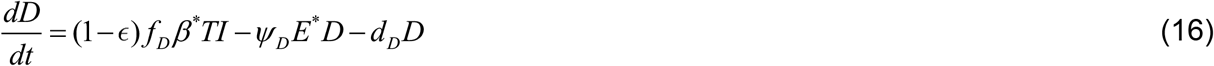

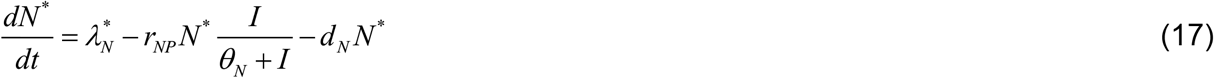

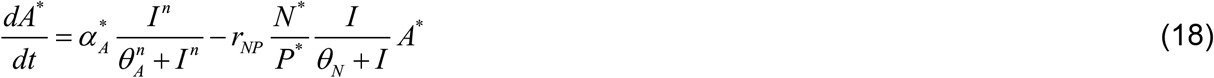

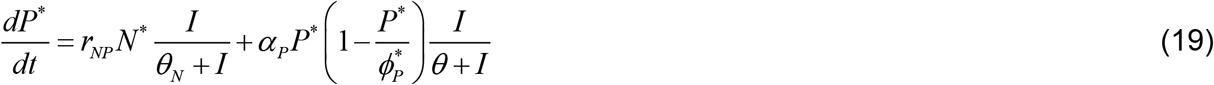

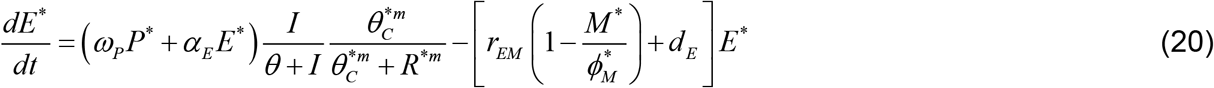

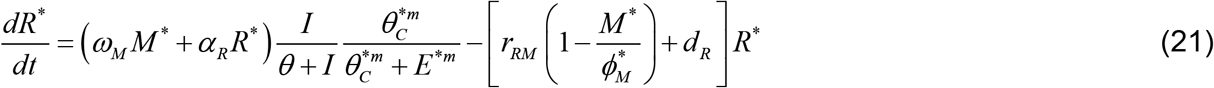

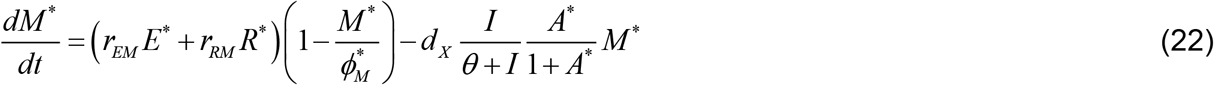

### Steady states and linear stability analysis

We estimated the steady states of the model and identified their nature (stable vs. unstable) using the following procedure (see Text S3). First, as all derivatives vanish at steady state, we equated all equations (13)–(22) in our model to zero. Then, we reduced the dimensionality of the system by writing as many variables as possible as functions of the remaining variables. This yielded a 4-dimensional system with equations for *I, P*^*^, *E*^*^, and *R*^*^ . Finally, we noted that in all but the infection-free state, 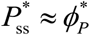. Incorporating this into our system further reduced the dimensionality to 3. Steady states were estimated by solving this 3-dimensional nonlinear system; equations are provided in Text S3. For any given set of parameter values, we estimated the steady states using NSolve of Wolfram Mathematica v12.2. The stability of those states was identified by calculating the eigenvalues of the Jacobian matrix. If the real parts of all the eigenvalues for a state are negative, then it is stable. Otherwise, it is unstable.

### Model fitting to data

We fit the model in two stages. In the first stage, we fit the part of the model described above in equations (13)–(22) without latently infected cells, memory cells, and recall responses to pre-ART data of the three groups. The corresponding equations are listed in Text S5. In this stage, we estimated *β* ^*^, *γ, ψ*_*D*_, *ω, α*_*E*_, and *T* (0) . Then, in the second stage, we fit our full model to complete data (pre-, during-, and post-ART phases of the three groups) while fixing the fixed and random effects of the six parameters above to their respective values estimated in the first stage. In the second stage, we additionally estimated *ϵ* and 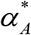, yielding eight parameter estimates in total. The two different effector cells, *E*^*^ and *R*^*^, were assumed to have the same intrinsic properties. Thus, in our model, we fixed *ω*_*P*_ = *ω*_*M*_ = *ω, α*_*E*_ = *α*_*R*_, *r*_*EM*_ = *r*_*RM*_, and *d*_*E*_ = *d*_*R*_ . A similar procedure was employed to fit other, extant models describing post-treatment control (see Results), detailed in Text S6.

At each stage, estimable parameters were identified using the differential-algebraic elimination method for structural identifiability analysis implemented in the Julia package StructuralIdentifiability.jl^98^. Models were fit using the nonlinear mixed effects (NLME) approach implemented in Monolix 2024R1 (https://lixoft.com/), employing the stochastic approximation of the expectation-maximization (SAEM) algorithm. The statistical model linking the viral load (SIV-RNA) and total PBMC proviral reservoir (SIV-DNA) measurements to the model variables is given below.

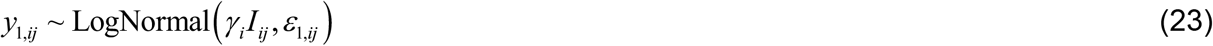

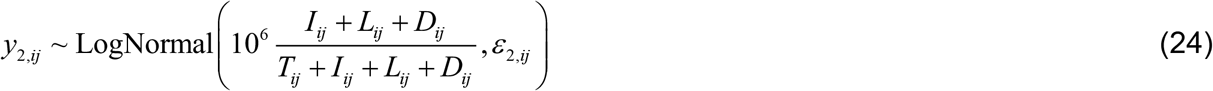

Here, *y*_1,*ij*_ and *y*_2,*ij*_ denote SIV-RNA and SIV-DNA measurements for the individual *i* at the time point *j* . SIV-RNA is the viral RNA copies per mL^-1^, which is *V* = *γ I* . SIV-DNA is the total number of proviral DNA copies (*I* + *L* + *D*) per million CD4 T cells (*T* + *I* + *L* + *D*) in the blood. *ε*_1_ and *ε*_2_ are the residual Gaussian errors. Values of fixed parameters and initial conditions are provided in Table S3. Population parameter estimates are provided in Table 1, and individual estimates in Table S4. Prior to the infection, there are target cells, *T* (0) = *λ* / *d*_*T*_, and naïve CD8 T cells, 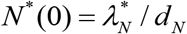, and infection is established by a viral inoculum of size *V* (0) . Given that *P*^*^ is in the denominator of equation (21), to avoid numerical integration errors, *P*^*^ starts with a value in our simulations very close to but not equal to zero. All other variables start from zero.

### Virtual population

In the NLME description of our model, the parameter for ART efficacy, *ϵ*, followed a logit-normal distribution ranging between 0 and 1. All other estimable parameters followed a log-normal distribution to ensure non-negativity. Consequently, log_10_ *β* ^*^, log_10_ *ω*_*P*_, and log_10_ *T* (0) are normally distributed (Table 1). We reconstructed these parameter distributions using values estimated from fitting our model to the data and randomly drew 10,000 samples from these distributions. Each set of these parameter draws represents a virtual macaque.

## Sensitivity analysis

We analyzed the sensitivity of set-point viral load to changes in model parameters using the eFAST routine implemented in GlobalSensitivity.jl package for Julia v1.7.3^99^.

## Data availability

All the raw data files, Monolix codes used for model fitting, and scripts written in Julia v1.7.3 and Wolfram Mathematica v12.2 for analysis and plotting are available on the GitHub repository https://github.com/vembha/SIV_pVISCONTI/.

## Acknowledgements

We thank Jessica M. Conway, Rustom Antia, and Rajat Desikan for comments. This study was supported by IFCPAR/CEFIPRA Project #64T4-2 (A.S.-C., N.M.D, J.G.).

