## Supplementary information for "Modeling how memory CD8 T cells can elicit post-treatment control of HIV infection"

**Details:**

Supplementary information: Text notes: 6, Figures: 23, Tables: 7, References: 20

### Text S1: Competition between primary and recall effector CD8 T cells

In this text note, we derive equations (12) and (13) presented in the main text.

Following the notation used in the main text, the densities of naïve-derived 'primary' effectors and memory-derived 'recall' effector cells are denoted $\bar{E}$ and $\bar{R}$, respectively. Their dynamical equations are:

$$\frac{d\bar{E}}{dt} \equiv \frac{\partial \bar{E}}{\partial t} + \frac{da}{dt}\frac{\partial \bar{E}}{\partial a} = \Gamma_{\bar{E}} - \Lambda_{\bar{E}} \qquad \text{(S1)}$$

$$\frac{d\bar{R}}{dt} \equiv \frac{\partial \bar{R}}{\partial t} + \frac{da}{dt}\frac{\partial \bar{R}}{\partial a} = \Gamma_{\bar{R}} - \Lambda_{\bar{R}} \qquad \text{(S2)}$$

Here, $\Gamma_i$ and $\Lambda_i$ denote the processes for gain and loss of $i$ cells. Cumulative antigenic stimulation is denoted $a$. The total concentrations of primary and recall effectors are given by

$$E(t) = \int_0^{\infty} \bar{E}(a,t)\,da \qquad \text{(S3)}$$

$$R(t) = \int_0^{\infty} \bar{R}(a,t)\,da \qquad \text{(S4)}$$

We account for loss, $\Lambda_i$, by death and differentiation into memory cells. These processes are modeled as first-order and saturable processes with rate constants $d_E$ and $r_{EM}$, respectively, for $\bar{E}$, and $d_R$ and $r_{RM}$ for $\bar{R}$:

$$\Lambda_{\bar{E}} = \left[ d_E + r_{EM}\left(1 - \frac{M}{\phi_M}\right) \right] \bar{E} \qquad \text{(S5)}$$

$$\Lambda_{\bar{R}} = \left[ d_R + r_{RM}\left(1 - \frac{M}{\phi_M}\right) \right] \bar{R} \qquad \text{(S6)}$$

$M$ is the total size of the memory CD8 T cell pool and $\phi_M$ is its carrying capacity.

Gain, $\Gamma_i$, is by two processes: (a) Production of effectors from their respective progenitors. The progenitors for $\bar{E}$ are $\bar{P}$ and for $\bar{R}$ are $\bar{M}$, respectively. (b) Proliferation of effectors. These processes require that the progenitors (for recruitment) and the effectors (for proliferation) form a complex with antigen-bearing antigen-presentation cells (APCs)[1]. Below, we first estimate the concentration of APCs that harbor antigen, given the concentration of total APCs and the antigen load in the host. Then we estimate how many of these antigen-bearing APCs engage with primary ($\bar{E}/\bar{P}$) and recall ($\bar{R}/\bar{M}$) cells by accounting for competition[2]. Finally, $\Gamma_{\bar{E}}$ and $\Gamma_{\bar{R}}$ are modeled as proportional to the levels of antigen-bearing APCs complexed with their respective CD8 T cell populations.

**Concentration of antigen-bearing APCs**

Let $G_T$ and $G_A$ denote the concentrations of total and antigen-bearing APCs, respectively. APCs without antigen, $G_F$, take up antigen from their milieu for presentation to CD8 T cells. Viral peptide–loaded MHCs (pMHCs) presented on the surface of the antigen-bearing APCs have a specific half-life[3], which results in the transition of $G_A$ back to $G_F$. These two processes are captured in the reaction scheme below.

$$G_F + I \xrightarrow{k_1} G_A \xrightarrow{k_2} G_F \tag{S7}$$

Here, $I$ is the concentration of infected cells, the marker for antigen load, while $k_1$ and $k_2$ are rate constants. Assuming quasi steady state for $G_A$[4], we get $k_1 I G_F^{ss} \approx k_2 G_A^{ss}$, where the superscript $ss$ indicates steady state. Because $G_T = G_F^{ss} + G_A^{ss}$, it follows that $k_1 I \left(G_T - G_A^{ss}\right) = k_2 G_A^{ss}$ and hence

$$G_A^{ss} = G_T \frac{I}{\theta + I} \quad \text{(S8)}$$

Here, $\theta = k_2 / k_1$. Thus, the number of antigen-bearing APCs, $G_A^{ss}$, increases with the antigen load and saturates to the maximum number of APCs, $G_T$. Next, we derive how many of the $G_A^{ss}$ cells complex with $\bar{E} / \bar{P}$ and $\bar{R} / \bar{M}$ cells.

**Number of antigen-bearing APCs complexed with $\bar{E} / \bar{P}$ and $\bar{R} / \bar{M}$ cells**

We assume that an antigen-bearing APC is in one of the three possible states: signaling to $\bar{E} / \bar{P}$ cells, signaling to $\bar{R} / \bar{M}$ cells, and free from engagement with any CD8 T cells. Because effector cells have high proliferation rates and an APC engages at most 10 cells at once[5], we assume that an engaged APC is surrounded by and exclusively signals to either $\bar{E} / \bar{P}$ or $\bar{R} / \bar{M}$ cells. Let $\bar{Z}_E$ and $\bar{Z}_R$ denote the densities of antigen-bearing APCs complexed to cells of primary ($\bar{P}$ or $\bar{E}$) and recall ($\bar{M}$ or $\bar{R}$) subtype, respectively. Following (S3) and (S4), $Z_E = \int_0^\infty \bar{Z}_E da$ and $Z_R = \int_0^\infty \bar{Z}_R da$ are the aggregate levels of antigen-bearing APCs complexed to all CD8 T cells of primary and recall subtypes, respectively. Let $Z_F$ denote the concentration of free antigen-bearing APCs. The dynamical equations describing this system are below (Fig. S23).

$$\frac{d\bar{Z}_E}{dt} = \left(\eta_P \bar{P} + \eta_E \bar{E}\right) Z_F - \eta_F \bar{Z}_E + \mu_1 \left(\bar{P}^m + \bar{E}^m\right) \int_0^\infty \bar{Z}_R da - \mu_2 \left(M^m + R^m\right) \bar{Z}_E \quad \text{(S9)}$$

$$\frac{d\bar{Z}_R}{dt} = \left(\eta_M \bar{M} + \eta_R \bar{R}\right) Z_F - \eta_F \bar{Z}_R + \mu_1 \left(\bar{M}^m + \bar{R}^m\right) \int_0^\infty \bar{Z}_E da - \mu_2 \left(P^m + E^m\right) \bar{Z}_R \quad \text{(S10)}$$

$$\frac{dZ_F}{dt} = \int_0^\infty \eta_F \left(\bar{Z}_E + \bar{Z}_R\right) da - \int_0^\infty \left(\eta_P \bar{P} + \eta_E \bar{E} + \eta_M \bar{M} + \eta_R \bar{R}\right) Z_F da \quad \text{(S11)}$$

Equation (S9) for $\bar{Z}_E$ accounts for the following processes. Free antigen-bearing APCs, $Z_F$, complex with $\bar{P}$ or $\bar{E}$ in the milieu and become $\bar{Z}_E$ [6]. The rate constants are $\eta_P$ and $\eta_E$, respectively. APCs of $\bar{Z}_E$ could disengage and become free at the rate $\eta_F \bar{Z}_E$. As the concentration of recall cells increases in the milieu, $\bar{Z}_E$ APCs get surrounded by them. This may cause the APCs to switch their engagement to $\bar{R}/\bar{M}$ cells. The terms $\mu_2 M^m \bar{Z}_E$ and $\mu_2 R^m \bar{Z}_E$ denote these processes, where $\mu_2$ is the rate constant and $m$ is the exponent for competitive exclusion of APCs $\bar{Z}_E$ by recall cells of any $a$. Since the replacement of $\bar{E}/\bar{P}$ cells surrounding $\bar{Z}_E$ requires multiple $\bar{R}/\bar{M}$ cells to surround the corresponding APC, we let $m > 1$ to account for the associated cooperative effects. Similarly, $\mu_1 \bar{P}^m \int_0^\infty \bar{Z}_R da$ and $\mu_1 \bar{E}^m \int_0^\infty \bar{Z}_R da$ denote the switching of the engagement of $\bar{Z}_R$ APCs, with $\mu_1$ the rate constant. Equation (S10) for $\bar{Z}_R$ accounts for analogous processes: $Z_F$ engagement (with $\bar{R}/\bar{M}$ cells), disengagement, $Z_E$ engagement switching, and competitive exclusion (by $\bar{E}/\bar{P}$ cells). Equation (S11) for $Z_F$ describes the loss of antigen-bearing APCs due to engagement with either of the four CD8 T cell types and gain with their disengagement.

We employ two assumptions to simplify the above equations. First, progenitor cells are much lower in concentration than their respective effector cells due to the high proliferation rates of the latter [7]. Thus, $E^m \gg P^m$ and $R^m \gg M^m$. We therefore neglect the terms with progenitors in the terms for engagement switching. Second, gain from effectors engaging with free APCs is likely to be far more than the gain from engaged APCs switching their engagement to alternative CD8 T cell subtypes. We thus neglect the switching-based gain terms in (S9) and (S10). With these two assumptions, the equations above become

$$\frac{d\bar{Z}_E}{dt} = \left(\eta_P \bar{P} + \eta_E \bar{E}\right) Z_F - \eta_F \bar{Z}_E - \mu_2 R^m \bar{Z}_E \tag{S12}$$

$$\frac{d\bar{Z}_R}{dt} = \left(\eta_M \bar{M} + \eta_R \bar{R}\right) Z_F - \eta_F \bar{Z}_R - \mu_2 E^m \bar{Z}_R \tag{S13}$$

$$\frac{dZ_F}{dt} = \eta_F \left(G_A^{ss} - Z_F\right) - \left(\eta_P P + \eta_E E + \eta_M M + \eta_R R\right) Z_F \tag{S14}$$

Here, $Z_F + Z_E + Z_R = G_A^{ss}$. Because these processes of engagement evolve considerably faster than cellular events like proliferation and death [5], we assume quasi steady state for equations (S12)–(S14), yielding

$$Z_F^{ss} = \frac{\eta_F}{\eta_F + \left(\eta_P P + \eta_E E + \eta_M M + \eta_R R\right)} G_A^{ss} \tag{S15}$$

$$\bar{Z}_E^{ss} = \frac{\left(\eta_P \bar{P} + \eta_E \bar{E}\right)}{\eta_F + \mu_2 R^m} Z_F^{ss} \tag{S16}$$

$$\bar{Z}_R^{ss} = \frac{\left(\eta_M \bar{M} + \eta_R \bar{R}\right)}{\eta_F + \mu_2 E^m} Z_F^{ss} \tag{S17}$$

The superscript $ss$ indicates steady state. APCs are highly motile and rapidly scan their milieu and complex with CD8 T cells. An APC engages with hundreds of T cells in an hour[5]. We therefore assume that an APC disengages after signaling a set of CD8 T cells and scans the milieu for other CD8 T cells rather than stay engaged with the former for a long time. Accordingly, in equation (S15), we let $\eta_F \gg \left(\eta_P P + \eta_E E + \eta_M M + \eta_R R\right)$, which yields $Z_F^{ss} \approx G_A^{ss} = G_T \dfrac{I}{\theta + I}$ from equation (S8). Hence,

$$\bar{Z}_E^{ss} = \left(\frac{\eta_P G_T}{\eta_F} \bar{P} + \frac{\eta_E G_T}{\eta_F} \bar{E}\right) \frac{I}{\theta + I} \frac{(\eta_F / \mu_2)}{(\eta_F / \mu_2) + R^m} \tag{S18}$$

$$\bar{Z}_R^{ss} = \left( \frac{\eta_M G_T}{\eta_F} \bar{M} + \frac{\eta_R G_T}{\eta_F} \bar{R} \right) \frac{I}{\theta + I} \frac{(\eta_F / \mu_2)}{(\eta_F / \mu_2) + E^m} \quad \text{(S19)}$$

Let $\eta_F / \mu_2 = \theta_C^m$. Substituting this into (S18) and (S19) yields

$$\bar{Z}_E^{ss} = \left( \frac{\eta_P G_T}{\eta_F} \bar{P} + \frac{\eta_E G_T}{\eta_F} \bar{E} \right) \frac{I}{\theta + I} \frac{\theta_C^m}{\theta_C^m + R^m} \quad \text{(S20)}$$

$$\bar{Z}_R^{ss} = \left( \frac{\eta_M G_T}{\eta_F} \bar{M} + \frac{\eta_R G_T}{\eta_F} \bar{R} \right) \frac{I}{\theta + I} \frac{\theta_C^m}{\theta_C^m + E^m} \quad \text{(S21)}$$

The gain terms, $\Gamma_{\bar{E}}$ and $\Gamma_{\bar{R}}$, in equations (S1) and (S2) are proportional to $\bar{Z}_E^{ss}$ and $\bar{Z}_R^{ss}$, respectively. Assuming the proportionality constant to be $w$ and substituting (S18) and (S19) in the former equations yields

$$\frac{\partial \bar{E}}{\partial t} + \frac{da}{dt} \frac{\partial \bar{E}}{\partial a} = \left( \frac{w \eta_P G_T}{\eta_F} \bar{P} + \frac{w \eta_E G_T}{\eta_F} \bar{E} \right) \frac{I}{\theta + I} \frac{\theta_C^m}{\theta_C^m + R^m} - \left[ d_E + r_{EM} \left( 1 - \frac{M}{\phi_M} \right) \right] \bar{E} \quad \text{(S22)}$$

$$\frac{\partial \bar{R}}{\partial t} + \frac{da}{dt} \frac{\partial \bar{R}}{\partial a} = \left( \frac{w \eta_M G_T}{\eta_F} \bar{M} + \frac{w \eta_R G_T}{\eta_F} \bar{R} \right) \frac{I}{\theta + I} \frac{\theta_C^m}{\theta_C^m + E^m} - \left[ d_R + r_{RM} \left( 1 - \frac{M}{\phi_M} \right) \right] \bar{R} \quad \text{(S23)}$$

Let $w \eta_P G_T / \eta_F = \omega_P$, $w \eta_E G_T / \eta_F = \alpha_E$, $w \eta_M G_T / \eta_F = \omega_M$, and $w \eta_R G_T / \eta_F = \alpha_R$. Substituting these terms into (S22) and (S23) gives

$$\frac{\partial \bar{E}}{\partial t} + \frac{da}{dt} \frac{\partial \bar{E}}{\partial a} = \left( \omega_P \bar{P} + \alpha_E \bar{E} \right) \frac{I}{\theta + I} \frac{\theta_C^m}{\theta_C^m + R^m} - \left[ r_{EM} \left( 1 - \frac{M}{\phi_M} \right) + d_E \right] \bar{E} \quad \text{(S24)}$$

$$\frac{\partial \bar{R}}{\partial t} + \frac{da}{dt} \frac{\partial \bar{R}}{\partial a} = \left( \omega_M \bar{M} + \alpha_R \bar{R} \right) \frac{I}{\theta + I} \frac{\theta_C^m}{\theta_C^m + E^m} - \left[ r_{RM} \left( 1 - \frac{M}{\phi_M} \right) + d_R \right] \bar{R} \quad \text{(S25)}$$

These are equations (12) and (13) in the main text.

### Text S2: Transformed model with population-averaged cumulative antigenic stimulation

Equations (11)–(14) in the main text describe activated CD8 T cell populations as they accumulate cumulative antigenic stimulation, $a$. Different cells accumulate different levels of this marker over the course of infection, $t$, starting from their initial activation time. Most virus-specific naïve cells present at the time of infection are activated within a short time window immediately after inoculation. Because new naïve cells are produced at low levels[8], the influx of naïve cells into the activated CD8 T cell pool is negligible shortly after this time window. Thus, all activated cells may be assumed to begin accumulating antigenic stimulation at the time of inoculation. All cells present at any time $t$ post-inoculation will thus have the same cumulative antigenic stimulation $A$. We recognize that $A$ may be linked to $a$ as the population average:

$$A = \frac{\int_0^\infty a\bar{P}(a,t)da}{\int_0^\infty \bar{P}(a,t)da} \tag{S26}$$

Below, we (a) incorporate this link to transform equations (11)–(14) in the main text and describe the dynamics of activated CD8 T cells in terms of $A$, and (b) derive a dynamical equation for $A$.

**Deriving the dynamical equations for the total levels of activated CD8 T cell subtypes**

We convert the equations (11)–(14) in the main text, one by one, by integrating the variable $a$ out and using the definition in (S26). The definition $\chi = \chi(t) \equiv \int_0^\infty \bar{\chi}(a,t)da \;\; \forall\; \chi \in \{P, E, R, M\}$ and the boundary condition $\lim_{a\to\infty} \bar{\chi}(a,t) = 0$ are used throughout these derivations.

We begin with Equation (11): $\frac{\partial \bar{P}}{\partial t}+\frac{da}{dt}\frac{\partial \bar{P}}{\partial a}=\alpha_P \bar{P}\left(1-\frac{P}{\phi_P}\right)\frac{I}{\theta+I}$. Integrating $a$ out yields $\int_0^\infty \frac{\partial \bar{P}}{\partial t}da+\int_0^\infty \frac{da}{dt}\frac{\partial \bar{P}}{\partial a}da=\alpha_P\left(1-\frac{P}{\phi_P}\right)\frac{I}{\theta+I}\int_0^\infty \bar{P}da$. Interchanging the order of integration and differentiation, and integrating yields $\frac{dP}{dt}+\frac{da}{dt}\left[\bar{P}(a,t)\right]_{a=0}^{a\to\infty}=\alpha_P\left(1-\frac{P}{\phi_P}\right)P\frac{I}{\theta+I}$, which upon applying the boundary condition above becomes $\frac{dP}{dt}+\frac{da}{dt}\left[0-\bar{P}(0,t)\right]=\alpha_P P\left(1-\frac{P}{\phi_P}\right)\frac{I}{\theta+I}$. Naïve cells after initial activation become progenitor cells and start accumulating antigenic stimulation. Thus, $\bar{P}(0,t)da=N_p dt$, the number of naïve cells activated in time $dt$. Drawing the activation term from the equation for naïve cells and combining it with the equation for $\frac{da}{dt}$ yields $\bar{P}(0,t)=\left(r_{NP}N\frac{I}{\theta_N+I}\right)/\left(\alpha_A\frac{I^n}{\theta_A^n+I^n}\right)$. Substituting these in the equation above yields an ordinary differential equation for $P$:

$$\frac{dP}{dt}=r_{NP}N\frac{I}{\theta_N+I}+\alpha_P P\left(1-\frac{P}{\phi_P}\right)\frac{I}{\theta+I} \quad \text{(S27)}$$

Using the same procedure for Equation (12), $\frac{\partial \bar{E}}{\partial t}+\frac{da}{dt}\frac{\partial \bar{E}}{\partial a}=\left(\omega_P\bar{P}+\alpha_E\bar{E}\right)\frac{I}{\theta+I}\frac{\theta_C^m}{\theta_C^m+R^m}-\left[r_{EM}\left(1-\frac{M}{\phi_M}\right)+d_E\right]\bar{E}$, and recognizing that $\bar{E}(0,t)=0$ yields the ordinary differential equation for $E$.

$$\frac{dE}{dt}=\left(\omega_P P+\alpha_E E\right)\frac{I}{\theta+I}\frac{\theta_C^m}{\theta_C^m+R^m}-\left[r_{EM}\left(1-\frac{M}{\phi_M}\right)+d_E\right]E \quad \text{(S28)}$$

In an analogous manner, Equation (13), $\frac{\partial \bar{R}}{\partial t}+\frac{da}{dt}\frac{\partial \bar{R}}{\partial a}=\left(\omega_M\bar{M}+\alpha_R\bar{R}\right)\frac{I}{\theta+I}\frac{\theta_C^m}{\theta_C^m+E^m}-\left[r_{RM}\left(1-\frac{M}{\phi_M}\right)+d_R\right]\bar{R}$, becomes

$$\frac{dR}{dt} = \left(\omega_M M + \alpha_R R\right)\frac{I}{\theta + I}\frac{\theta_C^m}{\theta_C^m + E^m} - \left[r_{RM}\left(1 - \frac{M}{\phi_M}\right) + d_R\right]R \tag{S29}$$

Finally, Equation (14), $\frac{\partial \bar{M}}{\partial t} + \frac{da}{dt}\frac{\partial \bar{M}}{\partial a} = \left(r_{EM}\bar{E} + r_{RM}\bar{R}\right)\left(1 - \frac{M}{\phi_M}\right) - d_X\frac{I}{\theta + I}\mu(a)\bar{M}$, applying the same procedure yields $\frac{dM}{dt} = \left(r_{EM}E + r_{RM}R\right)\left(1 - \frac{M}{\phi_M}\right) - d_X\frac{I}{\theta + I}\int_0^\infty \mu(a)\bar{M}da$. Here, $\mu(a)$ is an increasing function of $a$, accounting for the increasing death rate of the memory cells with higher cumulative antigenic stimulation. At the population level, we let the average death rate be an increasing-saturating function of $A$. Thus, we write $\frac{1}{M}\int_0^\infty \mu(a)\bar{M}da = \frac{A}{\theta_X + A}$, where $\theta_X$ is the half-maximal sensitivity constant for the dependence of memory cell death rate on $A$. We assumed an increasing functional form here. The form is consistent with experimental studies, which suggest that the death rate of memory CD8 T cells progressively increases with accumulation of antigenic stimulation and becomes equivalent to that of terminally differentiated cells[9,10]. For instance, Angelosanto et al. report that in a chronic LCMV infection, memory CD8 T cells in mice are substantially lost by day-60 post-infection[9]. However, they found that the process begins by ~day-15 post-infection and as the infection progresses into the chronic phase, it becomes difficult for the memory functionality to be recovered. Such irreversible loss of memory functionality and commitment to dysfunction has been reported in many human studies as well[11]. In light of these reports, we chose the present form.

Using this form, we obtain an ordinary differential equation for $M$:

$$\frac{dM}{dt} = \left(r_{EM}E + r_{RM}R\right)\left(1 - \frac{M}{\phi_M}\right) - d_X\frac{I}{\theta + I}\frac{A}{\theta_X + A}M \tag{S30}$$

**Deriving the dynamical equation for $A$**

We know that $\int_0^\infty \bar{P}(a,t)da = P$. Substituting this in equation (S26) yields $PA = \int_0^\infty a\bar{P}(a,t)da$. Differentiating the above expression with respect to time $t$ results in

$$\frac{d}{dt}(PA) = \frac{d}{dt}\left[\int_0^\infty a\bar{P}da\right]$$

$$\Rightarrow A\frac{dP}{dt} + P\frac{dA}{dt} = \int_0^\infty \left[a\frac{d\bar{P}}{dt} + \bar{P}\frac{da}{dt}\right]da$$

$$\Rightarrow \frac{dA}{dt} = \frac{\Omega}{P} - \frac{A}{P}\frac{dP}{dt} \tag{S31}$$

Here, $\Omega = \int_0^\infty \left[a\frac{d\bar{P}}{dt} + \bar{P}\frac{da}{dt}\right]da$. The integral $\Omega$ is solved below.

$$\begin{aligned}\Omega &= \int_0^\infty \bar{P}\frac{da}{dt}da + \int_0^\infty a\frac{d\bar{P}}{dt}da \\ &= \frac{da}{dt}\int_0^\infty \bar{P}da + \int_0^\infty a\left[\frac{\partial\bar{P}}{\partial t} + \frac{da}{dt}\frac{\partial\bar{P}}{\partial a}\right]da \\ &= P\frac{da}{dt} + \int_0^\infty a\left[\alpha_P\bar{P}\left(1-\frac{P}{\phi_P}\right)\frac{I}{\theta+I}\right]da \\ &= P\frac{da}{dt} + \alpha_P\left(1-\frac{P}{\phi_P}\right)\frac{I}{\theta+I}\int_0^\infty a\bar{P}da \\ &= P\frac{da}{dt} + \alpha_P\left(1-\frac{P}{\phi_P}\right)\frac{I}{\theta+I}PA\end{aligned}$$

$$\Rightarrow \frac{\Omega}{P} = \frac{da}{dt} + \alpha_P A\left(1-\frac{P}{\phi_P}\right)\frac{I}{\theta+I} \tag{S32}$$

Substituting (S32) into (S31) yields

$$\frac{dA}{dt} = \frac{da}{dt} + \alpha_P A\left(1-\frac{P}{\phi_P}\right)\frac{I}{\theta+I} - \frac{A}{P}\left[r_{NP}N\frac{I}{\theta_N+I} + \alpha_P P\left(1-\frac{P}{\phi_P}\right)\frac{I}{\theta+I}\right]$$

$$\Rightarrow \frac{dA}{dt} = \frac{da}{dt} - r_{NP}\frac{N}{P}\frac{I}{\theta_N + I}A$$

Substituting the expression for $\frac{da}{dt}$ above results in

$$\frac{dA}{dt} = \alpha_A \frac{I^n}{\theta_A^n + I^n} - r_{NP}\frac{N}{P}\frac{I}{\theta_N + I}A \quad \text{(S33)}$$

The above equation is the dynamical equation for $A$. Equations (S27)–(S30) and (S33) comprise the dynamical equations for the activated CD8 T cells in our transformed model. The complete set of equations are given below.

$$\frac{dT}{dt} = \lambda - (1-\epsilon)\beta TV - d_T T \quad \text{(S34)}$$

$$\frac{dI}{dt} = (1-\epsilon)(1 - f_L - f_D)\beta TV + \rho L - k(E+R)I - d_I I \quad \text{(S35)}$$

$$\frac{dL}{dt} = (1-\epsilon) f_L \beta TV - \rho L - d_L L \quad \text{(S36)}$$

$$\frac{dD}{dt} = (1-\epsilon) f_D \beta TV - \psi_D kED - d_D D \quad \text{(S37)}$$

$$\frac{dV}{dt} = pI - d_V V \quad \text{(S38)}$$

$$\frac{dN}{dt} = \lambda_N - r_{NP} N \frac{I}{\theta_N + I} - d_N N \quad \text{(S39)}$$

$$\frac{dA}{dt} = \alpha_A \frac{I^n}{\theta_A^n + I^n} - r_{NP}\frac{N}{P}\frac{I}{\theta_N + I}A \quad \text{(S40)}$$

$$\frac{dP}{dt} = r_{NP} N \frac{I}{\theta_N + I} + \alpha_P P\left(1 - \frac{P}{\phi_P}\right)\frac{I}{\theta + I} \quad \text{(S41)}$$

$$\frac{dE}{dt} = \left(\omega_P P + \alpha_E E\right)\frac{I}{\theta + I}\frac{\theta_C^m}{\theta_C^m + R^m} - \left[r_{EM}\left(1 - \frac{M}{\phi_M}\right) + d_E\right]E \tag{S42}$$

$$\frac{dR}{dt} = \left(\omega_M M + \alpha_R R\right)\frac{I}{\theta + I}\frac{\theta_C^m}{\theta_C^m + E^m} - \left[r_{RM}\left(1 - \frac{M}{\phi_M}\right) + d_R\right]R \tag{S43}$$

$$\frac{dM}{dt} = \left(r_{EM} E + r_{RM} R\right)\left(1 - \frac{M}{\phi_M}\right) - d_X \frac{I}{\theta + I}\frac{A}{\theta_X + A} M \tag{S44}$$

Following earlier studies, we assumed a quasi–steady state between viral production and clearance rates[12] and simplified the equation for viremia (S38), resulting in $pI \approx d_V V \Rightarrow V = \gamma I$ , where $\gamma = p / d_V$. Thus, equations (S34)–(S44) are the equations (16)–(25) in the main text employed for our study.

### Text S3: Model equations for linear stability analysis

Following are the set of equations solved to obtain the steady states of our model.

$$\frac{dI_{\text{ss}}}{dt}=0 \Rightarrow (1-\epsilon)(1-f_L-f_D)\beta^* T_{\text{ss}} I_{\text{ss}} + \rho L_{\text{ss}} - (E^*_{\text{ss}}+R^*_{\text{ss}})I_{\text{ss}} - d_I I_{\text{ss}} = 0 \tag{S45}$$

$$\frac{dE^*_{\text{ss}}}{dt}=0 \;\Rightarrow\; \left(\omega_P P^*_{\text{ss}} + \alpha_E E^*_{\text{ss}}\right)\frac{I_{\text{ss}}}{\theta+I_{\text{ss}}}\frac{\theta_C^{*m}}{\theta_C^{*m}+R_{\text{ss}}^{*m}} - \left[r_{EM}\left(1-\frac{M^*_{\text{ss}}}{\phi^*_M}\right)+d_E\right]E^*_{\text{ss}} = 0 \tag{S46}$$

$$\frac{dR^*_{\text{ss}}}{dt}=0 \;\Rightarrow\; \left(\omega_M M^*_{\text{ss}} + \alpha_R R^*_{\text{ss}}\right)\frac{I_{\text{ss}}}{\theta+I_{\text{ss}}}\frac{\theta_C^{*m}}{\theta_C^{*m}+E_{\text{ss}}^{*m}} - \left[r_{RM}\left(1-\frac{M^*_{\text{ss}}}{\phi^*_M}\right)+d_R\right]R^*_{\text{ss}} \tag{S47}$$

Here, $T_{\text{ss}}=\dfrac{\lambda}{\beta^* I_{\text{ss}}+d_T}$, $L_{\text{ss}}=\dfrac{f_L\beta^* T_{\text{ss}} I_{\text{ss}}}{\rho+d_L}$, $D_{\text{ss}}=\dfrac{f_D\beta^* T_{\text{ss}} I_{\text{ss}}}{\psi_D E^*_{\text{ss}}+d_D}$, $N^*_{\text{ss}}=\dfrac{\lambda^*_N}{r_{NP}\dfrac{I_{\text{ss}}}{\theta_N+I_{\text{ss}}}+d_N}$, $P^*_{\text{ss}}=\phi^*_P$,

$M^*_{\text{ss}}=\dfrac{\left(r_{EM}E^*_{\text{ss}}+r_{RM}R^*_{\text{ss}}\right)}{\left[\left(r_{EM}E^*_{\text{ss}}+r_{RM}R^*_{\text{ss}}\right)\dfrac{1}{\phi^*_M}+d_X\dfrac{A^*_{\text{ss}}}{1+A^*_{\text{ss}}}\dfrac{I_{\text{ss}}}{\theta+I_{\text{ss}}}\right]}$, and $A^*_{\text{ss}}=\dfrac{\alpha^*_A\dfrac{I^n_{\text{ss}}}{\theta^n_A+I^n_{\text{ss}}}}{r_{NP}\dfrac{N^*_{\text{ss}}}{P^*_{\text{ss}}}\dfrac{I_{\text{ss}}}{\theta_N+I_{\text{ss}}}}$. These expressions were obtained by equating the respective dynamical equations (equations (16)–(25) from the main text) to zero. After solving the equations (S45)–(S47), for each steady state, the values $I_{\text{ss}}$, $E^*_{\text{ss}}$, and $R^*_{\text{ss}}$ were substituted back into the steady state expressions for the other variables to calculate the values of the latter for the corresponding steady state.

### Text S4: Origin of the third stable steady state of our model

The model also predicts a third stable steady state over short spans of some parameter ranges (Fig. S3, S4). This state is defined by high viral load, similar to that of the progressive infection state, with modest recall responses (Fig. S4a–c). Interestingly, the memory pool is smaller than the naïve-derived progenitor pool (source for primary responses) (Fig. S4d), with a high level of cumulative antigenic stimulation (Fig. S4e). Yet this state is stable because the recall responses are large enough to suppress the emergence of naïve-derived primary responses by engaging with most APCs, and the existence of this state is restricted to conditions like low death rates (Fig. S3) and high proliferation rates (Fig. S4) of effector cells, wherein the recall effectors could be maintained at high levels despite negligible recruitment from their progenitors, i.e., memory cells. However, this state would be inaccessible because effector cells are derived from their respective progenitors and it is unlikely for an effector response to be mounted when its progenitors are at negligible levels[2,4].

### Text S5: Our model equations fit in the first stage

We fit our model in two stages. In the first stage, we fit the part of our model described in equations (16)–(25) of the main text without latently infected cells, memory cells, and recall responses to pre-ART data of the three groups. Equations that fit in the first stage are provided below.

$$\frac{dT}{dt} = \lambda - \beta^* TI - d_T T \tag{S48}$$

$$\frac{dI}{dt} = (1 - f_D)\beta^* TI - E^* I - d_I I \tag{S49}$$

$$\frac{dD}{dt} = f_D \beta^* TI - \psi_D E^* D - d_D D \tag{S50}$$

$$\frac{dN^*}{dt} = \lambda_N^* - r_{NP} N^* \frac{I}{\theta_N + I} - d_N N^* \tag{S51}$$

$$\frac{dA^*}{dt} = \alpha_A^* \frac{I^n}{\theta_A^n + I^n} - r_{NP} \frac{N^*}{P^*} \frac{I}{\theta_N + I} A^* \tag{S52}$$

$$\frac{dP^*}{dt} = r_{NP} N^* \frac{I}{\theta_N + I} + \alpha_P P^* \left(1 - \frac{P^*}{\phi_P^*}\right) \frac{I}{\theta + I} \tag{S53}$$

$$\frac{dE^*}{dt} = \left(\omega_P P^* + \alpha_E E^*\right) \frac{I}{\theta + I} - \left[r_{EM} + d_E\right] E^* \tag{S54}$$

Here, $V = \gamma I$ and we estimated $\beta^*$, $\gamma$, $\psi_D$, $\omega$, $\alpha_E$, and $T(0)$ in this stage.

### Text S6: Existing models from the literature fit to pVISCONTI data

We examined whether existing mathematical models could also describe the pVISCONTI data. We considered two models from the literature: (a) model #1, proposed by Conway and Perelson[13], and (b) model #2, a variant of model #1 that accounts for exhaustion of CD8 T cells as described by Johnson et al.[14]. Fits of these models are compared to those of our model. The fitting procedure is detailed below, followed by a summary of the fits.

**Model #1**

$$\frac{dT}{dt} = \lambda - (1-\epsilon)\beta TV - d_T T \tag{S55}$$

$$\frac{dI}{dt} = (1-f_L)(1-\epsilon)\beta TV + \rho L - kEI - d_I I \tag{S56}$$

$$\frac{dL}{dt} = f_L(1-\epsilon)\beta TV - \rho L - d_L L \tag{S57}$$

$$\frac{dE}{dt} = \lambda_E + \alpha_E E \frac{I}{\theta_E + I} - \alpha_X E \frac{I}{\theta_X + I} - d_E E \tag{S58}$$

$$\frac{dV}{dt} = pI - d_V V \tag{S59}$$

The equations describing uninfected target cells, different types of infected cells, and free virions are similar to those employed in our model with memory CD8 T cells. The equation for effector cell level, $E$, encodes four processes: recruitment, proliferation, exhaustion, and death. Effector cells are recruited at the rate $\lambda_E$, and they have a per-capita proliferation rate of $\alpha_E$. These cells are lost due to exhaustion at a per-capita rate $\alpha_X$ or due to natural death at rate $d_E E$. Proliferation and exhaustion are regulated by antigen levels, with $\theta_E$ and $\theta_X$ the respective half-maximal saturation constants.

We incorporated the quasi–steady state approximation for virion production and clearance rates[12], similar to that in our model, yielding $V=\gamma I$. $k$ was subsumed into $E$, yielding $E^{*}=kE$ and $\lambda_{E}^{*}=k\lambda_{E}$. Thus, the final set of equations fit to data are

$$\frac{dT}{dt}=\lambda-(1-\epsilon)\beta^{*}TI-d_{T}T \tag{S60}$$

$$\frac{dI}{dt}=(1-f_{L})(1-\epsilon)\beta^{*}TI+\rho L-E^{*}I-d_{I}I \tag{S61}$$

$$\frac{dL}{dt}=f_{L}(1-\epsilon)\beta^{*}TI-\rho L-d_{L}L \tag{S62}$$

$$\frac{dE^{*}}{dt}=\lambda_{E}^{*}+\alpha_{E}E^{*}\frac{I}{\theta_{E}+I}-\alpha_{X}E^{*}\frac{I}{\theta_{X}+I}-d_{E}E^{*} \tag{S63}$$

Prior to fitting, we assessed parameter identifiability to identify estimable parameters. Of those parameters, $\beta^{*}(=\gamma\beta)$, $\gamma$, $\epsilon$, $\alpha_{E}$, and $T(0)$ were estimable (Methods in main text). All other parameters were fixed to values from literature. Fits are presented in Fig. S11–S13 and the VPCs are in Fig. S14.

**Model #2**

$$\frac{dT}{dt}=\lambda-(1-\epsilon)\beta TV-d_{T}T \tag{S64}$$

$$\frac{dI}{dt}=(1-f_{L}-f_{D})(1-\epsilon)\beta TV+\rho L-kEI-d_{I}I \tag{S65}$$

$$\frac{dD}{dt}=f_{D}(1-\epsilon)\beta TV-d_{D}D \tag{S66}$$

$$\frac{dL}{dt}=f_{L}(1-\epsilon)\beta TV-\rho L-d_{L}L \tag{S67}$$

$$\frac{dE}{dt} = \lambda_E + \alpha_E E \frac{I}{\theta_E + I} - \xi E \frac{Q^w}{q_c{}^w + Q^w} - d_E E \tag{S68}$$

$$\frac{dQ}{dt} = \kappa \frac{I}{\theta_E + I} - d_Q Q \tag{S69}$$

$$\frac{dV}{dt} = pI - d_V V \tag{S70}$$

The model is similar to the Conway-Perelson model, except for how exhaustion has been described. In the Conway-Perelson model, exhaustion levels depend on the instantaneous antigen levels, while here, exhaustion level, $Q$, builds up and decays over time with exposure to antigen. The maximal exhaustion build up rate is $\kappa$ and decay rate constant is $d_Q$. Maximal per-capita rate of exhaustion-dependent loss of effector cells is $\xi$, half-maximal saturation constant for the exhaustion-dependent loss is $q_c$, and the Hill coefficient is $w$.

Similar to that in model #1, we incorporated the quasi–steady state approximation for virion production and clearance rates[12], yielding $V = \gamma I$. $k$ was subsumed into $E$, yielding $E^* = kE$ and $\lambda_E^* = k\lambda_E$. Thus, the final set of equations fit to data are

$$\frac{dT}{dt} = \lambda - (1-\epsilon)\beta^* TI - d_T T \tag{S71}$$

$$\frac{dI}{dt} = (1 - f_L - f_D)(1-\epsilon)\beta^* TI + \rho L - E^* I - d_I I \tag{S72}$$

$$\frac{dD}{dt} = f_D (1-\epsilon)\beta^* TI - d_D D \tag{S73}$$

$$\frac{dL}{dt} = f_L (1-\epsilon)\beta^* TI - \rho L - d_L L \tag{S74}$$

$$\frac{dE^*}{dt} = \lambda_E^* + \alpha_E E^* \frac{I}{\theta_E + I} - \xi E^* \frac{Q^w}{q_c{}^w + Q^w} - d_E E^* \tag{S75}$$

$$\frac{dQ}{dt} = \kappa \frac{I}{\theta_E + I} - d_Q Q \tag{S76}$$

Prior to fitting, we first assessed parameter identifiability to identify estimable parameters. Of those parameters, $\beta^*(=\gamma\beta)$, $\gamma$, $\epsilon$, $\alpha_E$, and $T(0)$ were estimable (Methods in main text). All other parameters were fixed to values from literature. Fits are presented in Fig. S15–S17 and the VPCs are in Fig. S18.

**Fitting the models to data**

The procedure for fitting both the above models involved two stages, similar to what has been employed to fit our model (Methods in main text). In the first stage, we fit the model without the exhaustion terms. That corresponds to the term $\alpha_X E^* \frac{I}{\theta_X + I}$ in model #1 and the term $\xi E^* \frac{Q^w}{q_c^{\,w} + Q^w}$ as well as the equation $dQ/dt$ in model #2. We estimated parameters $\beta^*$, $\gamma$, $\alpha_E$, and $T(0)$ in both the models in this stage (Table S5, S6). Then, in the second stage, we fit full models to the complete data to additionally estimate $\epsilon$. Outcomes are different between macaques only post-treatment and according to these models, the differences could be explained by difference in CD8 T cell exhaustion and proviral reservoir levels. So, we tried estimating parameters related to exhaustion only in the second stage. Note that parameters that govern reservoir dynamics, like $f_L$, $f_D$, $d_L$, $d_D$, and $\rho$ were not estimable.

**Inference**

The fits could not distinguish between PTCs and CPs (Fig. S11–S19, Table S5–S7). The resulting population parameter estimates predicted that both early and late treatment initiation would result in the same outcome, i.e., progressive infection. This is because their post-treatment outcomes are predicated on the latent reservoir size, which was not different in the PBMCs between the PTCs and CPs in the pVISCONTI study [15,16]. Accordingly, the fits of these models were substantially poorer, as shown by the VPC plots (Fig. S14, S18), than fits of our model (Fig. S8).

### Supplementary figures

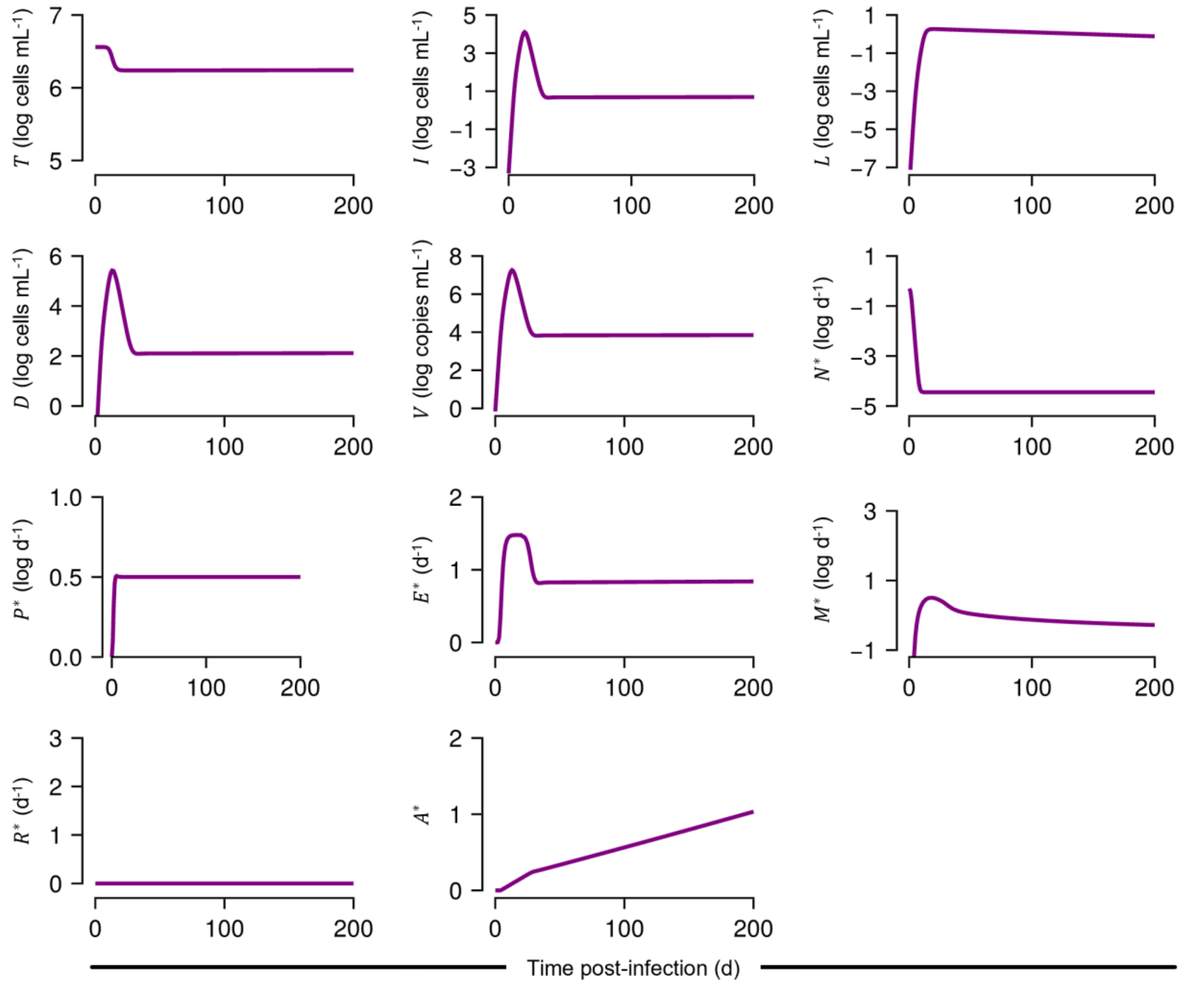


**Fig. S1. Untreated infection simulated by the model.**

Predictions of the model equations (16)–(25) from the main text for a representative set of parameter values (Table S1).

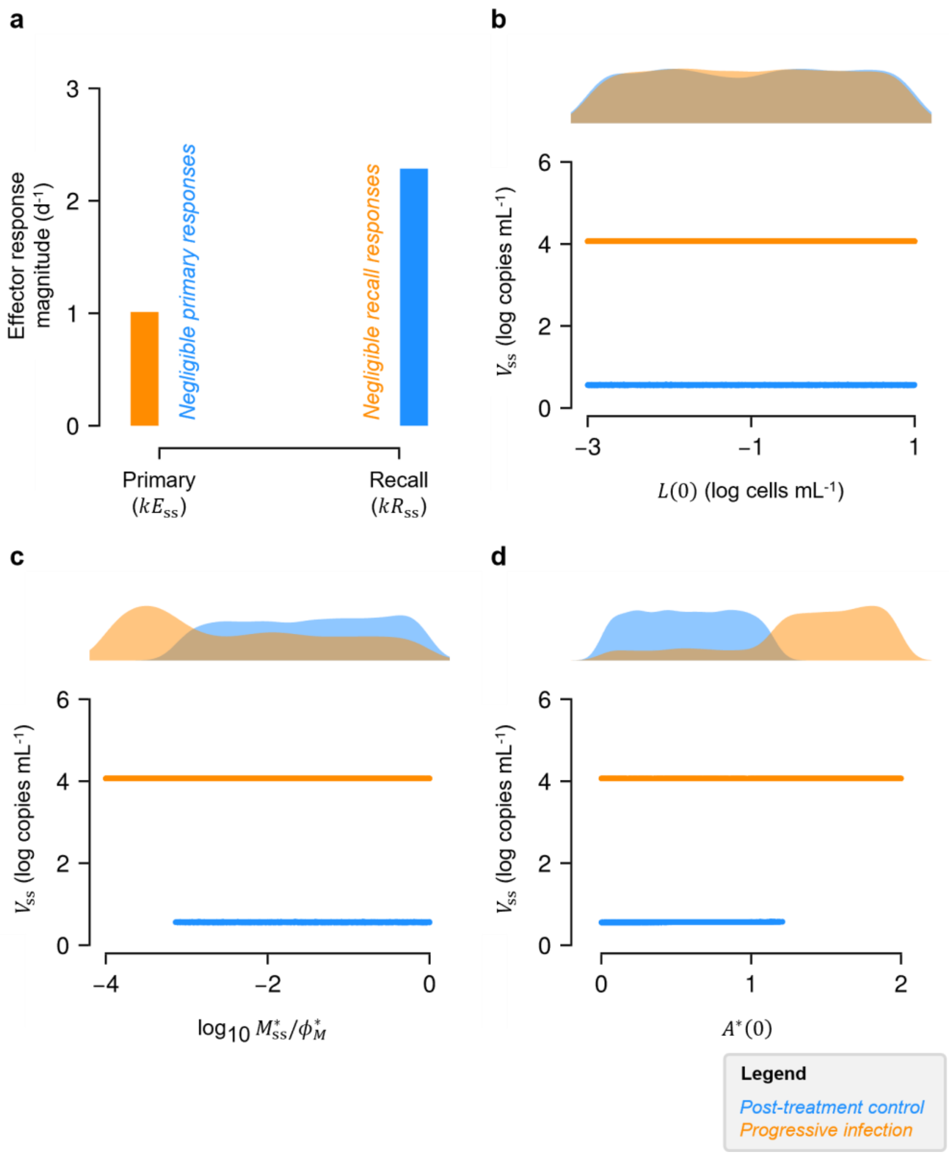


**Fig. S2. Post-treatment outcomes for a range of initial conditions at treatment interruption.**

(**a**) Steady state naïve-derived primary and memory-derived recall responses predicted for PTCs and CPs. Distributions of **(b)** latent reservoir size, **(c)** memory pool size, and **(d)** cumulative antigenic stimulation level at treatment interruption plotted against the post-treatment outcomes.

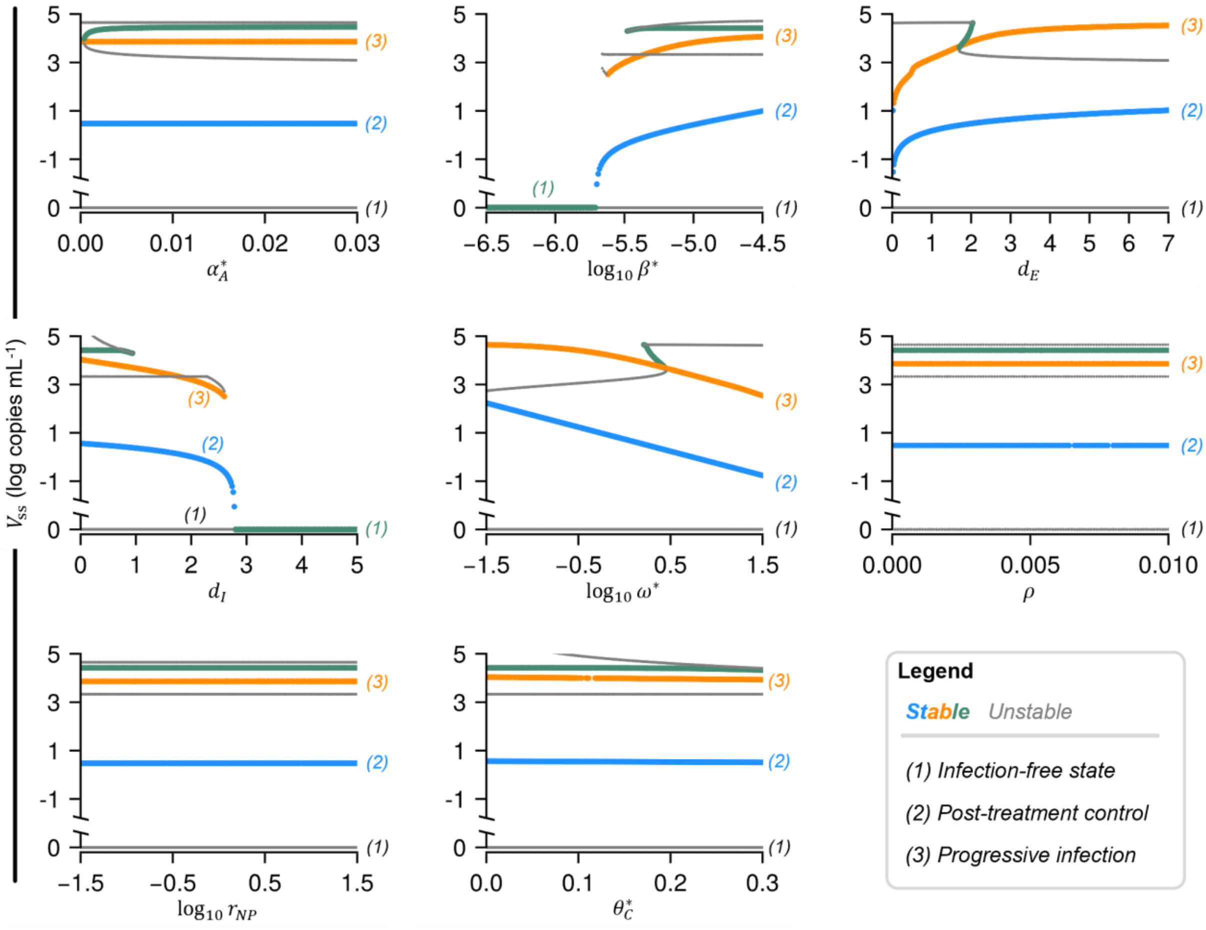


**Fig. S3. Linear stability analysis of the model.**

Steady state viral load (denoted with the subscript ss ) of the model estimated by solving it over ranges of key model parameters (Text S4, Methods in the main text).

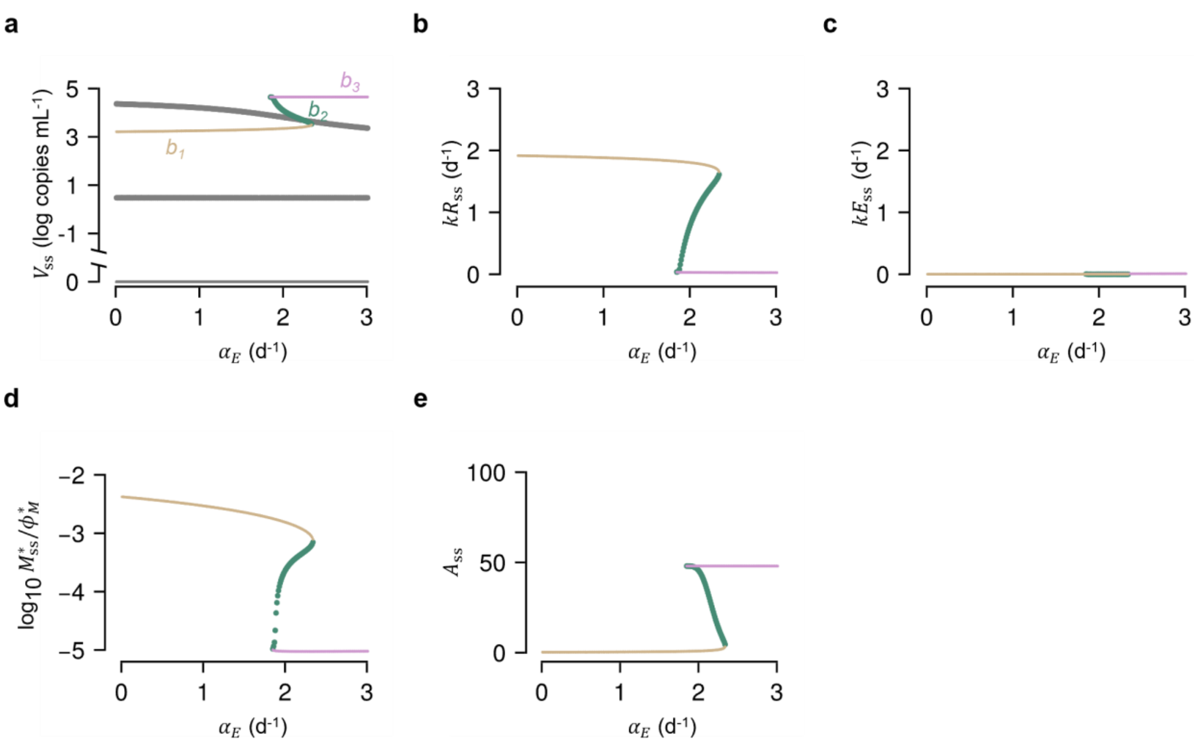


**Fig. S4. Bifurcation diagram indicating the third stable steady state.**

Steady state **(a)** viral load, **(b)** memory-derived effector response level, **(c)** naïve-derived effector response level, **(d)** memory pool size scaled by its carrying capacity $\phi_M^*$ , and **(e)** cumulative antigenic stimulation level of the model estimated by solving it for different values of per-capita proliferation rate of effector cells, $\alpha_E$ (Fig. 4a, Text S4, Methods in the main text). Thin branches $b_1$ (brown) and $b_3$ (pink) are unstable; thick branch $b_2$ (green) is stable.

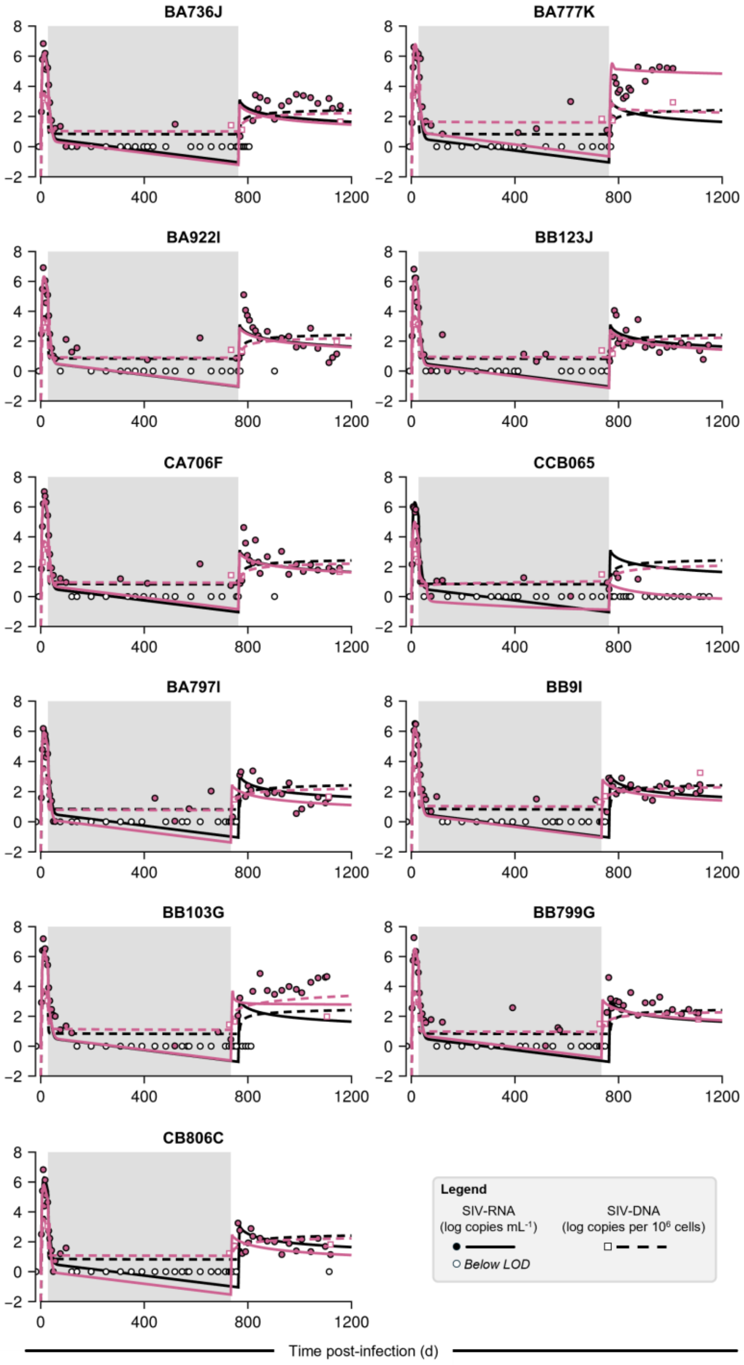


**Fig. S5. Fits of our model to data of early-treated macaques.**

Our model fits to the longitudinal viral load (SIV-RNA) and total proviral reservoir size (SIV-DNA) for all early-treated macaques. Colored circles are viral load measurements and empty circles are observations below the limit of detection ($10^0$ copies $mL^{-1}$); colored squares are reservoir size measurements[15]. The grey-shaded area indicates ART. Individual parameter estimates are listed in Table S4. Black curves indicate population parameter predictions (Table 1).

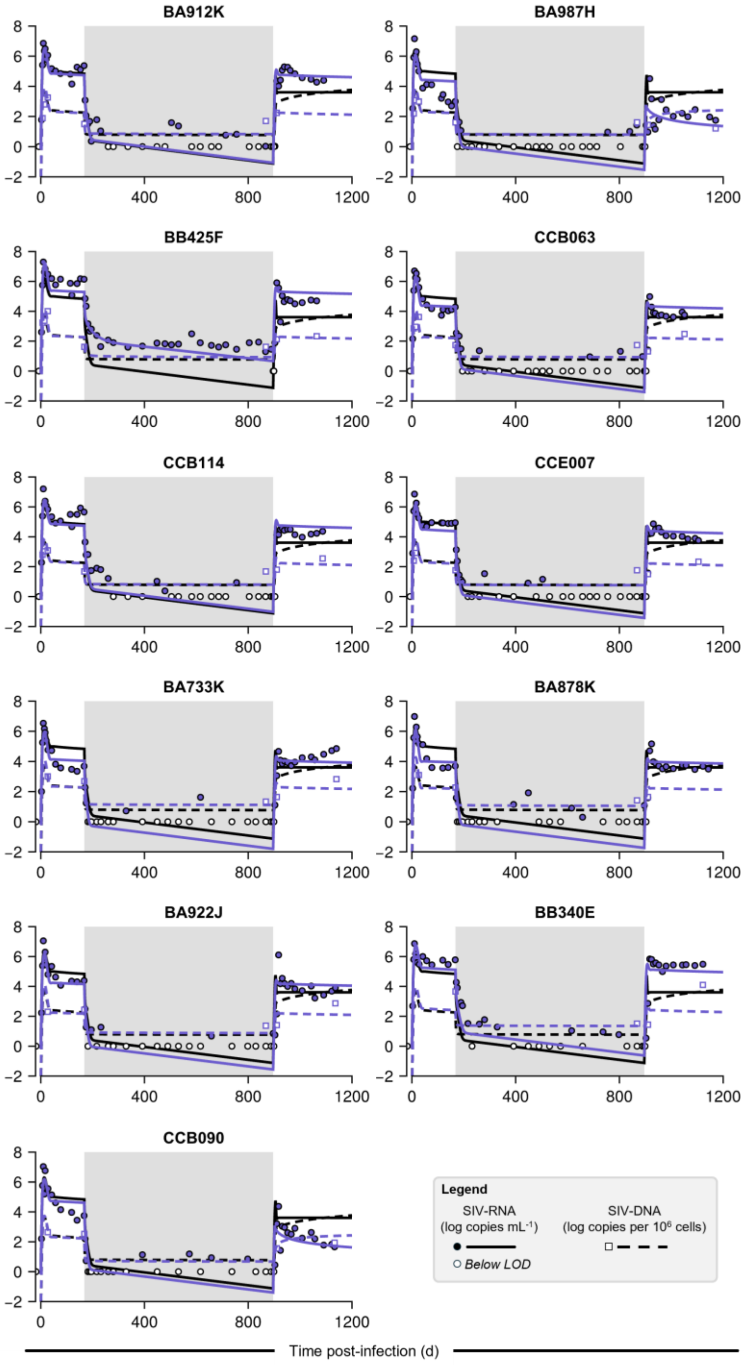


**Fig. S6. Fits of our model to data of late-treated macaques.**

Our model fits to the longitudinal viral load (SIV-RNA) and total proviral reservoir size (SIV-DNA) for all late-treated macaques. Colored circles are viral load measurements and empty circles are observations below the limit of detection ($10^0$ copies $mL^{-1}$); colored squares are reservoir size measurements[15]. The grey-shaded area indicates ART. Individual parameter estimates are listed in Table S4. Black curves indicate population parameter predictions (Table 1).

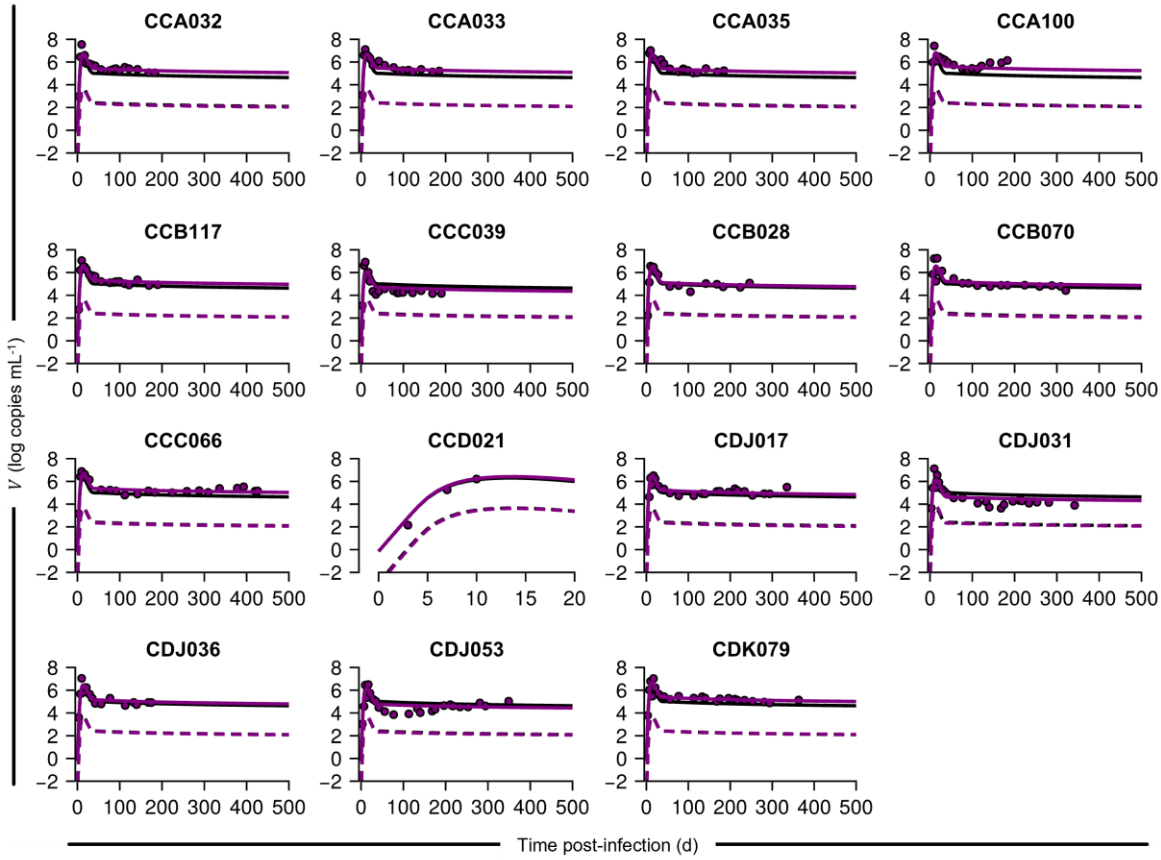


**Fig. S7. Fits of our model to data of untreated macaques.**

Our model fits to the longitudinal viral load (SIV-RNA) for all early-treated macaques. Colored circles are viral load measurements[15]. Individual parameter estimates are listed in Table S4. Black curves indicate population parameter predictions (Table 1).

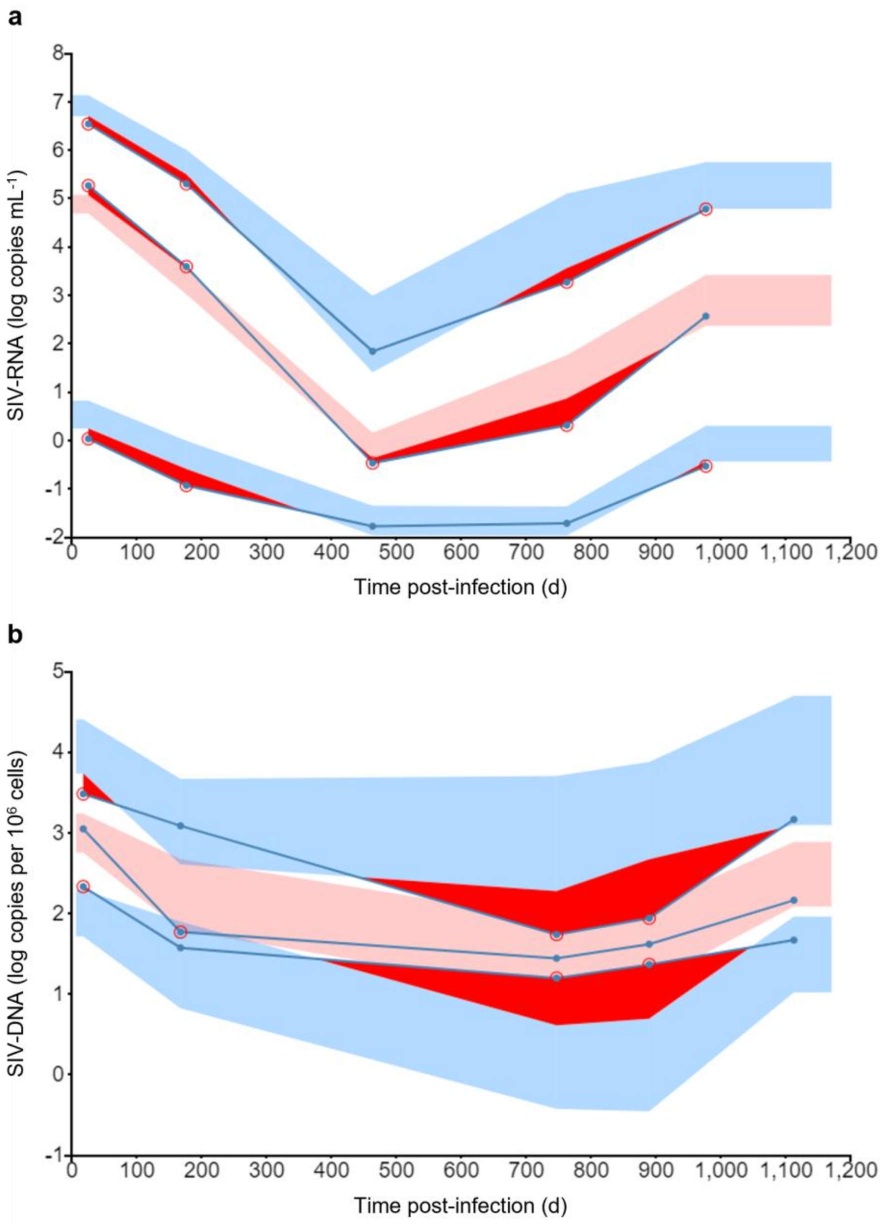


**Fig. S8. Visual predictive check (VPC) plots for our model fits.**

Colored bands indicate 90th percentile (top blue), median (orange), and 10th percentile (bottom blue) prediction intervals, while the dots connected by lines indicate the 90th percentile (top), median (middle), and 10th percentile (bottom) values of the data.

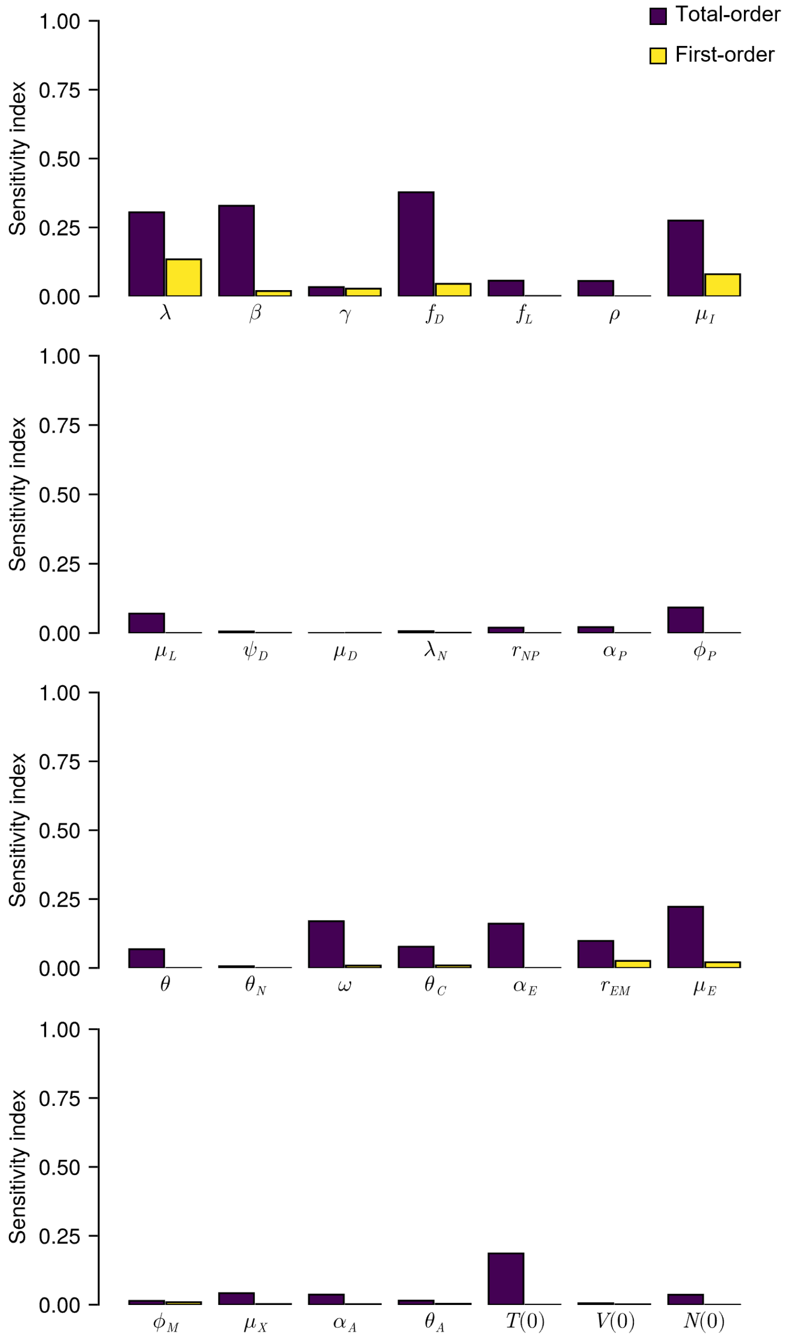


**Fig. S9. Sensitivity analysis for untreated macaques.**

Sensitivity of the set-point viral load to model parameter values quantified using eFAST method.

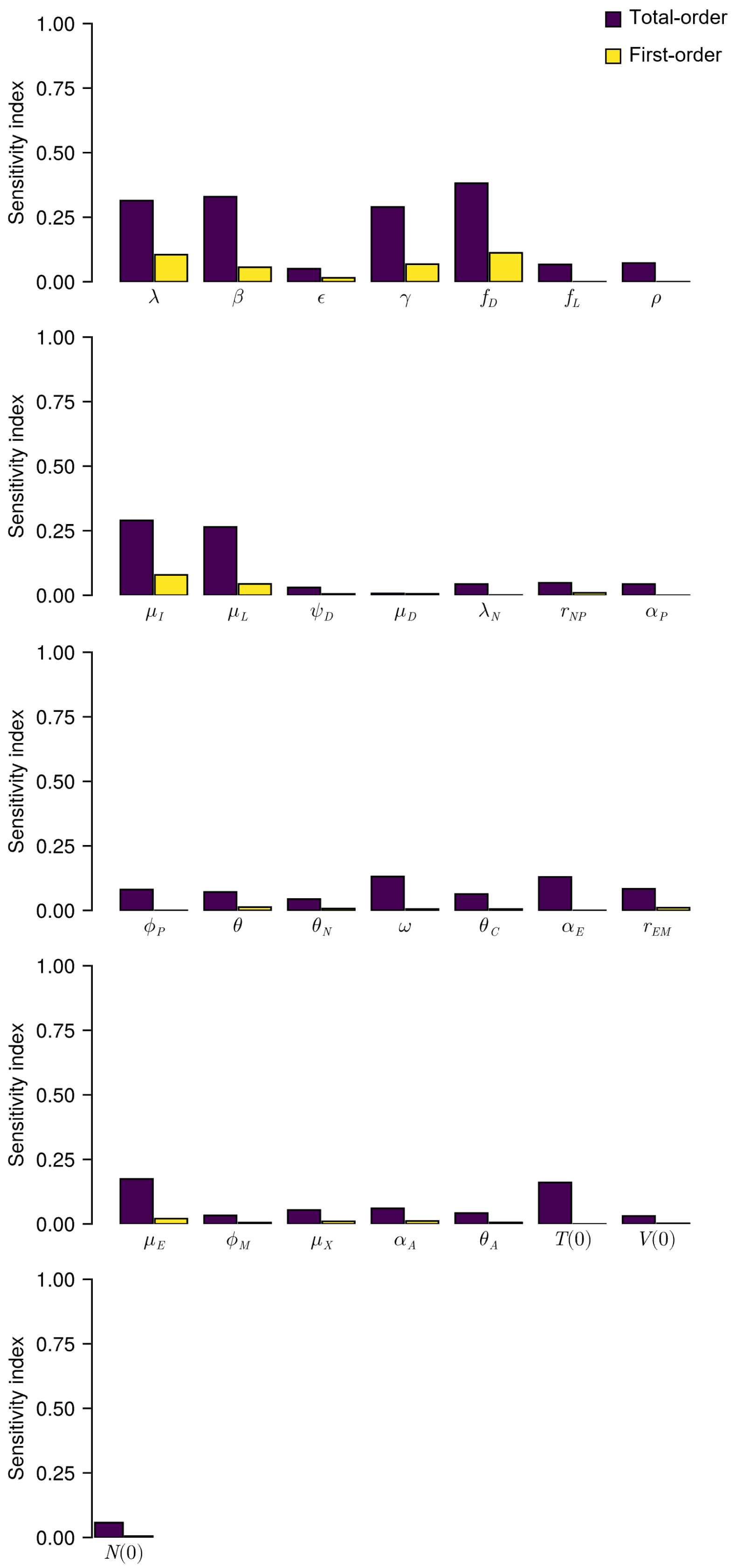


**Fig. S10. Sensitivity analysis for treated macaques.**

Sensitivity of the set-point viral load to model parameter values quantified using eFAST method.

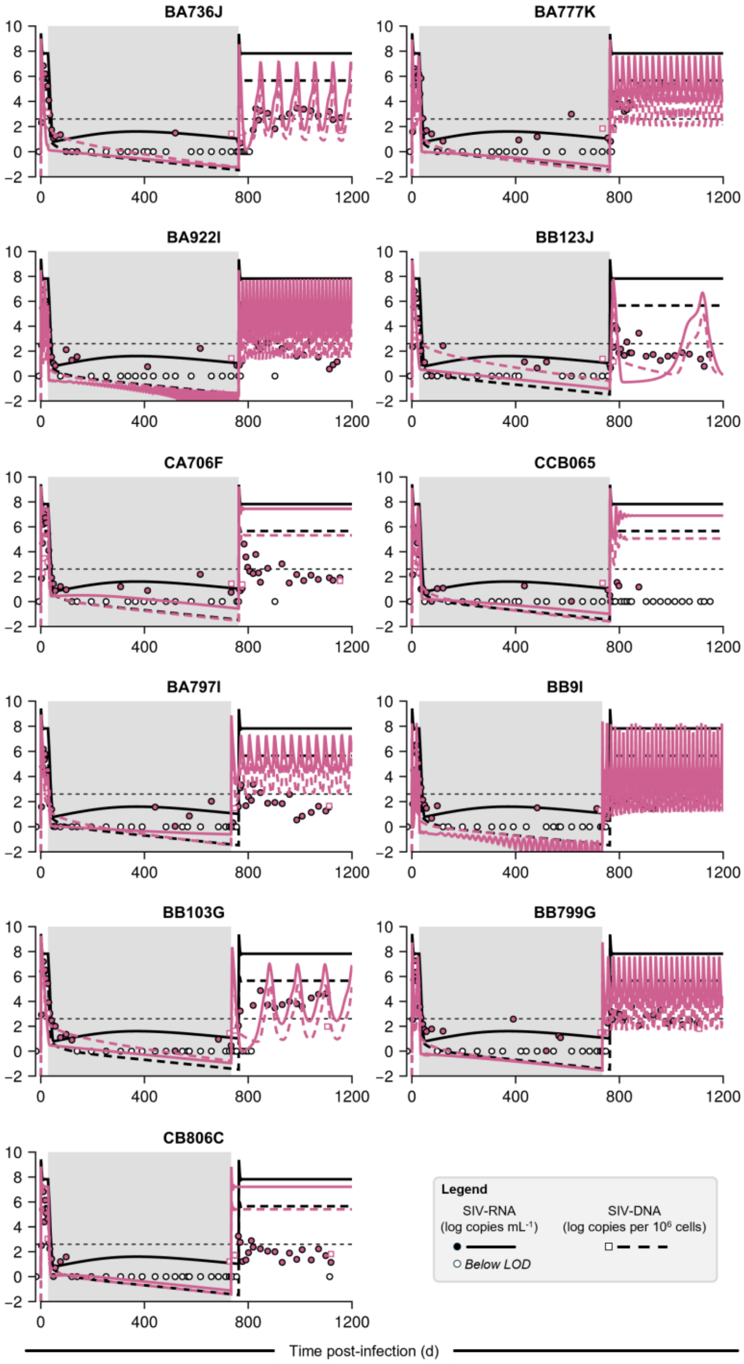


**Fig. S11. Fits of model #1 to data of early-treated macaques.**

Fits of model #1 to the longitudinal viral load (SIV-RNA) and total proviral reservoir size (SIV-DNA) for all early-treated macaques. Colored circles are viral load measurements and empty circles are observations below the limit of detection ($10^0$ copies mL$^{-1}$); colored squares are reservoir size measurements[15]. The grey-shaded area indicates ART. Black curves indicate population parameter predictions (Table S5).

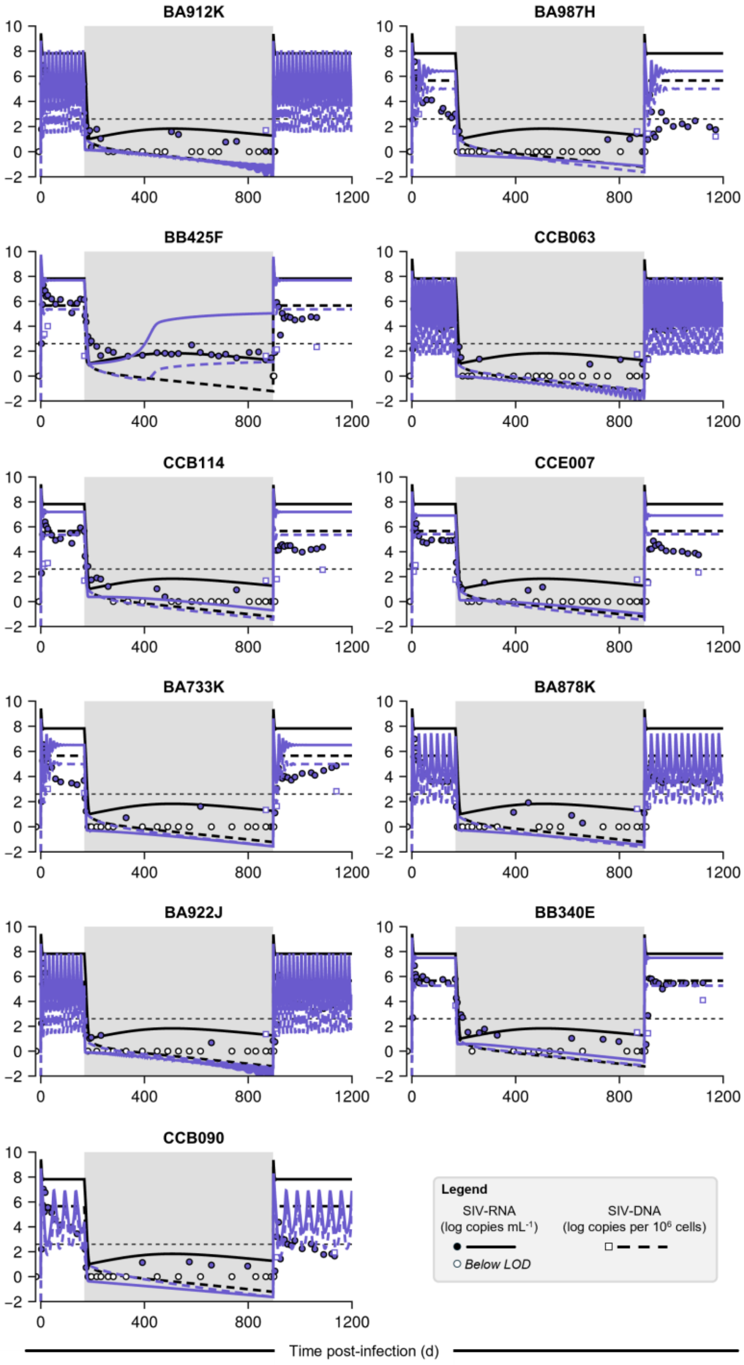


**Fig. S12. Fits of model #1 to data of late-treated macaques.**

Fits of model #1 to the longitudinal viral load (SIV-RNA) and total proviral reservoir size (SIV-DNA) for all late-treated macaques. Colored circles are viral load measurements and empty circles are observations below the limit of detection ($10^0$ copies mL$^{-1}$); colored squares are reservoir size measurements[15]. The grey-shaded area indicates ART. Black curves indicate population parameter predictions (Table S5).

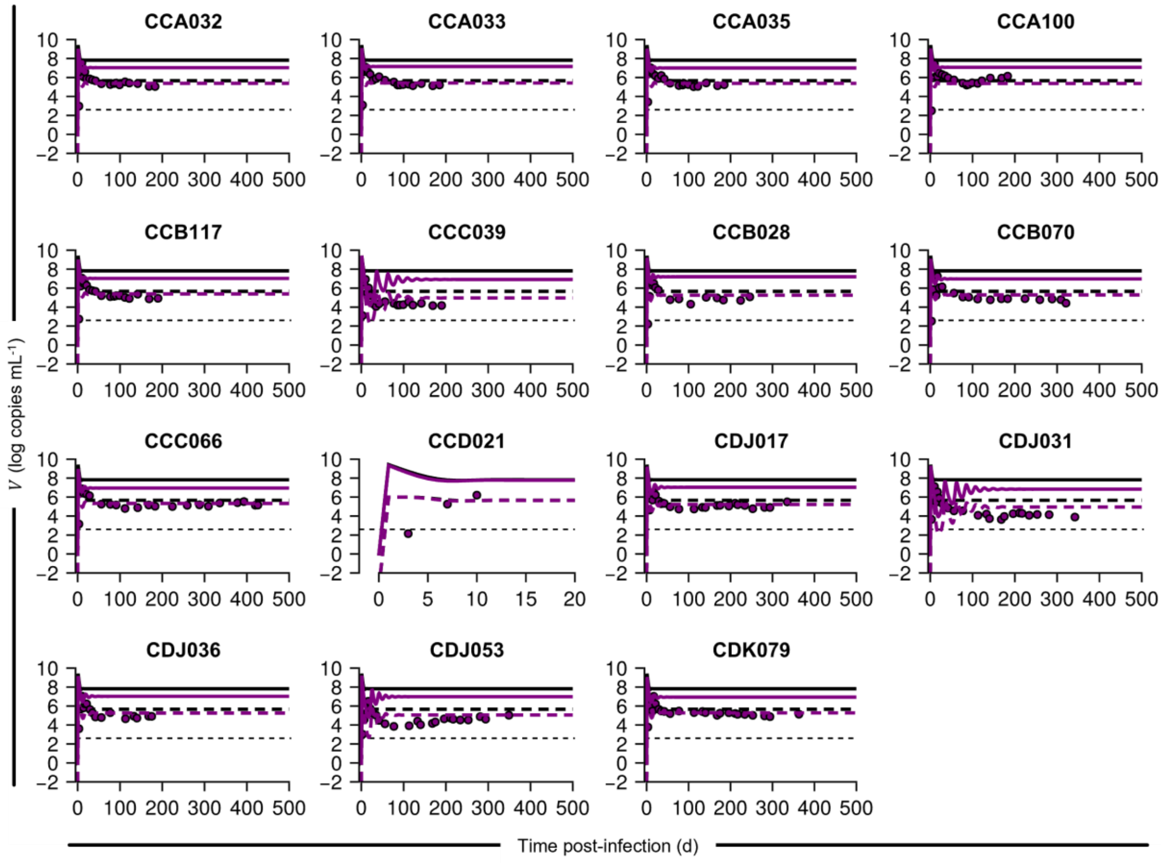


**Fig. S13. Fits of model #1 to data of untreated macaques.**

Fits of model #1 to the longitudinal viral load (SIV-RNA) for all early-treated macaques. Colored circles are viral load measurements[15]. Individual parameter estimates are listed in Table S4. Black curves indicate population parameter predictions (Table S5).

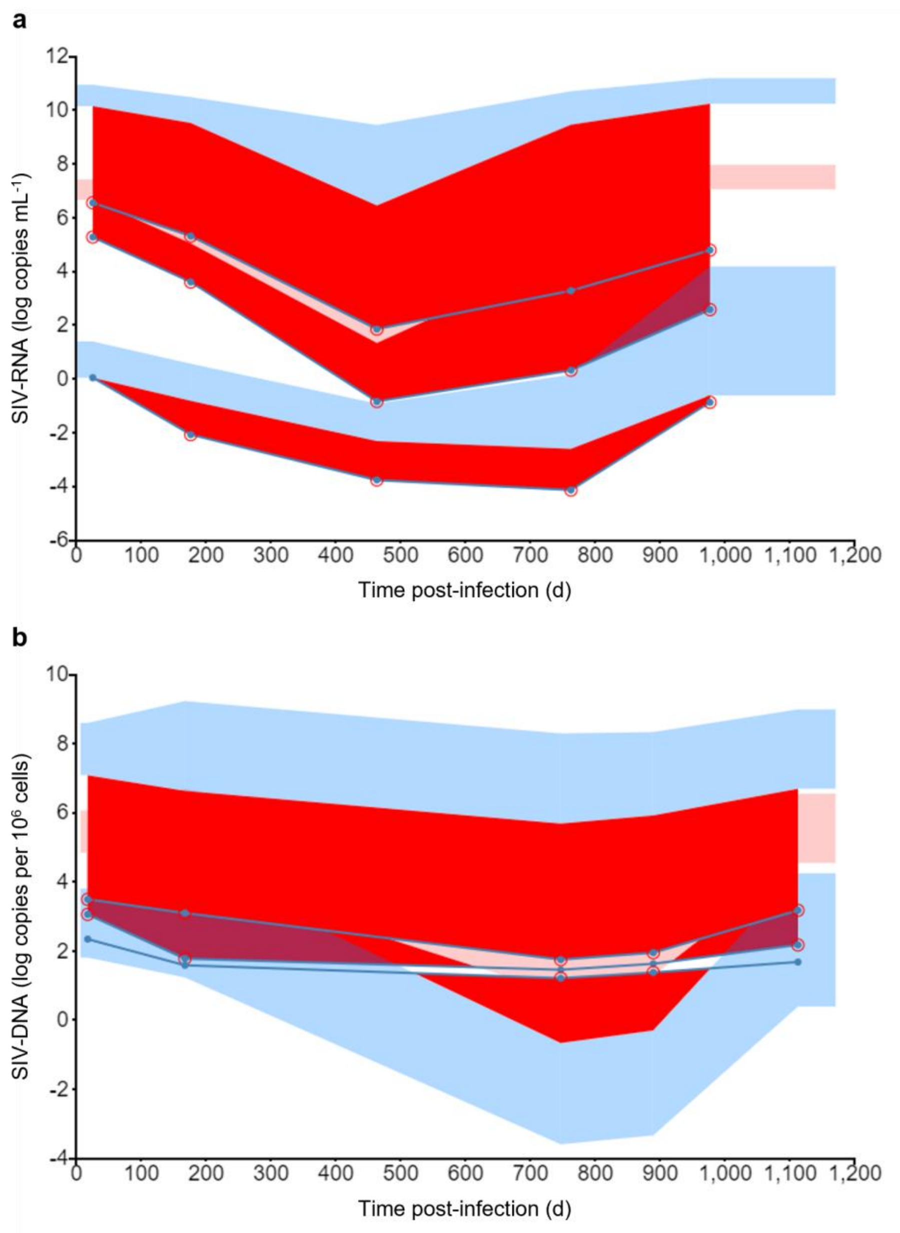


**Fig. S14. Visual predictive check (VPC) plots for fits of model #1.**

Colored bands indicate 90th percentile (top blue), median (orange), and 10th percentile (bottom blue) prediction intervals, while the dots connected by lines indicate the 90th percentile (top), median (middle), and 10th percentile (bottom) values of the data.

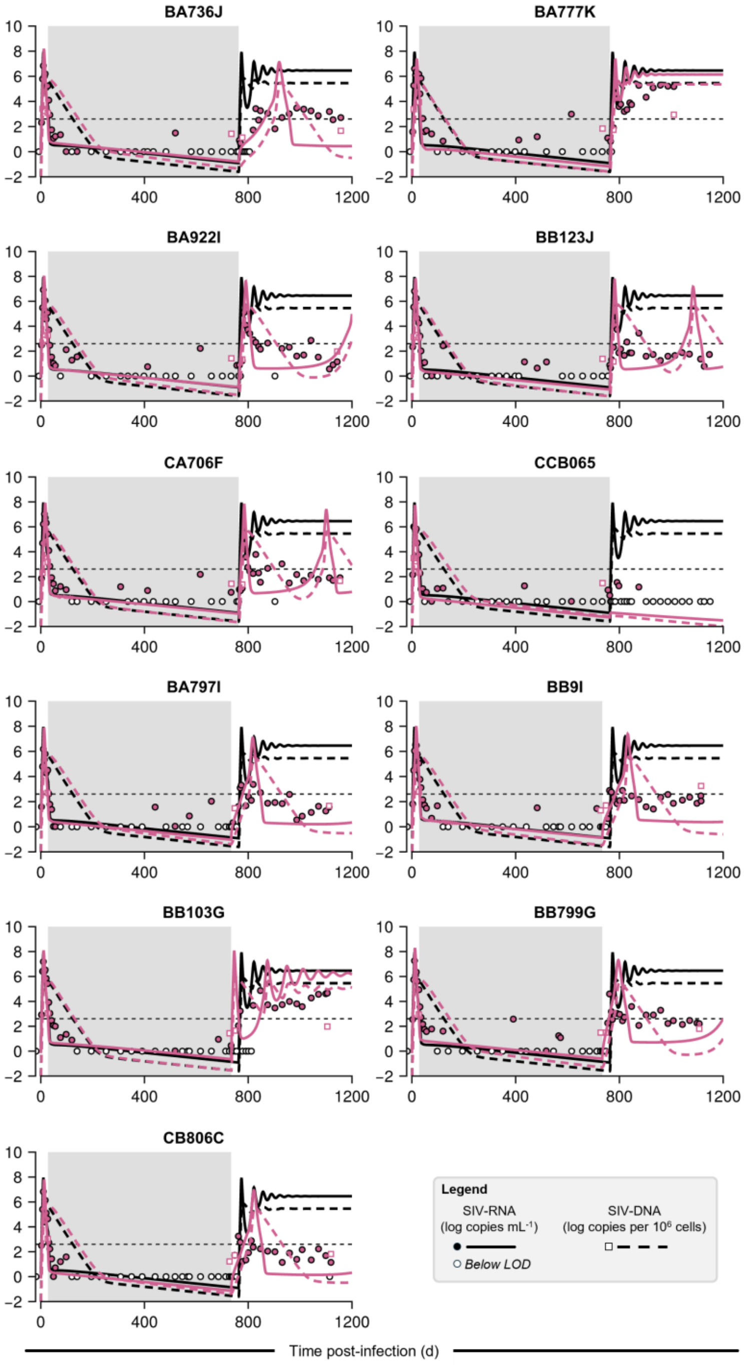


**Fig. S15. Fits of model #2 to data of early-treated macaques.**

Fits of model #2 to the longitudinal viral load (SIV-RNA) and total proviral reservoir size (SIV-DNA) for all early-treated macaques. Colored circles are viral load measurements and empty circles are observations below the limit of detection ($10^0$ copies $mL^{-1}$); colored squares are reservoir size measurements[15]. The grey-shaded area indicates ART. Black curves indicate population parameter predictions (Table S6).

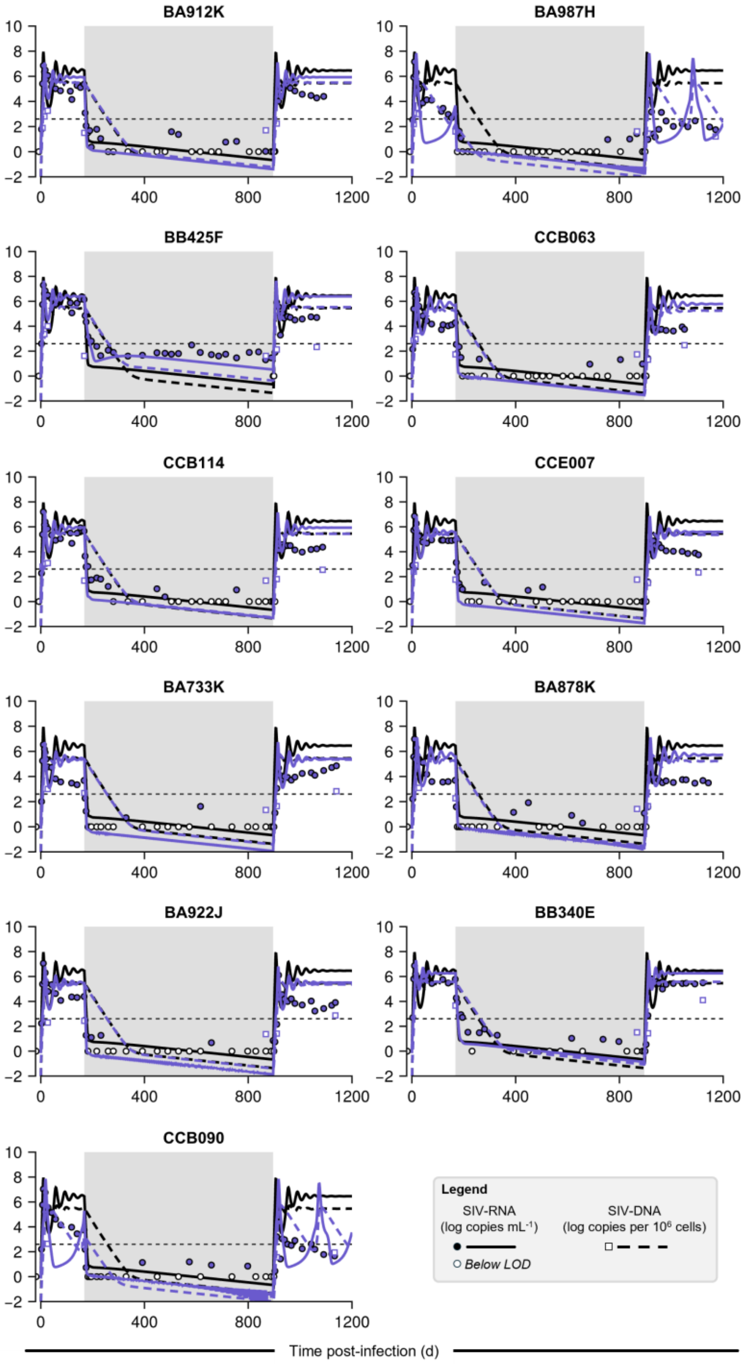


**Fig. S16. Fits of model #2 to data of late-treated macaques.**

Fits of model #2 to the longitudinal viral load (SIV-RNA) and total proviral reservoir size (SIV-DNA) for all late-treated macaques. Colored circles are viral load measurements and empty circles are observations below the limit of detection ($10^0$ copies $mL^{-1}$); colored squares are reservoir size measurements[15]. The grey-shaded area indicates ART. Black curves indicate population parameter predictions (Table S6).

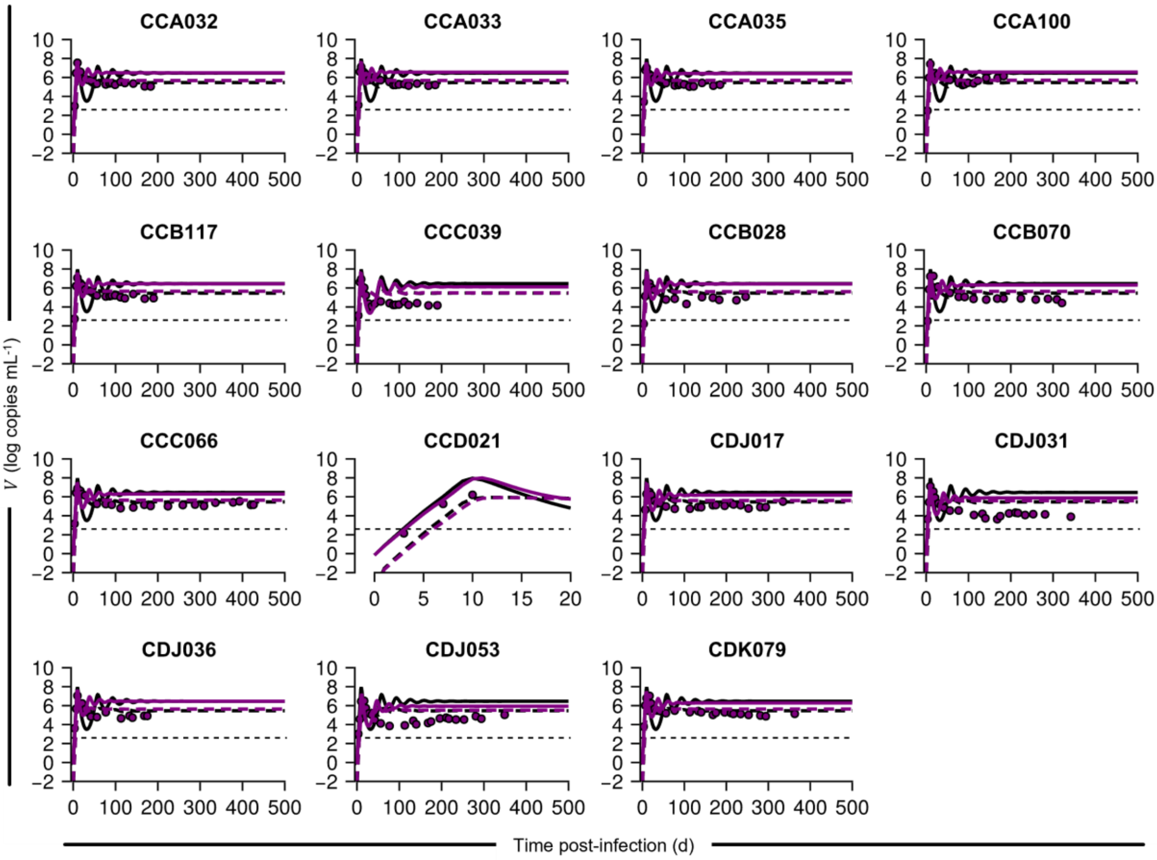


**Fig. S17. Fits of model #2 to data of untreated macaques.**

Fits of model #2 to the longitudinal viral load (SIV-RNA) for all early-treated macaques. Colored circles are viral load measurements[15]. Individual parameter estimates are listed in Table S4. Black curves indicate population parameter predictions (Table S6).

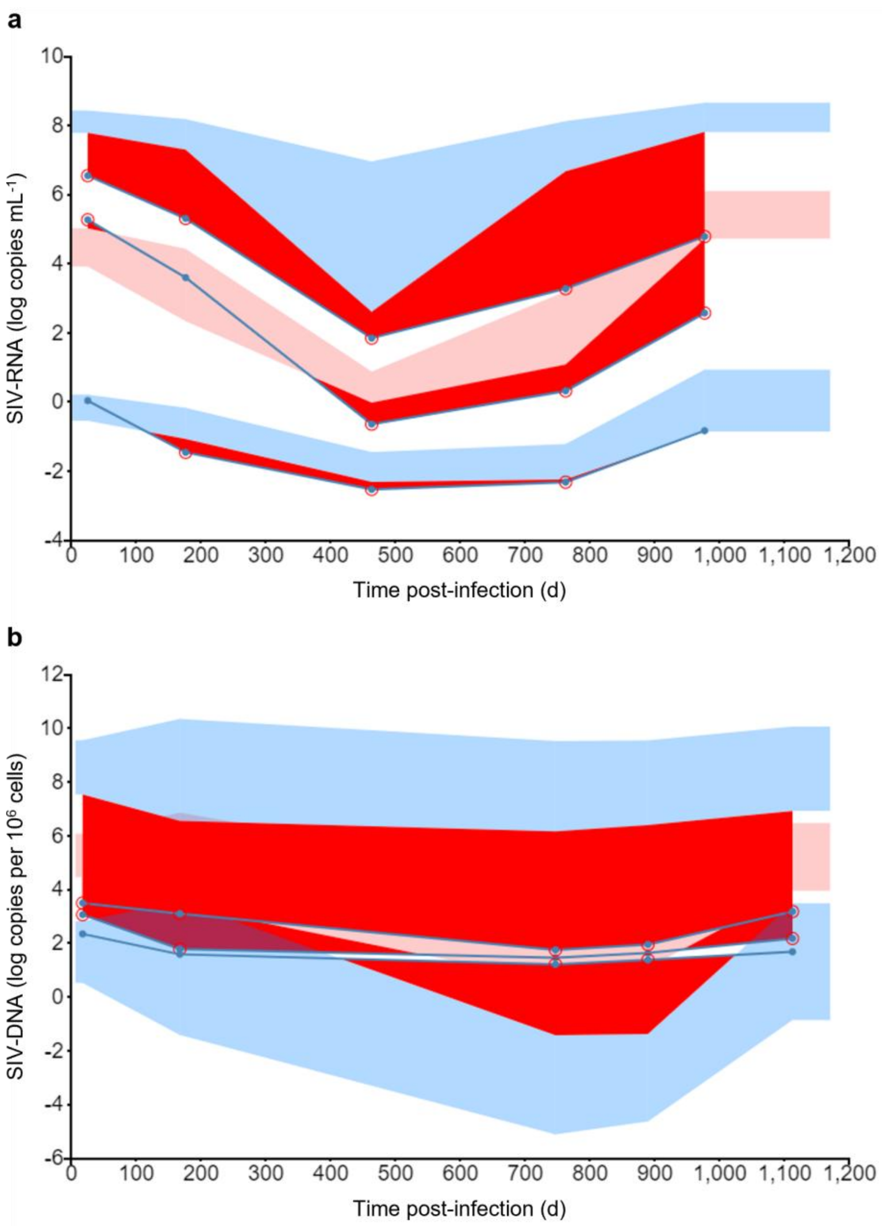


**Fig. S18. Visual predictive check (VPC) plots for fits of model #2.**

Colored bands indicate 90th percentile (top blue), median (orange), and 10th percentile (bottom blue) prediction intervals, while the dots connected by lines indicate the 90th percentile (top), median (middle), and 10th percentile (bottom) values of the data.

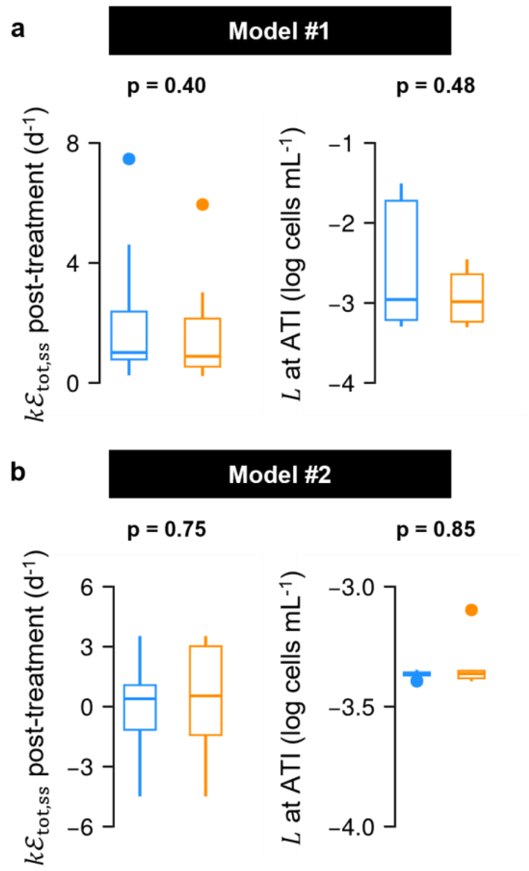


**Fig. S19. Comparison of predictions of models #1 and #2.**

Predictions of post-treatment steady state effector responses (**a, c**) and latent reservoir sizes at treatment interruption (**b, d**) compared for model #1 (**a, b**) and model #2 (**c, d**) between PTCs (blue) and CPs (orange). Statistical comparison was done using Mann-Whitney U test.

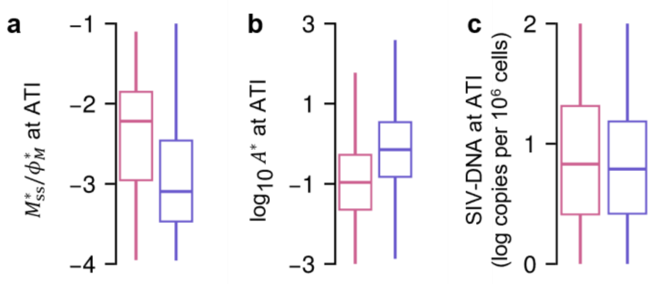


**Fig. S20. Comparing memory pool and SIV-DNA proviral reservoir sizes at treatment interruption for the virtual population between treatment groups.**

**(a)** Memory CD8 T cell pool size, **(b)** cumulative antigenic stimulation levels, and **(c)** SIV-DNA levels at ART interruption compared between the early-treated (pink) and late-treated (slate blue) groups.

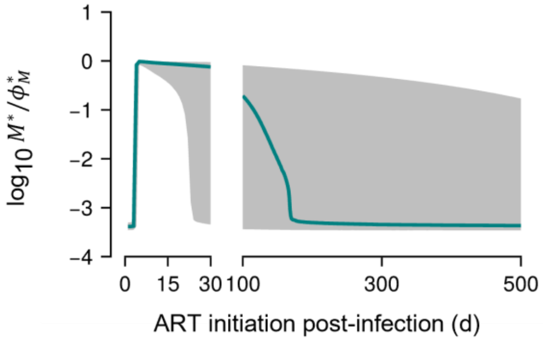


**Fig. S21. Steady state memory pool size post-treatment with different ART initiation times.**

Memory pool size, $M^*$, is scaled by its carrying capacity, $\phi_M^*$. The green line indicates the median value, the grey band is the inter-quartile range. Estimations were obtained using population parameter estimates for our model (Table 1, Fig. 6).

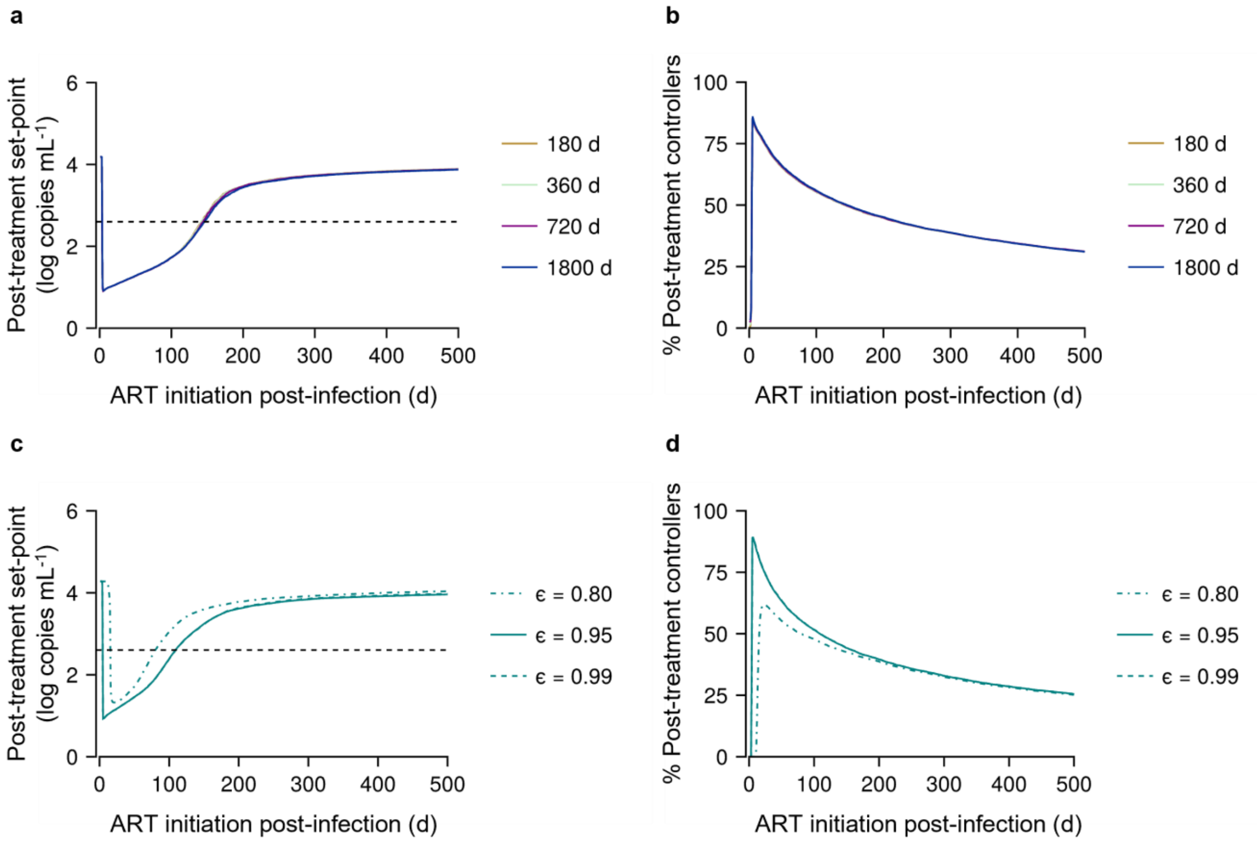


**Fig. S22. Window of opportunity for different ART administration settings.**

**(a, c)** Post-ART set-point viral load and **(b, d)** percentage of PTCs predicted in the virtual population for **(a, b)** different ART durations and **(c, d)** efficacies. The dashed line is the threshold value of 400 copies mL[-1] distinguishing PTCs from CPs. Note that in **(a)** and **(b)**, all curves overlap, while in **(c)** and **(d)**, curves for $\epsilon$ values 0.95 and 0.99 overlap.

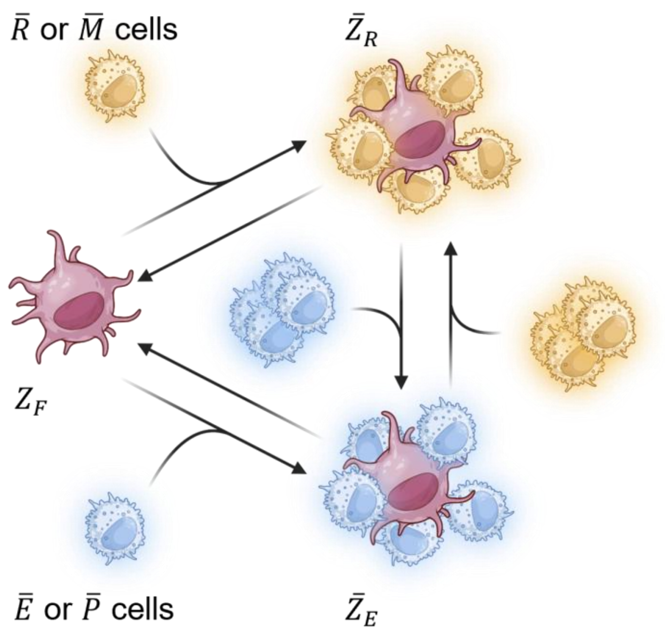


**Fig. S23. Schematic for the different states of antigen-bearing APCs.**

Antigen-bearing APCs (brown) are in one of the three states: engaged with primary ($\bar{P}$ or $\bar{E}$ cells), engaged with recall ($\bar{M}$ or $\bar{R}$ cells) subtype, or free from any engagement. Their levels are denoted $\bar{Z}_E$, $\bar{Z}_R$, and $Z_F$, respectively. Processes governing the transition of an APC between these states are encoded in equations (S9)–(S11), followed by their description.

### Supplementary tables

**Table S1. Representative parameter values of an individual.**

Parameter values of the model used for calculations in Fig. 2, 3e, 3f, S1, and S2. These values correspond to those of macaque #9 from Table S4 with $\alpha_E$ and $\alpha_A^*$ tuned slightly.

| **Parameter** | **Value** | **Parameter** | **Value** |
|---|---|---|---|
| $\lambda$ (cells/mL-d) | 385.29 | $\theta_N$ (cells mL$^{-1}$) | 0.01 |
| $\log_{10}\beta^*$ (log mL cells$^{-1}$ d$^{-1}$) | -4.96 | $\log_{10}\omega$ (log d$^{-1}$) | 0.26 |
| $\gamma$ (copes cells$^{-1}$) | 713.48 | $\theta_C^*$ (cells mL$^{-1}$) | 0.01 |
| $\epsilon$ | 0.88 | $\alpha_E$ (d$^{-1}$) | 1.7 |
| $f_D$ | 0.93 | $\log_{10} r_{EM} \left(=\log_{10} r_{RM}\right)$ (d$^{-1}$) | -0.5 |
| $f_L$ | 10$^{-6}$ | $d_E \left(=d_R\right)$ (d$^{-1}$) | 2.0 |
| $\rho$ (d$^{-1}$) | 0.001 | $\phi_M^*$ (d$^{-1}$) | 1000.0 |
| $d_I$ (d$^{-1}$) | 0.5 | $d_X$ (d$^{-1}$) | 1.2 |
| $d_L$ (d$^{-1}$) | 0.004 | $\alpha_A^*$ (d$^{-1}$) | 0.01 |
| $\psi_D$ | 0.81 | $\theta_A^*$ | 5.0 |
| $d_D$ (d$^{-1}$) | 10$^{-5}$ | $m$ | 4 |
| $\log_{10}\lambda_N^*$ (log d$^{-2}$) | -4.3 | $n$ | 4 |
| $\log_{10} r_{NP}$ (log d$^{-1}$) | 0.15 | $\log_{10} T(0)$ (log cells mL$^{-1}$) | 6.56 |
| $\alpha_P$ (d$^{-1}$) | 0.63 | $\log_{10} V(0)$ (log copies mL$^{-1}$) | -0.459 |
| $\phi_P^*$ (d$^{-1}$) | 0.5 | $\log_{10} N^*(0)$ (log d$^{-1}$) | -0.301 |

| Parameter | Value | Parameter | Value |
|---|---|---|---|
| $\theta$ (cells mL$^{-1}$) | 1.0 | $\log_{10} P^*(0)$ (log d$^{-1}$) | -3.301 |

**Table S2. Initial conditions for simulations in Fig. 2.**

5,000 samples were drawn from each of these distributions, with each triplet of $L$, $M^*$, and $A^*$ representing one individual condition.

| Variable | Distribution of the initial condition |
|---|---|
| $\log_{10} L$ | Uniform(-3.0, 1.0) |
| $\log_{10} M^*$ | Uniform(-1.0, 3.0) |
| $A^*$ | Uniform(0.0, 2.0) |

---

**Table S3. Fixed parameters in our model and the initial conditions.**

All the parameters listed in the table are fixed to a value obtained from the literature, where possible, except for $\lambda$, which used reported values for both fixed and random effects[12]. Thus, the parameter is different between individuals. Note that although $P^*(0)=0$, we provide a small non-zero starting value ($N^*(0)/1000$) to prevent divergence of the model estimates as $P^*$ is in the denominator of the rate equation $dA^*/dt$.

| Parameter | Value | Reference |
|---|---|---|
| *Fixed parameters* | | |
| $\lambda$ (cells mL$^{-1}$ d$^{-1}$) | Fixed effect = 352.7<br>Random effect = 1.32 | 12 |
| $f_D$ | 0.93 | 12 |
| $\rho$ (d$^{-1}$) | 0.001 | 13 |
| $d_I$ (d$^{-1}$) | 0.5 | 17,18 |
| $d_L$ (d$^{-1}$) | 0.004 | 13 |
| $d_D$ (d$^{-1}$) | 0.069 | 12 |
| $\log_{10} \lambda_N^*$ (log d$^{-2}$) | -4.3 | 8 |
| $\log_{10} r_{NP}$ (log d$^{-1}$) | 0.15 | Assumed to be faster than the differentiation of effectors to memory cells, $r_{EM} (= r_{RM})$, as the initial recruitment of naïve cells must be rapid |
| $\alpha_P$ (d$^{-1}$) | 0.63 | 12 |
| $\phi_P^*$ (d$^{-1}$) | 0.5 | Equal to the initial naïve pool size that gets activated in response to infection |

| Parameter | Value | Reference |
|---|---|---|
| $\theta$ (cells mL$^{-1}$) | 1.0 | [13] |
| $\theta_N$ (cells mL$^{-1}$) | 0.01 | Assumed; required $<\theta$ for quick activation of naïve cells |
| $\theta_C^*$ (cells mL$^{-1}$) | 0.01 | Assumed; required $=\theta_N$ to ensure the same number of APCs that trigger the initial response later continue providing antigenic stimulation |
| $\log_{10} r_{EM} \left(=\log_{10} r_{RM}\right)$ | -0.5 | Assumed; ~5% of effector cells become memory cells [19] |
| $d_E \left(=d_R\right)$ (d$^{-1}$) | 2.0 | [13] |
| $\phi_M^*$ (d$^{-1}$) | 1000.0 | Assumed; required $>\phi_P^*$ |
| $d_X$ (d$^{-1}$) | 1.2 | Assumed $\sim d_E$; highest death rates among CD8 T cell subtypes are for effectors [20], but the half-life of memory cells in a chronic infection is similar to that of terminally differentiated cells |
| $\theta_A^*$ | 5.0 | Assumed $\sim\theta$ since effectors dominate the CD8 T cell pool |
| $m$ | 4 | [14] |
| $n$ | 4 | [14] |
| *Initial conditions* | | |
| $\log_{10} V(0)$ (log copies mL$^{-1}$) | -0.46 | [15] |

| Parameter | Value | Reference |
|---|---|---|
| $\log_{10} N^{*}(0)$ (log $d^{-1}$) | -0.30 | [8] |

**Table S4. Parameter estimates of individual macaques for our model.**

For the parameter $\lambda$, we used both fixed and random effects from a previous study[12] (Table S3), and thus, upon fitting, the parameter was different between macaques. Random effects for $\beta^*$, $\alpha_E$, and $T(0)$ were removed as they were low (Table 1 in the main text), rendering them to be identical between macaques. Fixed parameters are listed in Table S3.

| **Macaque** | $\lambda$ | $\gamma$ | $\epsilon$ | $\psi_D$ | $\alpha_E$ | $\alpha_A^*$ |
|---|---|---|---|---|---|---|
| *Early-treated macaques* | | | | | | |
| 1 (BA736J) | 334.19 | 1131.23 | 0.90 | 0.74 | 1.93 | 0.00081 |
| 2 (BA777K) | 199.00 | 595.48 | 0.85 | 0.59 | 1.85 | 0.05721 |
| 3 (BA922I) | 334.16 | 1350.36 | 0.89 | 0.80 | 1.92 | 0.00208 |
| 4 (BB123J) | 448.71 | 1002.17 | 0.90 | 0.80 | 1.92 | 0.00255 |
| 5 (CA706F) | 361.69 | 1624.04 | 0.90 | 0.80 | 1.91 | 0.00175 |
| 6 (CCB065) | 407.26 | 31.15 | 0.78 | 1.05 | 1.90 | 0.00037 |
| 7 (BA797I) | 341.23 | 573.15 | 0.89 | 0.96 | 1.92 | 0.00090 |
| 8 (BB9I) | 309.60 | 934.25 | 0.88 | 0.75 | 1.92 | 0.00294 |
| 9 (BB103G) | 384.84 | 596.88 | 0.87 | 0.76 | 1.90 | 0.01541 |
| 10 (BB799G) | 334.05 | 2035.39 | 0.90 | 0.75 | 1.92 | 0.00233 |
| 11 (CB806C) | 379.75 | 482.03 | 0.91 | 0.77 | 1.93 | 0.00171 |
| *Late-treated macaques* | | | | | | |
| 12 (BA912K) | 181.31 | 1065.03 | 0.89 | 0.89 | 1.89 | 0.03139 |
| 13 (BA987H) | 309.16 | 356.83 | 0.90 | 0.86 | 1.89 | 0.00095 |
| 14 (BB425F) | 169.96 | 3112.36 | 0.78 | 0.76 | 1.89 | 0.01333 |
| 15 (CCB063) | 161.55 | 358.21 | 0.89 | 0.83 | 1.86 | 0.02240 |
| 16 (CCB114) | 158.22 | 940.93 | 0.87 | 0.86 | 1.89 | 0.03233 |

| **Macaque** | $\lambda$ | $\gamma$ | $\epsilon$ | $\psi_D$ | $\alpha_E$ | $\alpha_A^*$ |
|---|---|---|---|---|---|---|
| 17 (CCE007) | 138.50 | 423.11 | 0.90 | 0.88 | 1.88 | 0.04925 |
| 18 (BA733K) | 220.56 | 218.27 | 0.92 | 0.77 | 1.87 | 0.03974 |
| 19 (BA878K) | 156.49 | 230.41 | 0.92 | 0.81 | 1.85 | 0.03438 |
| 20 (BA979I) | 186.38 | 307.63 | 0.90 | 0.86 | 1.87 | 0.03808 |
| 21 (BB340E) | 206.26 | 2232.36 | 0.86 | 0.60 | 1.90 | 0.04114 |
| 22 (CCB090) | 325.05 | 649.50 | 0.93 | 0.90 | 1.90 | 0.00089 |
| *Untreated macaques* | | | | | | |
| 23 (CCA032) | 354.15 | 2582.37 | 0.90 | 0.91 | 1.91 | 0.00500 |
| 24 (CCA033) | 353.21 | 2870.42 | 0.90 | 0.91 | 1.91 | 0.00519 |
| 25 (CCA035) | 354.19 | 2614.28 | 0.90 | 0.91 | 1.91 | 0.00535 |
| 26 (CCA100) | 359.64 | 3781.01 | 0.90 | 0.91 | 1.91 | 0.00491 |
| 27 (CCB117) | 351.92 | 2118.53 | 0.90 | 0.91 | 1.91 | 0.00517 |
| 28 (CCC039) | 341.61 | 500.66 | 0.90 | 0.87 | 1.91 | 0.00475 |
| 29 (CCB028) | 352.86 | 1249.49 | 0.90 | 0.91 | 1.90 | 0.00493 |
| 30 (CCB070) | 348.59 | 1552.66 | 0.90 | 0.91 | 1.91 | 0.00506 |
| 31 (CCC066) | 362.02 | 2464.93 | 0.90 | 0.89 | 1.91 | 0.00491 |
| 32 (CCD021) | 352.64 | 1141.28 | 0.90 | 0.91 | 1.91 | 0.00482 |
| 33 (CDJ017) | 361.22 | 1584.62 | 0.90 | 0.91 | 1.91 | 0.00495 |
| 34 (CDJ031) | 311.07 | 540.58 | 0.90 | 0.95 | 1.90 | 0.00573 |
| 35 (CDJ036) | 352.04 | 1360.84 | 0.90 | 0.91 | 1.91 | 0.00496 |
| 36(CDJ053) | 360.17 | 631.85 | 0.90 | 0.94 | 1.90 | 0.00480 |
| 37 (CDK079) | 356.49 | 2267.53 | 0.90 | 0.91 | 1.91 | 0.00491 |

**Table S5. Population parameter estimates for model #1.**

Values for fixed parameters were obtained from the literature[12,13]: $\lambda$ ~ Normal(352.7, 1.32) cells mL$^{-1}$ d$^{-1}$, $f_L$ = 10$^{-6}$, $\rho$ = 10$^{-3}$ d$^{-1}$, $d_I$ = 0.5 d$^{-1}$, $d_L$ = 0.004 d$^{-1}$, $\log_{10}\lambda_E^*$ ~ Normal(0.0, 0.25) cells mL$^{-1}$ d$^{-1}$, $\theta_E$ = 0.1 cells mL$^{-1}$, $\alpha_X$ = 2.0 d$^{-1}$, $\theta_X$ = 5.0 cells mL$^{-1}$, $d_E$ = 2.0 d$^{-1}$. Also, $\log_{10}V(0)$ = 0.459 copies mL$^{-1}$. All other variables start from zero. RSE: Relative standard error.

| **Parameter** | **Fixed effect (%RSE)** | **Random effect (%RSE)** |
|---|---|---|
| $\log_{10}\beta^*$ (log mL cells$^{-1}$ d$^{-1}$) | -2.71 (1.18) | - |
| $\gamma$ (copies cells$^{-1}$) | 8.63×10$^4$ (15.9) | 0.72 (16.9) |
| $\epsilon$ | 0.98 (0.61) | 0.46 (115) |
| $\alpha_E$ (d$^{-1}$) | 1.44 (2.59) | - |
| $\log_{10}T(0)$ (log cells mL$^{-1}$) | 4.36 (0.91) | 0.04 (23.3) |

**Table S6. Population parameter estimates for model #2.**

Values for fixed parameters were obtained from the literature[12,13]: $\lambda$ ~ Normal(352.7, 1.32) cells mL$^{-1}$ d$^{-1}$, $f_L$ = $10^{-6}$, $f_D$ = 0.93, $\rho$ = $10^{-3}$ d$^{-1}$, $d_I$ = 0.5 d$^{-1}$, $d_L$ = 0.004 d$^{-1}$, $d_D$ = 0.069 d$^{-1}$, $\log_{10}\lambda_E^*$ ~ Normal(0.0, 0.25) cells mL$^{-1}$ d$^{-1}$, $\theta_E$ = 0.1 cells mL$^{-1}$, $\xi$ = 3.0 d$^{-1}$, $q_c$ = 2.0 cells mL$^{-1}$, $w$ = 3, $d_E$ = 2.0 d$^{-1}$, $\kappa$ = 1.0 d$^{-1}$, $d_Q$ = 0.1 d$^{-1}$. Also, $\log_{10}V(0)$ = 0.459 copies mL$^{-1}$. All other variables start from zero. RSE: Relative standard error.

| Parameter | Fixed effect (%RSE) | Random effect (%RSE) |
|---|---|---|
| $\log_{10}\beta^*$ (log mL cells$^{-1}$ d$^{-1}$) | -2.62 (0.32) | 0.03 (29.1) |
| $\gamma$ (copies cells$^{-1}$) | 11.63×10$^4$ (17.2) | 0.73 (16.7) |
| $\epsilon$ | 0.83 (31.8) | 1.11 (158) |
| $\alpha_E$ (d$^{-1}$) | 1.45 (2.26) | 0.06 (85.5) |
| $\log_{10}T(0)$ (log cells mL$^{-1}$) | 4.33 (0.08) | - |

**Table S7. Comparison of fit statistics of different models.**

| Statistic | Our model | Model #1 | Model #2 |
|---|---|---|---|
| -2×Log likelihood | 4011.03 | 6250.33 | 5273.03 |
| Bayesian information criterion (BIC) | 4032.69 | 6264.78 | 5287.47 |
| SIV-RNA fit error | 1.01 | 2.23 | 1.47 |
| SIV-DNA fit error | 0.66 | 1.77 | 2.42 |
